# Subtle introgression footprints at the end of the speciation continuum in a clade of *Heliconius* butterflies

**DOI:** 10.1101/2022.12.19.520581

**Authors:** Quentin Rougemont, Bárbara Huber, Simon Martin, Annabel Whibley, Catalina Estrada, Darha Solano, Robert Orpet, W. Owen McMillan, Brigitte Frérot, Mathieu Joron

## Abstract

Quantifying gene flow between lineages at different stages of the speciation continuum is central to understanding speciation. *Heliconius* butterflies have undergone an adaptive radiation in wing colour patterns driven partly by natural selection for local mimicry. Colour patterns are also known to be used as assortative mating cues. Therefore, wing pattern divergence is considered to play a role in speciation. A corollary is that mimicry between closely-related species may be associated with hybridization and interfere with reproductive isolation. Here, we take a multifaceted approach to explore speciation history, species boundaries, and traits involved in species differentiation between the two closely-related species *H. hecale* and *H. ismenius.* We focus on geographic regions where the two species mimic each other, and contrast this with geographic regions where they do not mimic each other. To examine population history and patterns of gene flow, we tested and compared a four-population model accounting for linked selection. This model suggests that the two species have remained isolated for a large part of their history, yet with a small amount of gene exchange. Accordingly, signatures of genomic introgression were small except at a major wing pattern allele and chemosensing genes, and stronger in the mimetic populations compared to non-mimetic populations. Behavioural assays confirm that visual confusion exists but that short-range cues determine strong sexual isolation. Tests for chemical differentiation between species identified major differences in putative pheromones which likely mediate mate choice and the maintenance of species differences.

## Introduction

Understanding the modalities of species formation and the maintenance of species barriers is central to evolutionary biology. Toward this goal, an understanding of the demographic history and quantifying gene flow and isolation among populations, is fundamental. Recent empirical population genomics studies have documented the near ubiquity of the heterogeneous landscape of differentiation across a continuum of increasing divergence (Ravinet et al. 2017). These landscapes can arise under divergence hitchhiking, where continuous gene flow will impede genetic differentiation outside of areas involved in local adaptation (Via and West 2008; Feder et al. 2012; Flaxman et al. 2013). Unfortunately, similar patterns can arise under models of secondary contact, where gene flow erodes past genetic differentiation in regions without barrier loci (Barton and Bengtsson 1986). Another complicating factor is the effect of selection at linked sites, where hitchhiking of neutral alleles linked to a selective sweep can also contribute to a heterogeneous landscape (Noor and Bennett 2009; Cruickshank and Hahn 2014). Local variation in recombination and in the density of selected sites within the region determines the intensity of linked selection (Kaplan et al. 1989; Nordborg et al. 1996; Payseur and Nachman 2002; Burri 2017). Linked selection reduces polymorphism at sites harbouring positive as well as deleterious variants and at surrounding sites, especially in areas of low recombination (Hill and Robertson 1966; Charlesworth et al. 1993). Linked selection can be modelled as a local reduction in effective population size (*Ne*), although this is a simplified approach. A modelling approach that jointly allows for local genomic variation in effective population size and migration rate can improve our understanding of the demographic processes at play during population divergence.

*Heliconius* butterflies constitute a clade composed of numerous hybridising species, providing an excellent system to understand the contribution of gene flow, linked selection and demography to the genomic landscape of divergence (Martin et al. 2019; Van Belleghem et al. 2020). Many colour pattern morphs within a species are common across the radiation with, for instance, up to 7 sympatric forms within a single population of *H. numata* and over 25 distinct wing pattern populations in the *H. melpomene* or *H. erato* lineages (Mallet and Gilbert 1995). *Heliconius* butterflies are unpalatable to predators, advertise their toxicity through their colour pattern and are classic examples of Müllerian mimicry, where distasteful species converge to a common warning signal (Sheppard et al. 1985). The loci responsible for colour pattern variation have been identified using a combination of genetic mapping, GWAS, gene expression analyses and more recently functional knockouts. These studies have revealed that a few highly conserved genes interact to modulate much of the wing pattern variation both within and between *Heliconius* species. This includes *i)* the gene *optix,* which controls the distribution of red-orange pattern elements (Reed et al. 2011; Zhang et al. 2017; Huber et al. 2015; Lewis et al. 2019), *ii)* the gene *cortex,* which controls the presence and position of white and yellow pattern elements (Nadeau et al. 2016; Livraghi et al. 2021), *iii)* the gene *wntA,* which controls melanic patterning across the wing (Martin et al. 2012; Moest et al. 2020; Fenner et al. 2020; Van Belleghem et al. 2020) and *iv) aristaless 1,* which controls a white/yellow switch (reviewed in McMillan et al. 2020).

Here, we focus on a pair of closely-related species, *H. hecale* and *H. ismenius*, which belong to the so-called silvaniform lineage and have a complex history of wing pattern mimicry and diversification (Huber et al. 2015). In Eastern Panama, *H. hecale melicerta* and *H. ismenius boulleti* are perfect co-mimics of each other and both display a pattern made of orange (proximal) and yellow elements (distal), bordered by a thick black margin and a black wing tip. In Western Panama, *H. hecale zuleika* and *H. ismenius clarescens* resemble each other but participate in different local mimicry rings. *H. h. zuleika* shows a pattern composed of a largely black forewing with yellow dots and an orange hindwing, and mimics *H. hecalesia formosus, Eueides procula* and various highly distasteful butterflies in the tribe Ithomiini, notably *Tithorea tarricina*. In contrast, *H. i. clarescens* has largely orange wings striped with black and yellow and does not seem to belong to any obvious mimicry ring (Rosser et al. 2015). Previous studies have demonstrated that *Heliconius* co-mimics share the same habitats and that details in the colour patterns matter in butterflies species recognition (Mallet and Gilbert 1995). However, chemical signalling has been suggested to also be involved in maintaining species integrity between pairs of sympatric co-mimetic species where colour-based recognition may be compromised (Mérot et al. 2015, González-Rojas et al. 2020) including in the species we studied here (Mann et al. 2017). The contrasting situation observed here, *i.e.* perfect mimicry vs. non-mimicry (hereafter considered as co-mimics vs non co-mimics) offers a unique opportunity to study the role of gene flow in the evolution of mimicry. We tested the hypothesis of adaptive introgression of wing patterning genes among co-mimics, and the expectation of local gene flow to be stronger between co-mimics than between non co-mimics. We combined whole genome analyses, as well as behavioural and chemistry assays to understand the modalities of gene flow among species. We aimed at i) reconstructing the divergence history of *H. hecale* and *H. ismenius,* taking into account barriers to gene flow and linked selection in a four-population model that includes powerful statistics for gene flow detection, ii) quantifying the extent of fine scale variation in gene flow along the genome and its role in the maintenance of co-mimicry and convergence in wing pattern among *H. hecale melicerta* and *H. ismenius boulleti,* and iii) evaluating the role of sex, behaviour and chemical factors in the maintenance of the strong isolation observed along the genome of this species complex.

## Results

### Strong divergence between species

A total of 73 individuals were sampled and genotyped using whole genome sequencing **(Fig 1A** and **Fig 1B, Table S01)**. These include the two perfect co-mimics *H. hecale melicerta* and *H. ismenius boulleti* from Eastern Panama (n = 10) and their two sympatric but non-mimetic counterparts *H. hecale zuleika* and *H. ismenius clarescens* from Western Panama (n = 9). In addition, *H. hecale felix* (n = 4)*, H. hecale clearei* (n = 4) *and H. ismenius telchinia* (n = 2) were included to test for gene flow and assess global population structure. To further evaluate introgression and population structure, *H. pardalinus* (n = 15) and *H. numata silvana* (n = 17) specimens were included in our analyses as well as *H. melpomene* (n = 12). These species are all closely related outgroups to *H. hecale* and *H. ismenius*.

**Figure 1.**
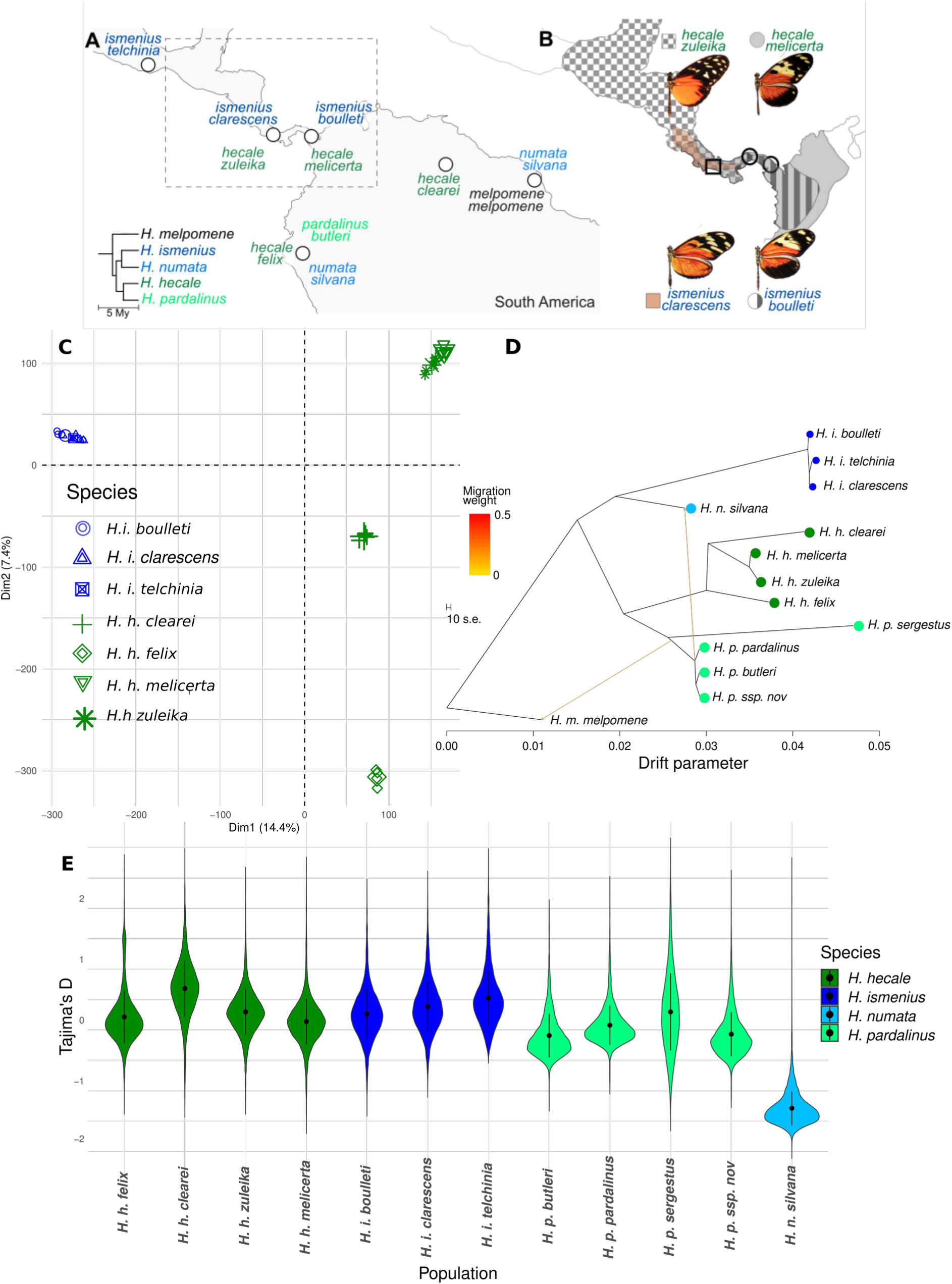
Sampling, population structure and evolutionary relationship among populations. (**A**) Broad scale sampling design and (**B**) Fine scale sampling design as well as phenotypic difference between co-mimics and non co-mimic species. (**C**) Population structure among sampled species and (**D**) evolutionary tree highlighting genetic exchange among populations. (**E**) Tajima’s D in 50 kb windows along the genome. The black dot represents the mean value +/-1 standard deviation.

To estimate the structure of genetic variation among species and populations, we performed a principal component analysis (PCA) on the whole genome data. This revealed an unambiguous separation of the two species, *H. hecale* and *H. ismenius*, along the first axis. The second axis separated individuals of *H. hecale* into three geographic clusters corresponding to 1) *H. hecale felix,* 2) *H. hecale clearei* and 3) a group comprising *H. hecale zuleika* and *H. hecale melicerta* individuals (**Fig 1C**). High levels of net sequence divergence between species (*D_A_* range = 0.16 to 0.22), absolute divergence (*D_XY_* range = 0.15 to 0.22) and differentiation (*F_ST_* range = 0.87 to 0.92) were observed at coding sites and mirrored these observations. Accordingly, measures of relative genetic differentiation (*F_ST_*, **Fig 2A**) and of absolute genetic divergence (*D_XY_*, **Fig 2B, Fig S01** for *D_A_*) across the whole genome were high both between co-mimic and non co-mimic populations.

**Figure 2.**
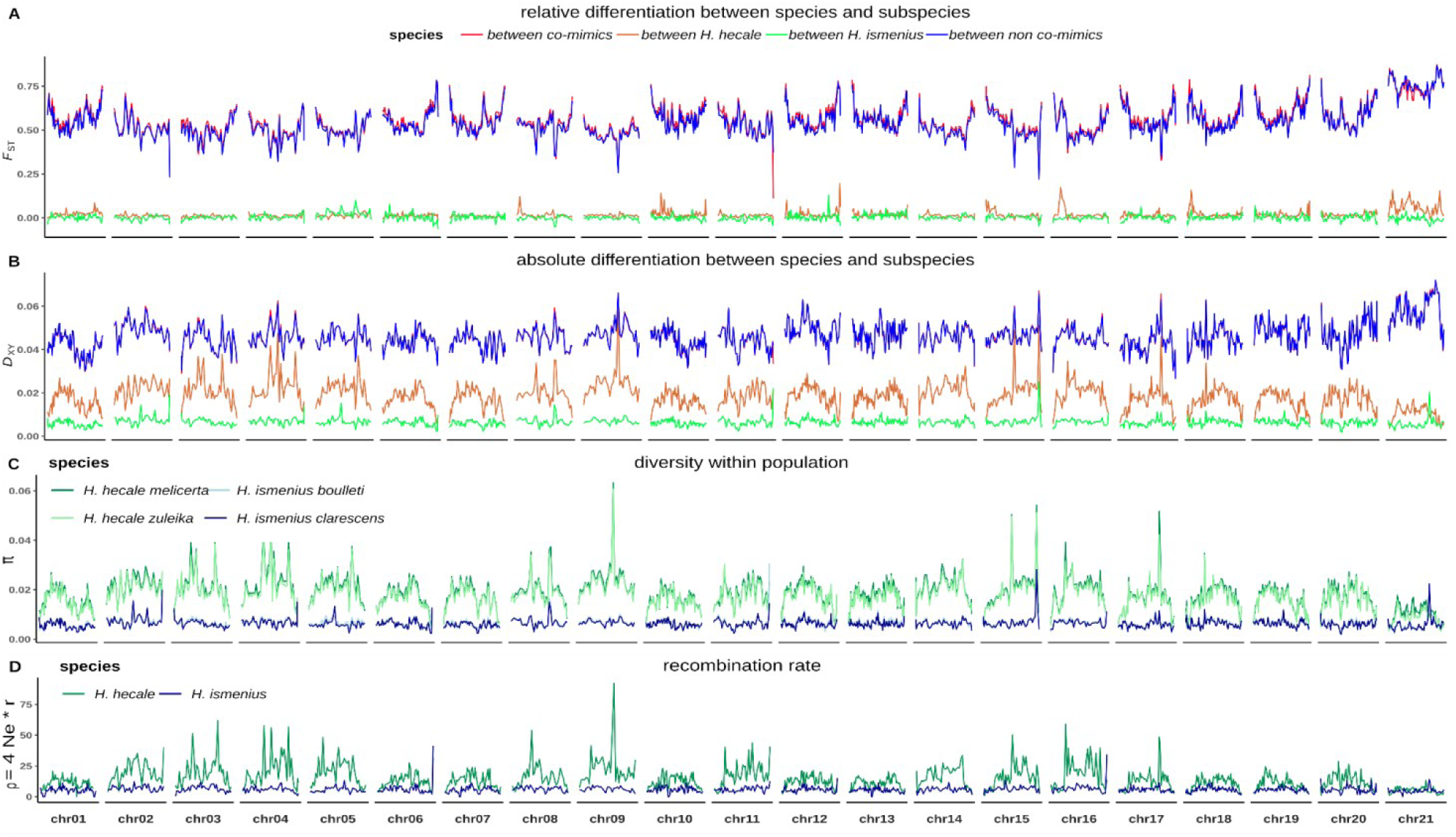
Conserved and high divergence between species. (**A**) Genetic differentiation (*F_ST_*) landscape along each chromosome between sympatric co-mimics (*H. h. melicerta* vs *H. i. boulleti*, red line) and between non co-mimic species (*H. h. zuleika vs H. i. clarescens*, blue line). Genetic differentiation between subspecies of each species is also shown (*H. h. melicerta* vs *H. h. zuleika*, chocolate; *H. i. boulleti* vs *H. i. clarescens*, green). (**B**) Absolute genetic divergence (*D_XY_*) along the genome (see supplementary Figure S01 for *D_A_*). The same colour coding is used. In general, divergence and differentiation landscapes are so conserved between co-mimics and non co-mimics that they overlap. (**C**) Landscape of genetic diversity (π) observed along the genome for each species. Again, landscapes within species are highly conserved. (**D**) Population scaled recombination landscape along the genome for *H*. *hecale* and *H. ismenius*. Computation averaged in 250 kb windows.

These landscapes were conserved across species comparisons, even when including more distant groups of species from the present study (**Fig S02**). Recombination landscapes were conserved among species (**Fig S03**) which likely explained the correlated levels of diversity and divergence at a broad spatial scale as detailed in **Fig S04-S05,** although a full investigation of the role of recombination in shaping divergence landscape was outside the scope of this paper. Genetic differentiation and divergence between subpopulations of *H. hecale* and *H. ismenius* (**Fig 2A** and **Fig 2B**, green and chocolate lines) were lower than between species, as expected under ongoing gene flow, high effective population size, and/or very recent divergence. Interestingly, a few peaks of high differentiation (especially *F_ST_*) and divergence between *H. h. zuleika and H. h. melicerta* were observed in an ocean of low divergence (**Fig 2B**). Accordingly, shared peaks of *F_ST_* and *D_XY_* between *H. h. zuleika and H. h. melicerta* (above the 80% quantile) involved a total of 55 genes spread on 11 chromosomes.

To better assess evolutionary divergence, and test whether the phylogeny followed a simple bifurcating tree or a network with gene flow, we reconstructed the phylogeny using Treemix (Pickrell and Pritchard 2012) and included all outgroups (*H. numata* and *H. pardalinus*, **Fig 1D**). Including migration events only modestly improved the model fit (**Fig S06**). Accordingly, these migration events were inferred to have occurred in the ancient past, from *H. pardalinus* into *H. numata and from H. melpomene into H. pardalinus.* This analysis did not show evidence of gene flow between co-mimics *H. h. melicerta* and *H. i. boulleti*. This network revealed higher genetic drift in *H. ismenius*, a result that translates into increased Tajima’s D values (**Fig 1E**) as compared to *H. hecale* or to other species with a higher effective population size such as *H. pardalinus* or *H. numata* (de Cara et al. 2023) and may be associated with a recent population decline in *H. ismenius*.

### Nearly complete isolation is maintained through evolutionary time

Strong differences in levels of genetic diversity can complicate estimating gene flow between *H. hecale* and *H. ismenius*. Therefore, we assessed levels of genetic diversity π and Tajima’s D in 50kb windows. As expected from previous research on the silvaniform clade (de Cara et al. 2023), genetic diversity was approximately 2.85 times as high in *H. hecale* (range = 0.0024 to 0.00248) as in *H. ismenius* (range = 0.00080 to 0.00087). Negative Tajima’s D were observed in both *H. hecale* and *H. ismenius* with averaged values approximately 1.5 times lower in *H. ismenius* than in *H. hecale,* indicating an expansion in the latter species. These differences in patterns of diversity suggest that effective population sizes may be different in the two species and that *H. ismenius* may not be at equilibrium. These conditions may further complicate the interpretation of traditional tests to detect gene flow. Thus, we tested whether gene flow occurred during the species demographic history by inferring the demographic history of the species and comparing scenarios with or without gene flow. To do so, we extended the previously developed two-population models implemented in the software DILS (Demographic-Inference with Linked Selection, Fraïsse et al. 2021) in order to consider four populations under four main demographic scenarios and include the possibility of (asymmetric) gene flow between species and/or subpopulations. We modified the coalescent simulation pipeline accordingly and included ABBA-BABA-related summary statistics that are informative to reveal localised gene flow. Last, our models included the confounding effects of barriers to gene flow that affect the effective migration rate (Barton & Bengtsson, 1986), and of linked selection that locally reduces the effective population size (Roux et al. 2016).

We first used a Random-Forest procedure (ABC-RF hereafter; Pudlo et al. 2016) and an ABC neural-network (Csilléry et al. 2012) to compare statistically a model of divergence followed by gene flow (SC or Secondary Contact) to a model of divergence without gene flow (SI or Strict Isolation, **Fig S07**). Statistical comparison was performed hierarchically to improve interpretability. First, we compared SI models integrating the effect of linked selection (modelled as a local reduction in Ne, SI-*Nhetero*) versus a null model without local reduction in *Ne* along the genome (SI-*Nhomo*). Similarly we compared SC models integrating both the confounding effect of linked selection as well as barriers to gene flow, locally reducing the effective migration rate (*m*), against models integrating only one of the two effects and against a null model ignoring these confounders (resulting in four alternative models, namely SC*-NhomoMhomo* for homogeneous effective population size and homogeneous gene flow, SC-*Mhetero* for heterogeneous effective population size only, SC-*Nhetero* for heterogeneous migration only and SC-*MheteroNhetero* for both effects).

Both the ABC and ABC-RF strongly support a model of linked selection [SI-*Nhetero* (posterior probability ∼ 1, **Table S03**)] as well as the different models of secondary contact [SC (-*Nhetero, -Mhetero, -MheteroNhetero,* posterior probability ∼ 1, **Table S04**)] as compared to the alternative null model. Further analyses based on the relationship between diversity, divergence, differentiation and recombination metrics further support the impact of linked selection (see **Fig S04-S05**). In addition, ABC-RF confusion matrices indicated that our classifier was able to accurately discriminate among these alternative models (OOB-prior rate = 6% for SI, **Table S02**). As expected, due to the similarity of the heterogeneous versions of the SC models, the OOB-prior rate increased to approximately 33% when considering these models (**Table S04**). Yet, the strong support for these heterogeneous models in our empirical data echoes the general observation in other species (Charlesworth and Jensen, 2021) and we therefore chose to make our next comparison by considering the model including both linked selection and barriers to gene flow (SC*-MheterNhetero*).

We compared seven distinct Secondary Contact models differing in the directionality of gene flow (**Fig S07**). Namely we tested for unidirectional migration from *H hecale melicerta* to *H. ismenius boulleti* (model SC_A_), the reverse direction (model SC_C_), from *H hecale zuleika* to *H. ismenius clarescens* (model SC_B_), the reverse (model SC_D_), for bidirectional gene flow between co-mimics (model SC_AC_) and between non co-mimics (model SC_BD_). The seventh and last model includes all possible directions of migration (model SC_AC-BD_). Here, our model selection procedure clearly rejected models with multiple directions of migration *p*(SC_AC_, SC_BD_) = 0.01, *p*(SC_AC_)= 0.06, *p*(SC_BD_) = 0.04 in favour of the model SC_C_ and SC_B_. The same results were obtained with ABC-RF (**Table S04**). A final comparison between SC_C_ and SC_B_ yielded stronger support for SC_C_ both in the ABC and ABC-RF approach (**Table S04**). Our final model choice test involved a comparison of the SI-*Nhetero* versus SC*_C_MheteteroNHetero.* Here, our two classifiers disagreed: The ABC-RF inferred the SC_C_*MheteroNhetero* as the best model (*p = 0.91*) whereas the ABC classified SI-*Nhetero* as the best (*p = 0.85*). We focused our investigations on the model chosen by the ABC-RF approach. Indeed, our cross-validations revealed that the ABC-RF displayed a lower classification error (1.5% and 5.3% for SC and SI models, respectively) than the ABC- neural network (6% and 22% for SC and SI models, respectively). Moreover, unlike ABC-RF, ABC suffers from a loss of information associated with the choice of a tolerance parameter that can affect the model choice (Robert et al. 2011, **Table S05**, **Fig S08**).We next examined the data to determine if it supported a model where the two species diverged in allopatry (expected under SC) or in the face of ongoing gene flow, as expected under models of isolation with migration (IM, **Fig S07**) or divergence with initial migration (AM, **Fig S07**). To address this question, we constructed two IM models, one with gene flow between co-mimics and one with gene flow between non co-mimics (**Fig S07**) and a model of ancient migration (AM), and compared the best IM models against AM. The best supported model was that of gene flow between co-mimics (*p* = 0.54). Yet, this IM model was clearly rejected in favour of the AM model (*p*(AM) = 0.83). Therefore, we compared this AM model against SC_C_ and found higher support for AM against SC (*p*(SC) = 0.87, supporting the hypothesis of allopatric divergence. All these comparisons were highly robust **(Table S05)**.

We next estimated parameters under the best two models, namely SI and SCc. Larger credible intervals were obtained under SC than SI **(Table S06, Fig S08-09).** Under the SI models, effective population sizes estimated with the ABC for both *H. hecale* and *H. ismenius* were large, being ∼20M in *H. hecale* and less than half this figure in *H. ismenius* (**Fig 3**), concordant with observed levels of genetic diversity (π) in these two species. Differences in mean effective population sizes within subspecies were minor (**Fig 3**). Estimates of gene flow were largely symmetric between *H. hecale* subpopulations and approximately ten times as strong as in *H. ismenius* (**Fig 3**). Interestingly, estimates of effective population size increased largely under models of SC so that the difference between species was not detected any more (**Table S06**). Split time estimates displayed narrow credible intervals (**Fig S09**), and accordingly, *H. hecale zuleika* and *H. hecale melicerta* would have diverged the most recently (∼1.3 million years ago). *H. ismenius boulleti* likely diverged approximately 1.6 My ago from *H. ismenius clarescens.* Yet, considering uncertainty in parameters estimates, it is possible that all subpopulations diverged approximately at the same time (**Table S06**). Finally, *H. hecale* most likely diverged from *H. ismenius* approximately 3.6 My ago. According to our secondary contact model, migration from *H. hecale melicerta* into *H. ismenius boulleti* started approximately 130 KyA [CI = 1.5KyA– 2MyA] at a rate m = 2.15e-05 [95%CI: 5.8e-7 – 7.3e-5]). Other parameters, namely the shape of the beta distribution used to quantify effective population size reduction (Ne) and barriers to gene flow (m) were also difficult to estimate (see also **Fig. S05** for further analyses of linked selection). In all cases, the beta distribution associated with *Ne* suggested that effective population size would have been strongly reduced in regions affected by linked selection. The ABC-RF procedure generally failed to generate any informative posterior estimation, in spite of a very high accuracy on simulated data (R^2^ = 0.96; **Table S07, Fig S08**) suggesting over fitting. We also attempted to estimate parameters through extreme gradient boosting (Xgboost, Chen et al. 2022). This method displayed modest accuracy on trained data (R^2^ = 0.635) but also generally failed to estimate parameters (**Table S07**).

**Figure 3.**
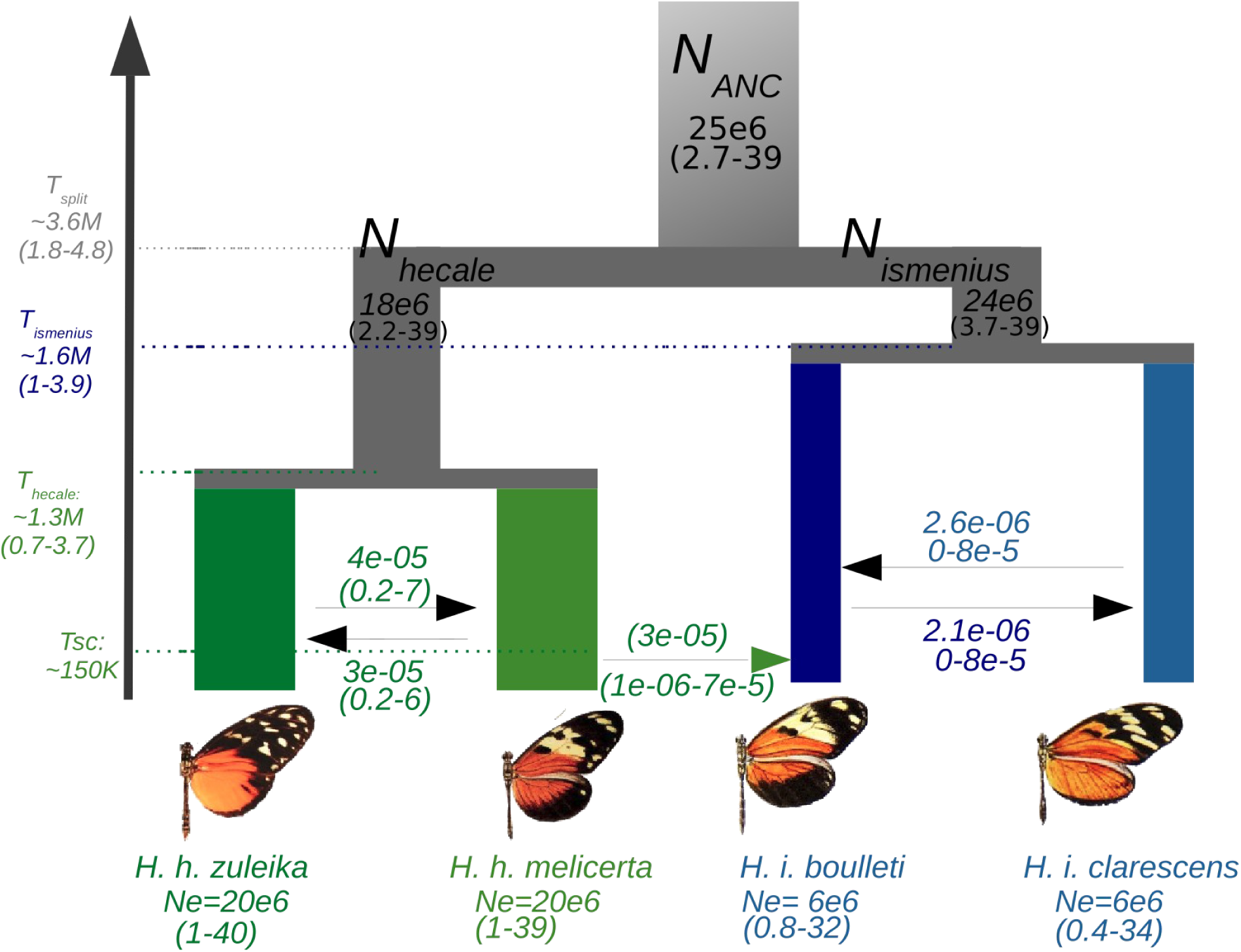
Most likely scenario of divergence inferred from ABC and ABC-RF. The most likely model was obtained through both ABC and ABC-RF. Parameters were estimated with ABC. Arrows display migration rate. The green arrow displays possible migration (inferred with ABC-RF only) between the co-mimics. Split time estimated in years assuming four generations per year. All models included the effect of selection at linked sites and of barriers to gene flow.

### Localised, asymmetric and scattered adaptive introgression between co-mimics

We used the ABBA-BABA family of tests, namely D (Green et al. 2010), *f_d_* (Martin et al. 2015), and D*_FOIL_* (Pease and Hahn 2015), to test for introgression globally and in windows among co-mimics and with other closely related species. At the genome scale, we used all possible donor taxa, including other subspecies such as *H. h. felix, H. i. telchinia* and more distant species (*H. pardalinus* and *H. numata silvana).* As expected from our ABC modelling approach, we found limited evidence for gene flow among the co-mimics at a genome wide scale based on D, *f_d_* or *f_d_*M (**Table S08).** Using the more distantly related *H. pardalinus* as a P1 suggested introgression between *H. h. melicerta* and *H. i. boulleti* as well as between *H. h. zuleika* and *H. i. clarescens.* However, in all these cases, admixture proportions (*f_d_* and *f_d_*M) across the whole genome were not significant. For instance, with introgression from a single individual in a set of four individuals in each population, the maximal *f_d_* value is 0.125 (Van Belleghem et al. 2021).

Next, we looked at *f_d_* values along the genome assessing the topology P1 = *H. i. clarescens*, P2 = *H. i. boulleti,* P3 = *H. h. melicerta* (**Fig 4A**) to test for allele sharing between the co-mimics (P2 and P3) and contrasting this to the non co-mimic pair (*H. i. boulleti*, *H. i. clarescens, H. h. zuleika,* respectively) (**Fig 4B**). In our co-mimic pair comparisons, there were a total of 26 windows with *f_d_* >0.125 (mean length = 15kb), with a striking ∼55 kb peak at the gene *optix* on chromosome 18 whose role on wing patterning is well established. Moreover, strong peaks on chromosome 14 and chromosome 19 were also found (**Fig 4A)**. For the non co-mimics comparisons, we observed a total of 35 shared windows (mean length = 14kb). We observed a very high peak on chromosome 7 which corresponds to a ∼34 kb block containing four genes of unknown function in *H. melpomene.* As in the analysis of co-mimics, a peak is found on chromosome 19, but the signal on chromosome 18 involving *optix* was not observed (**Fig 4B**), suggesting that the wing pattern alleles were not shared between these subspecies. Using more divergent species to increase our power to detect gene flow between co-mimics always showed the existence of a shared introgression signal (**Fig 4A**, **Fig 4B**) at chromosome 19 and an introgression/shared allele signal at *optix* on chromosome 18 among co-mimics but no signal on chromosome 7 suggesting a recent introgression event between *H. h. zuleika* and *H. i. clarescens* (**Fig S10-12**).

**Figure 4.**
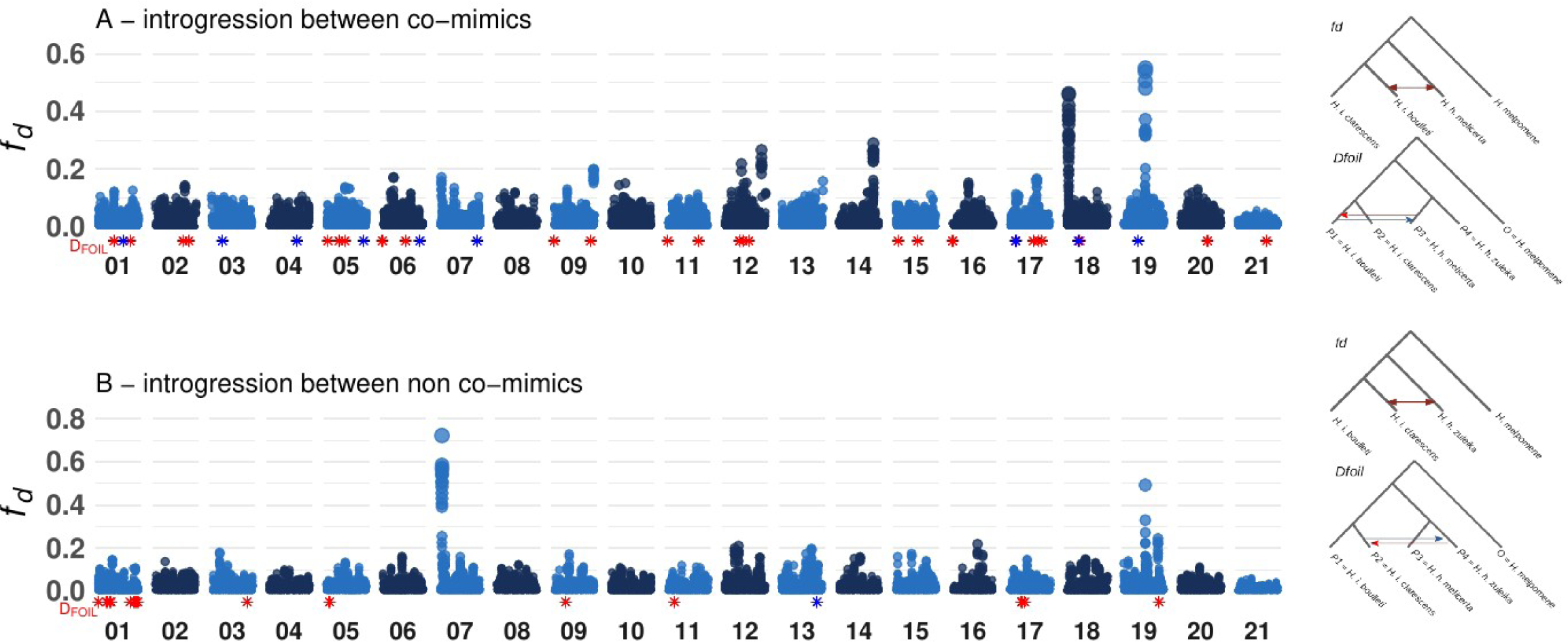
Localised, asymmetric and stronger introgression between co-mimics (A) than non co-mimics (B). 10kb sliding windows *f_d_* values are shown for each comparison, with each chromosome being coloured in dark and light blue. The *f_d_* topology is displayed on the right. Significant D*_FOIL_* values are displayed as blue and red stars for each significant introgression direction and window along the genome. The corresponding D*_FOIL_* topologies and directions are displayed on the right.

The peak on chromosome 19 corresponds to a ∼102.5 kb block containing four chemosensory genes (HmGR50, HmGR47, HmGR48, HmGR51) belonging to the family of Gustatory Receptors (GR genes). A phylogenetic tree of this chromosome 19 region across all species confirmed this introgression signal, with all *H. ismenius* nested within *H. hecale* (Fig S13). This contrasted with the tree inferred from the sex chromosome, which acts as a barrier to gene flow (Van Belleghem et al. 2020; Martin et al. 2019; Fig S14). Similarly, the tree reconstructed around the *optix* gene (chromosome 18, Fig S15) departs from the topology of the species tree, notably with *H. ismenius* appearing sister to *H. hecale*. No particular signal stands out from chromosome 7 and no specific tree was constructed for it.

### More introgression between co-mimics than non co-mimics

To directly test the differences in the directionality of gene flow in relation to mimicry, we used the D*_FOIL_* statistics. Unlike the above ABBA-BABA tests, which necessitate performing separate comparisons between different P1, P2, P3 species and then comparing the number of introgressed windows inferred for each tested, D*_FOIL_* relies on a five taxon phylogeny (here, the two *H. hecale* and the two *H. ismenius* populations, plus an outgroup). It computes all possible combinations of four taxon D-statistics, making it possible to test the excess of gene flow directly, the directionality of gene flow and the taxa involved. To do so, we tested the topology P1, P2, P3, P4, O in which P1 = *H. h. melicerta,* P2 = *H. h. zuleika,* P3 = *H. i. boulleti* and P4 = *H. i. clarescens.* Under stronger gene flow between co-mimics we should expect an excess of P1 → P3, and P3 → P1 over P2 → P4 and P4 → P2.

Since this analysis can only handle a single sequence per population, we performed all possible comparisons between individuals from each species/population and we performed tests in 200 kb windows as it follows a χ^2^ distribution over such a window size. We observed a total of 481 significant windows in either the P1 → P3 direction or P3 → P1 direction versus 218 significant windows when considering P2/P4 supporting the idea that gene flow between co-mimics is stronger (**Figure 4**, **Figure 5, Table S09**). This pattern was not influenced by the removal of *optix*, suggesting that the higher introgression between the co-mimic pair is not solely reflecting the origin of mimicry via introgression of *optix*, but might rather be a result of shared patterns due to higher hybridisation. We observed twice as many significant tests in the P12 → P3 direction as in the P12 → P4 direction (P12 being the ancestral *H. ismenius* branch), suggesting that there has been significant gene flow in ancestral populations, whose footprint may have been removed by selection and recombination.

**Figure 5.**
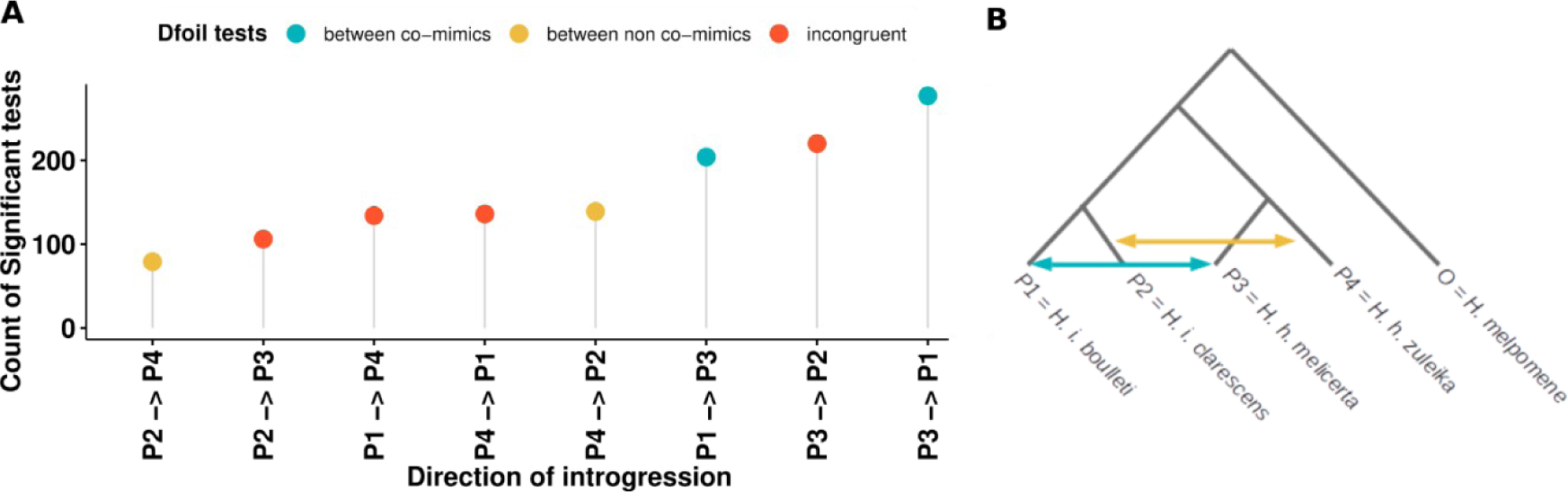
More introgression between co-mimics than non co-mimics revealed by D*_FOIL_* analyses. (**A**) Number of significant D*_FOIL_* tests between each possible direction. (**B**) Topology tested in the D*_FOIL_* analysis. Here we use the term “incongruent” to refer to situations where the inter-group gene flow is between a co-mimic and a non co-mimic species, which does not allow testing our hypotheses.

In order to confirm our previous results, we attempted to refine the distribution of introgression tracts by weighting the support for each fifteen different topologies among the four species with *twisst*, using 50 SNP windows (Martin and Van Belleghem 2017). This analysis largely confirmed that the majority of the genome (84%) supports the species tree (topology 3, **Fig S16**) and this increases to 98.8% when considering additional topologies compatible with the species tree (topology 15 & 10). Overall, one window on chromosome 18 around *optix* was indicative of introgression between co-mimics (topology 9, 12 & 13) and received a weight close to 40% (all topologies are displayed in **Fig S17**). Another topology on chromosome 11 received a weight close to 30%. Considering gene flow between *H. i. clarescens* and *H. h. zuleika,* two windows on chromosomes 15 and 08 received a weight of ∼32 and 34%, respectively. All remaining topologies were equally compatible with gene flow between all species, incomplete lineage sorting or selection against introgression, as expected due to the strength of the species barriers (**Fig S17**). Since we expect introgression to be relatively old and introgression signals to have been broken into smaller segments by recombination, we also tested windows of 25 and 10 SNPs. While results largely confirmed our first inference, 38 highly localised peaks were consistent with the topologies indicating gene flow between co-mimics, again on chromosome 18, but also on a few other chromosomes, including the sex chromosome (relative support > 0.5, **Fig S18-S19**, **Table S10**). Accordingly, we find that the support for the species tree tends to decrease in areas of high recombination (**Fig S20**). To further test the hypothesis that introgressed windows are old, we estimated allele age at *optix* using the ancestral recombination graph. The resulting local tree topologies around *optix* were congruent with gene flow and indicated a coalescence of alleles more recent than the species split, as presented in details in **Fig S21A-B**.

Considering the high levels of genomic divergence and putative reproductive isolation of *H. hecale* and *H. ismenius*, and the indication that mimicry may interfere with reproductive isolation (albeit subtly), we investigated whether the cues and behaviours associated with mate choice in this pair of species showed patterns consistent with genomic inferences.

### Courtship behaviour in *H. hecale*

To investigate mating behaviour in con- and heterospecific encounters, we studied courtship behaviour in *Heliconius hecale.* This species displayed a simple sequence of behavioural units **(Fig 6A; Table S11**; see also Crane, 1957): An aerial phase with both sexes flying fig 6A *localisation*) Tinbergen et al. 1942; Pliske 1975; Brower 1996), followed by an aerial- ground phase where female alights and the male hovers over her (Fig 6A: *hovering*) where both chemical, visual, and tactile cues may be evaluated. A ground-ground phase follows where the male alights, makes physical contact with the female and attempts mating (Fig 6A *attemps)* . At any time, the female may reject the male by adopting a typical *rejection* posture (Fig 6A Obara 1964; Rutowski and Schaefer 1984).

**Figure 6.**
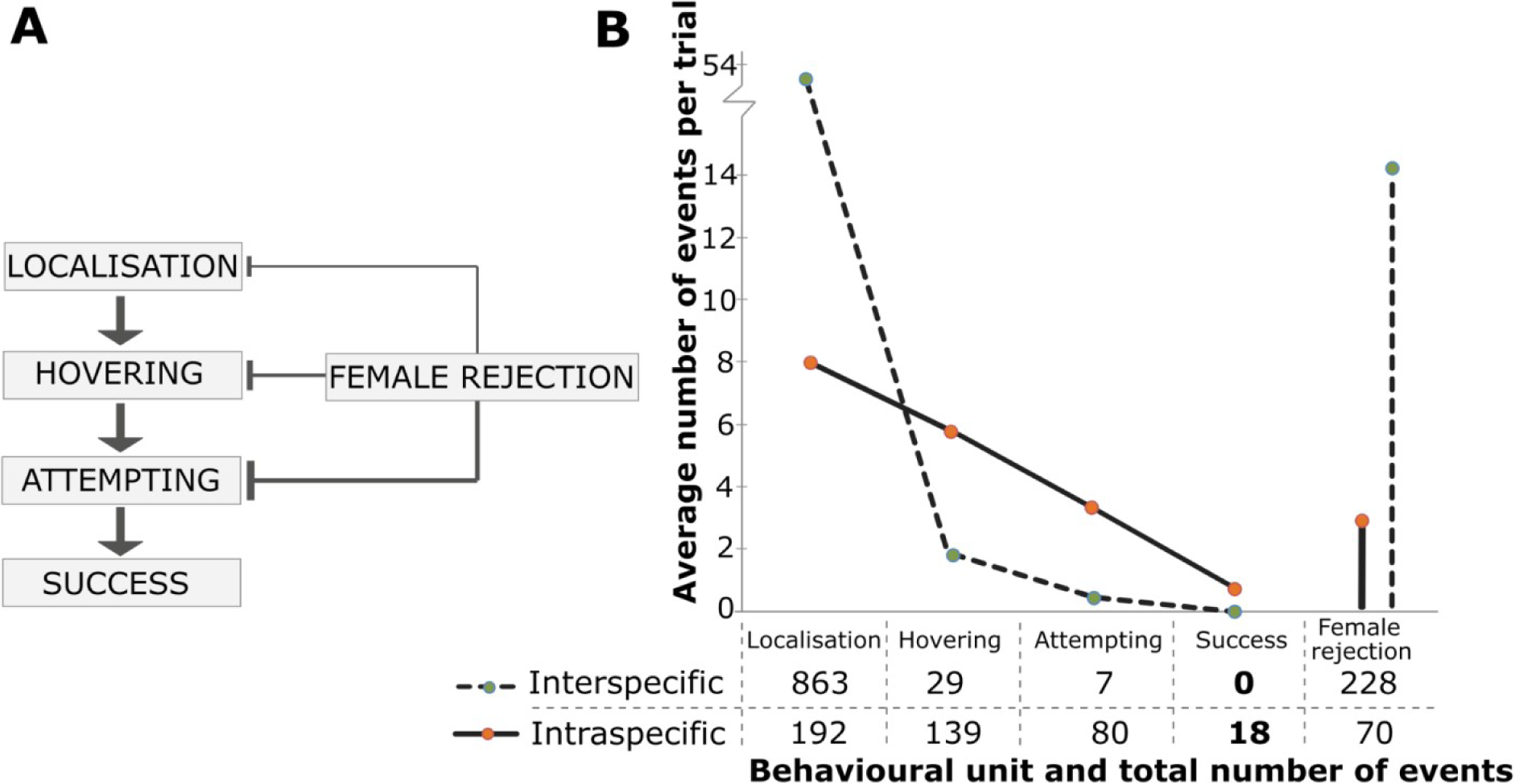
Interspecific recognition relies on short-range cues. (**A**) Main steps of courtship and mating behaviour in *H. hecale.* Courting steps named after the nomenclature used for *Bicyclus anynana* (Nieberding et al. 2008). *Female rejection* is more frequent at later stages. (**B**) Comparison of inter- and intraspecific male-female encounters. The number of events in each main courtship step is shown for interspecific trials (dashed line; *H. h. melicerta* and *H. i. boulleti* tested together, n=16) and intraspecific trials (solid line; only *H. hecale* tested, n=24). Table: total number of events at each behavioural unit registered across the totality of the trials. Y-axis: number of events at each step averaged by the number of trials.

### Interspecific recognition relies on short-range cues

To investigate the signals involved in interspecific recognition, we contrasted inter to intraspecific courtship trials. We observed very strong premating isolation between species and no interspecific mating. This pattern contrasted with intraspecific trials where mating occurred within the first 15 minutes in 75% of our trials (**Fig 6B**). Intraspecific trials showed a gradual transition between main steps, suggesting a gradually increasing selectivity across the courtship sequence. 72.4% of *localisation* events were followed by *hovering* events, and 57.6% of hovering events elicited mating *attempts* in males. In total, 22.5% of such attempts led to mating. By contrast, interspecific trials showed high numbers of approaches but very few were followed by active courtship, showing that heterospecific confusion occurs at long distances but species discrimination is efficient at short range. *Female rejection* is much higher in interspecific experiments, suggesting that both sexes assess the identity and quality of their mates at short range. All interspecific mating *attempting* events (n = 7 by 4 distinct males) were strongly rejected by the female (data not shown). Males approached conspecific and heterospecific female wing models equally (*localisation*; *G*=0.002 *P*=0.96; **Fig 7A, Table S12**). However, males of both species performed the *hovering* behaviour more often on conspecific than on heterospecific models (*G=*21.323 *P<*0.001; **Fig 7B, Table S13**) yet *H. h. melicerta* had a higher propensity to court conspecific models than H. *i. boulleti* males (**Fig 7B**). Finally, we explored the role that colour played in female *H. h. melicerta* behaviour by altering the wing patterns of male *H. h. melicerta.* In these experiments, males with unaltered (*sham*) vs. modified (*treated*) wing patterns (n=10 each) showed equal progression in courtship (**Fig 7C**), and female rejection rates were not significantly different (n*_sham_*=59, n*_treated_*=78, *χ^2^*=2.635 *P=*0.10; **Fig 7C**).

**Figure 7.**
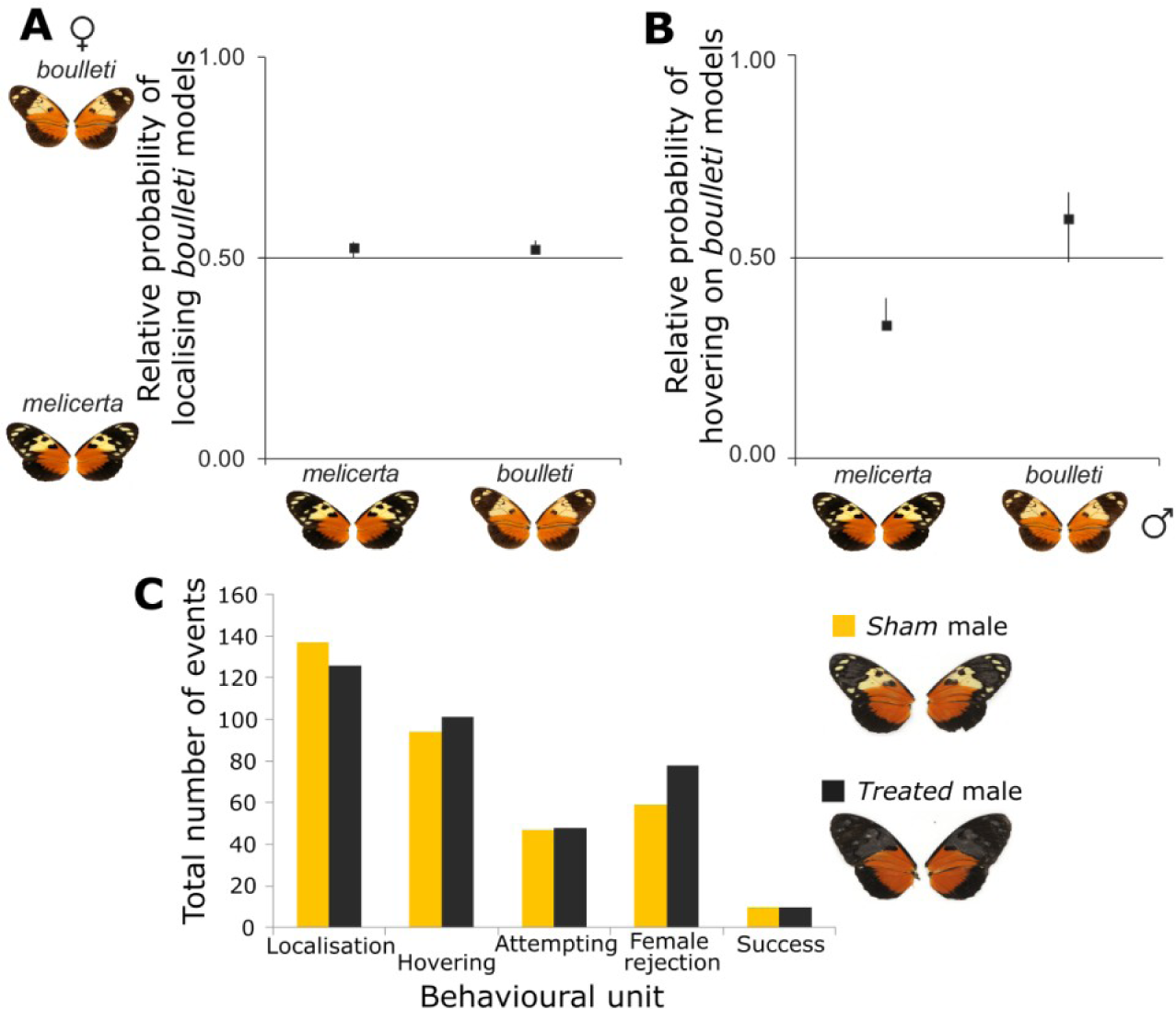
Wing cues contribute to mate discrimination by males but not by females. (**A** and **B**) Male choice of *H. h. melicerta* and *H. i. boulleti* between female wings models of both species. Probability of males approaching (**A**) shortly (*localisation*) and (**B**) sustainedly (*hovering*) the *H. i. boulleti* female wing models, where 1 means a complete choice of *boulleti* models and 0 a preference for *H. h. melicerta* models. A total of 42 *H. h. melicerta* males and 35 *H. i. boulleti* males were tested. Error bars show support limits equivalent to 95% confidence intervals. (**C**) Female choice experiments based on visual cues in *H. hecale*. The total events recorded in each of the main courtship steps during 3 hours long trials are shown on the bar chart. Each *H. hecale* female could choose between two males of its own species and race. One of them was *treated* by modifying its colour pattern (yellow and white patterns on both sides of the forewings blacked with *Sharpie*). The other male (called *sham*) was black-painted on the black regions of the forewing. Only *H. hecale melicerta* male wings are shown here.

### H. hecale melicerta and H. ismenius boulleti differ in their chemical blends

To understand the short range cues causing premating isolation, we investigated chemical differences between them. Chemical extracts from male genitalia (claspers) and female abdominal glands, wings, and cuticles revealed a high variance in the composition and abundance of chemical compounds both between and within sexes and species(**Fig 8AI, 8BI)**. Consistently, non-supervised multivariate analyses showed a good discrimination between males of the two species, and between males and females, but could not distinguish between females of the two species (perMANOVA for abdominal glands, both sexes together: *F=*14.4, *df=*24, *p*=0.000; perMANOVA for wings, both sexes together: *F=*7.6, *df*=17, *p*=0.000; perMANOVA for claspers: *F=*19.6, *df=*11, *p*=0.002, and for wings: *F=*18.7, *df=*10, *p*=0.003, males only; perMANOVA for glands: *F=*1.1, *df=*12, *p*=0.322, and for wings: *F=*0.7, *df=*6, *p*=0.845, females only; NMDS plots, **Fig 8AII, 8BII)**.

**Figure 8.**
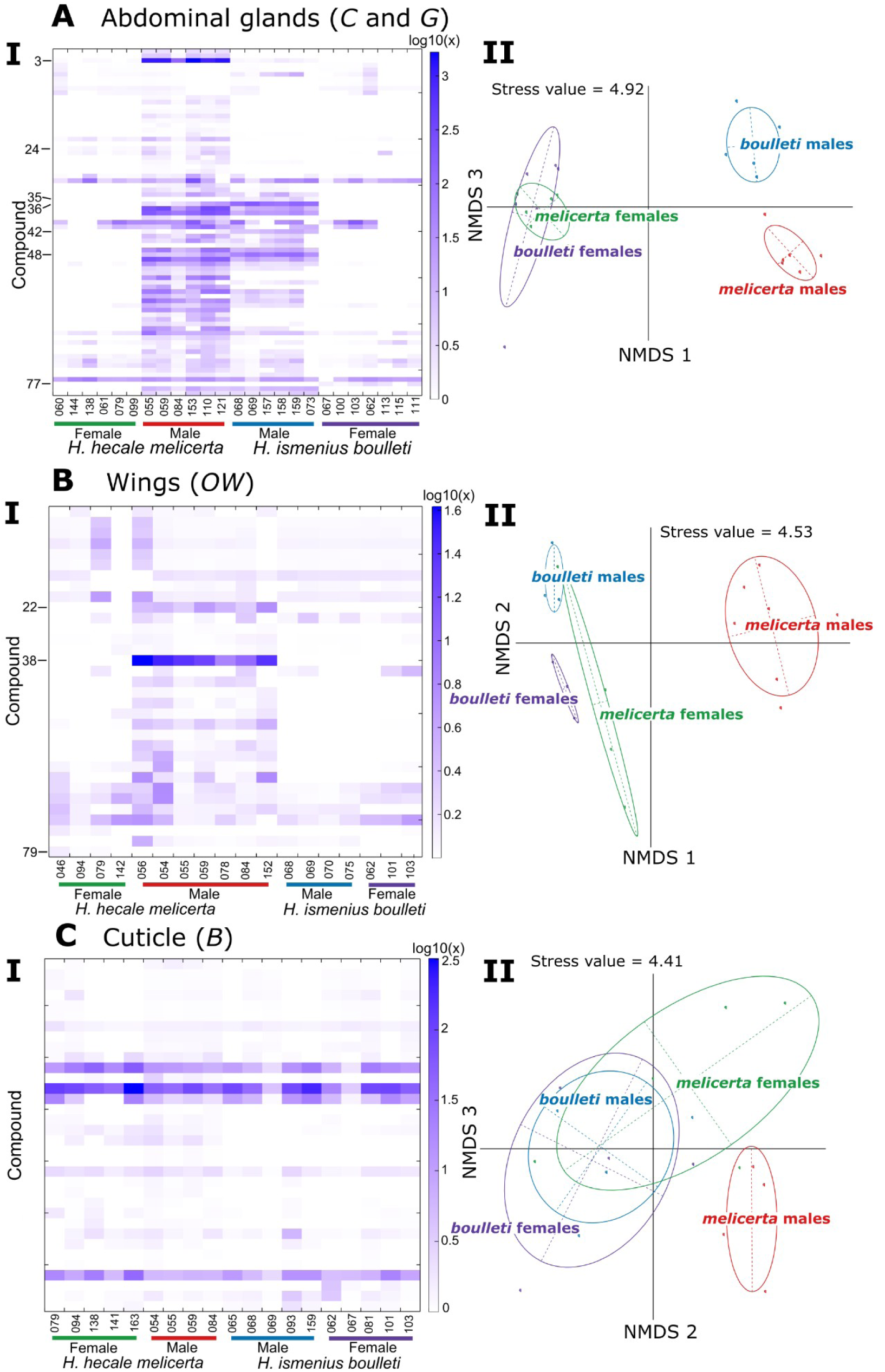
Comparison of the chemical cocktails extracted from three different tissues between *H. hecale melicerta* and *H. ismenius boulleti*. Chemical composition was analysed for abdominal glands (*C* and *G*) (part **A**), wing overlap of both forewings and hindwings together (*OW*) (part **B**) and the cuticle in the base of the wings (*B*) (part **C**). In each part **A**, **B** and **C**, section **I** shows a heatmap indicating the log-transformed [log_10_(x+1)] relative concentration of each compound (horizontal bands) for males and females of both species (columns). Single individuals are labelled under each column. A unique numeric label is indicated for some compounds on the left margin. Section **II** shows non-metric multidimensional scaling (NMDS) ordination of the chemical cocktails of *H. h. melicerta* males and females, and *H. i. boulleti* males and females. Each dot represents a sample of any of these categories. Stress values are shown on the upper part of the graphs. See Table S14-S16 for details on the concentration of the compounds separately. Also see Table S17 for a tentative identification of some of those compounds.

All compounds were associated with a unique numerical label (**Tables S14-S16**). A tentative identification of most compounds (including major peaks of interest) was achieved either at the compound level or at the class-level (**Table S17**) and male wing compoundsclosely matched the identification by Mann et al. (2017) The abundance of several cuticular hydrocarbons, which could play a role during courtship (Millar 2000; Jurenka et al. 2003; Ferveur 2005; Hay-Roe et al. 2007; Dapporto 2007; Heuskin et al. 2014; Klein and Araújo 2010, Loudon and Koehl 2000) consistently differed between species.

Males claspers were chemically more diverse than wing androconia (namely pheromone-producing glands, see also González-Rojas et al. 2020 and Darragh et al. 2020)and most male compounds discriminating between species were present or more concentrated in *H. h. melicerta* and absent or less concentrated in *H. i. boulleti* (**Figs 8AI** and **8BI**, **Tables S14**-**S15**). In *H. h. melicerta* claspers, a known antiaphrodisiac in H. melpomene, H. erato, and H. numata dominated (∼46.9 % of total) but was nearly absent in H. i. boulleti ((*E*)-β-ocimeneor compound number 3; **Fig. 8AI, Table S14;** Darragh et al. 2017, González-Rojas et al. 2020; Schulz et al. 2008; Estrada et al. 2011). Compoundα-ionone, previously found to be specific to *H. ismenius* claspers (Estrada et al. 2011), was not found, perhaps reflecting differences in geographic origins. One compoundin low concentration (number 35) was exclusive to *H. i. boulleti* males, and three were more concentrated in *H. i. boulleti*’s claspers (heneicosene [compound 36], tricosene [48], and compound 42) (**Fig 8AI** and **Table S14**). In total, 42 compounds differed significantly in concentration between male genitalia of the two species (**Fig 8AI; Table S14**). Chemical variation was strongly associated with the *H. hecale-H. ismenius* separation (**Fig S22**). Finally, the *indval* index outlined 44 compounds significantly associated with either species, but especially with *melicerta* males (**Table S14**). A total of 29 compounds were male-specific (**Table S14**).

Female abdominal glands (*AG*) showed few peaks **(Fig 8AI)**. Two low-concentration compounds differed between species. One was present only in *H. i. boulleti* glands (compound 77; **Fig 8AI, Table S14**), and was also exclusive to the cuticle of *H. i. boulleti* females (**Table S16**). This compound was also present on female wings of both species (**Table S15**). The second female gland compound is exclusive to *H. h. melicerta* (compound number 24; **Fig 8AI, Table S14**), but also found in male claspers of *H. h. melicerta.* Nevertheless, possibly due to their low general abundance, neither compound was significantly different between the females of both species (Wilcoxon rank-sum test) or indicative of species status (*indval* index; **Table S14**).

In male wing extracts, two highly volatile compounds with a putative role as air-borne pheromones dominated in *H. h. melicerta* and were absent or found in trace amounts in *H. i. boulleti* (**Fig 8BI** and **Table S15**): heneicosane (∼57.1 % of total; number 38) and hexahydrofarnesyl acetone (∼7.87 % of total; number 22)(as also found by Mann et al. (2017). Here, seven compounds were found in higher abundance in *H. h. melicerta* (**Fig 8BI** and **Table S15**) and contributed to the *H. h. melicerta*-*H. i. boulleti* separation (**Fig 8BII, Fig S23**). The *indval* index identified six compounds associated with *H. h. melicerta* (**Table S15**). Female wing extracts showed fewer compounds of low abundance, highly conserved across species. Only one low concentration compound was exclusive to *H. h. melicerta* wings (2-eicosanyl-5-heptyl-tetrahydrofuran [number 79]) although this difference was not significant (**Table S15**). Overall, two wing compounds were male-specific and one low-concentration compound was female-specific (**Table S15**).

Cuticular extracts (*B*) had fewer compounds than other tissues, and were highly conserved across species and sexes (**Fig 8CI, Table S16**). The clustering by sex or species was not significant (**Fig 8CII**; perMANOVA for all categories together: *F=*1.6, *df=*18, *p*=0.132; perMANOVA for males only: *F=*1.9, *df=*8, *p*=0.140; perMANOVA for females only: *F=*2.3, *df=*9, *p*=0.08).

## Discussion

Introgression among species is known to play a role in mimicry adaptation in *Heliconius* butterflies because of relatively permeable species boundaries following species divergence (Martin et al. 2013; Jay et al. 2018; Martin et al. 2019; Van Belleghem et al. 2021; *Heliconius* Genome Consortium 2012). *H. hecale* and *H. ismenius* fall at the end of the species continuum proposed in Roux et al. (2016) with levels of net sequence divergence as high 0.16 to 0.22. Models based on extant patterns of genome wide variation suggest that the two species likely diverged in allopatry over 3.5 million years ago and then came into secondary contact. Despite very strong premating isolation and divergence in male chemical signals, these genomic observations reinforce the growing realisation that species barriers remain permeable to introgression even among taxa that diverged in the relatively distant past. Our result contrasts with a previous study (Roux et al. 2016) that concluded that none of six pairs of species studied (with similar or higher levels of divergence) exchange genes. Indeed, Roux et al. (2016) found that above a net divergence greater than 0.1, the probability of gene flow was reduced. Yet, strong changes in population size may also affect these estimates by reducing within-population diversity, thus inflating *D_A_*. Here, D*_XY_* levels are similar to the observed *D_A_*, so our results should be robust to the confounding process of population size change. A long isolation period is theoretically expected to favour the accumulation of genetic incompatibilities, *e.g.* Bateson-Dobzhansky-Muller incompatibilities, in particular on the sex chromosome whose absolute divergence was strong, in line with observations in other *Heliconius* species (*e.g.* Van Belleghem et al. 2020).

Our demographic reconstruction provides insight regarding the role of gene flow at the end of the speciation continuum, though we did not include all other closely related *Heliconius* species in our demographic analysis due to the intractability of an ABC model with all putative donors. This may lead to false inferences of ongoing gene flow when there is none (Tricou et al. 2022). To circumvent this caveat we complemented our global inference analysis with several “local” analyses (*f_d_*, *twisst*, ARG) aiming at detecting gene flow along the genome and incorporating more candidate donor species.

Overall, gene flow between *H. hecale* and *H. ismenius* is much reduced relative to the pervasive gene flow observed between the ecologically, behaviourally, and chemically distinct *H. melpomene* and *H. cydno* (Bull et al. 2006; Kronforst et al. 2006; Martin et al. 2013). The reported acceleration of genome-wide divergence accumulation towards later stages of speciation (Kronforst et al. 2013) seems to be illustrated here. *Heliconius hecale* and *H. ismenius* have diverged for ∼3.5 My, which is ∼1.5 My longer than *H. melpomene* and *H. cydno* (Kozak *et al*. 2015).

Despite the overall low level of genome-wide gene flow detected,and indications that speciation is nearly complete, localised introgression windows were nearly twice as frequent in locations where the species share the same warning colour pattern compared to locations where the species diverge in wing pattern, consistent with a role of vision as a mating cue, and of mimicry as a source of sexual confusion. This was also supported by our *D_FOIL_* analysis revealing that the most significant tests of introgression involved co-mimics. One of the strongest signals we observed was around the transcription factor *optix,* determining red-orange colour pattern elements in *Heliconius* generally (Reed et al. 2011), and associated with a QTL for intraspecific differences in the extent of black vs. red-orange hindwing colour in both *H. hecale* and in *H. ismenius* crosses (Huber et al. 2015). Our genealogical analysis here suggests that one allele may have been transferred among species, bringing support for its implication in pattern evolution under mimicry selection. The few, short and localised segments of introgression lead us to hypothesise that mimicry on such a pattern is old. Similarly, a few other windows bearing introgression signals were revealed only when using small window sizes, suggesting an erosion of introgression signals around functional sites by recombination. The local ARGs with a recent TMRCA within *optix* also support this hypothesis.

While our results clearly revealed a strong chemical differentiation, an interesting introgression signal shared between both co-mimics and non co-mimics stands out on chromosome 19 within a block of ∼102 Kb harbouring four chemosensory genes belonging to the same family of gustatory receptors. Such families of genes were studied recently (van Schooten et al. 2020) and are hypothesised to play key roles in speciation in *Heliconius.* Interestingly, we found that they displayed increased admixture (*f_d_*, D*_FOIL_*) as well as a phylogenetic topology that does not support the species tree (**Fig 4)**. Interestingly, high admixture values were also inferred for 3 of the genes in a recent study on speciation between *H. melpomene* and *H. cydno* (van Schooten et al. 2020). Clearly, the role of these genes would be worthy of further investigations. For instance, a detailed investigation of the ancestral recombination graph (ARG) of these regions, together with functional validations may be useful. In all cases, the strong chemical differentiation that we observed mirrors the strong genome wide divergence.

Speciation in *Heliconius* is multifactorial (Mérot et al. 2017), so the strong genomic divergence we observe likely results from the built-up and strengthening of multiple barriers to gene flow. Colour pattern is often an early acting barrier in the mating sequence, and may be acting as an initial yet somewhat permeable barrier to gene flow during speciation (Van Belleghem et al. 2020). Our behavioural assays suggest mimicry confuses males visually in their approach behaviour from a long distance, yet short-range cues clearly play a central role in premating isolation. The chemical profiles of male *H. ismenius* and *H. hecale* are both extremely diverse and highly divergent, while female profiles are nearly identical, which is consistent with a role for selection acting on chemical signals produced by male wings and genitalia and assessed by females during courtship (Darragh et al. 2017; González-Rojas et al. 2020; Bergström and Lundgren 1973; Schulz et al. 1993; Andersson et al. 2007; Costanzo and Monteiro 2007; Nieberding et al. 2008). Note, however, that males approach both con-and heterospecific female wings at the same rate, but engage in courtship with conspecifics at a higher rate, highlighting the use of short-range cues to determine whether or not to court. It is possible that these cues are partly visual and may reflect subtle differences between species in pattern or how pattern is perceived (Dell’Aglio et al. 2018). Other aspects of ecological differentiation, such as host use or microhabitat segregation, may also contribute to the low probability of hybridisation and gene flow in those species, and may warrant detailed investigations.

Taken together, our results indicate that the pair of species studied here falls in the range of the speciation continuum where reproductive isolation is nearly complete, and may involve the recruitment of multiple barriers to hybridisation, including chemical signatures. Contrary to other pairs of taxa showing a high level of incomplete lineage sorting throughout the genome, here the genomes are indeed well sorted. Yet we were able to detect localised segments with increased gene flow among co-mimics as compared to non co-mimics, including around the wing patterning gene *optix* (Reed et al. 2011; Zhang et al. 2017; Huber et al. 2015). The small length of introgressed ancestry together with the ARG suggest a relatively distant event of adaptive introgression at *optix*. If this introgression is indeed relatively old, mimicry might therefore have evolved at a time where the species were less isolated genetically, followed by further chemical differentiation and the near completion of speciation. In this scenario, the origin of mimicry, associated with optix, could have caused increased introgression. Not mutually exclusive with such a model, ancient genome-wide introgression followed by a gradual erosion of introgression signals may also cause similar signals and cannot be formally rejected. A better understanding of the process by which barriers to gene flow accumulate during speciation requires an accurate understanding of the trajectory of key genes determining hybridisation during the history of lineage divergence.

## Materials and methods

### Specimens

In Eastern Panama, *H. hecale* and *H. ismenius* are excellent co-mimics of each other (subspecies *H. h. melicerta* and *H. i. boulleti*) and both display a pattern made of orange (proximal) and yellow elements (distal) bordered by a thick black margin and a black wing tip. In Western Panama, those species display distinct patterns and join different mimicry rings. *H. h. zuleika* shows a pattern with a largely black forewing and prominent yellow dots, and a merely orange hindwing. In contrast, *H i. clarescens* shows a largely orange wing pattern with black and yellow stripes alternating in the distal part of the forewing.

For genomic sequencing, three to six specimens of each subspecies were collected from Darién (East of Panama) and near David, Chiriquí (West of Panama), totalling eighteen specimens (**Fig 1** and **Table S01**). Bodies were preserved in a NaCl-saturated DMSO solution and wings in paper envelopes. In addition, four samples of *H. hecale felix* collected in Peru, four *H. hecale clearei* from Venezuela and two *H. ismenius telchinia* from Panama were used four broader context in PCA and structure analyses and as control allopatric populations of the study species in our test of gene flow in ABBA-BABA related methods (**Fig 1** and **Table S01**). Closely related silvaniforms, namely 15 *H. pardalinus* and 17 *H. numata silvana* (**Fig 1A**) were also used when assessing gene flow in ABBA-BABA-related methods (detailed in **Table S01**). Finally, a set of 12 *H. melpomene* individuals were used as an outgroup. For behavioural and chemical data *H. h. melicerta* and *H. i. boulleti* specimens were collected in Darién (Eastern Panama) and around Gamboa, Colón (Central Panama), while *H. h. zuleika* specimens were collected from Chiriquí (Western Panama) (**Fig 1**). Stocks of these races were reared at the Smithsonian Tropical Research Institute in Gamboa (Panama) (Huber et al. 2015).

### Population genomics

#### DNA extraction

DNA was isolated from preserved bodies using the DNeasy Blood and Tissue Kit (Qiagen). Separate Illumina paired-end libraries were generated according to the manufacturer’s protocol (Illumina Inc.). Each library was shotgun sequenced to an average coverage of ∼25X on an Illumina Hi-Seq 2000 with 2 x 100-base read length.

#### Genotyping and variant calling

Reads were trimmed using fastp (Chen et al. 2018), aligned to the *H. melpomene v2.5* genome with bwa-mem v.0.7.13 (Li 2013) and filtered with samtools (Li et al. 2009) requiring a minimum quality of 20. Duplicates were removed using picard (http://broadinstitute.github.io/picard/). SNP calling was then performed using GATK V4.1.9 (DePristo et al. 2011).Variable and invariable sites across the whole genome were included. Following GATK Best Practices, all sites that did not match the following criterion were marked: MQ < 30, QD < 2, FS > 60, MQRankSum < -20, ReadPosRankSum < 10, ReadPosRankSum > 10. The dataset with variant and invariant sites was used for ABC analyses below. SNPs were then extracted from the dataset and used for all remaining analyses. This dataset was filtered to keep SNPs with a minor allele count of 2. Genotypes without a mean depth between 5 and 60 (*i.e.* mean+2 standard deviations) and a genotype quality above 30 were set as missing, resulting in ∼5.5 millions SNPs.

#### Phasing and recombination rate estimates

Phasing was performed using beagle v.5.1 (Browning and Browning 2007) with default parameters. LDHat software (McVean et al. 2002) was then used to estimate effective recombination rates (⍴ = 4.Ne.r where r represents the recombination rate per generation and *Ne* is the effective population size) along the genome. The genome was split in chunks of 2000 SNPs with overlapping windows of 500 SNPs to compute recombination rate and data were then merged together. Recombination rates were averaged into 250-kb windows using a custom Python script. Recombination rate was inferred for each species separately (*H. hecale, H. ismenius*) as well as for outgroups (*H. numata, H. pardalinus*) to analyse how conserved the landscape is along the divergence continuum (see Supplementary Results). Genetic diversity (π) and Tajima’s D were measured for each species along 50 kb windows after converting whole genome vcf file into fasta using seq_stat (https://github.com/QuentinRougemont/PiNPiS).

#### Population relationships

Genetic relationships among individuals were assessed with a PCA using the R package ade4 (Dray and Dufour 2007) based on a LD-pruned dataset, further thinned to keep SNPs spaced at least 500 bp from each other (365K SNPs). Population treeness was tested using Treemix (Pickrell and Pritchard 2012) with the same LD-pruned dataset and allowing up to 20 migration events. The model that explained the highest level of variance when adding a migration edge was chosen. Five hundred bootstrap replicates of the best model were performed to obtain robustness of the nodes.

#### Global demographic history

To test whether the species have diverged in the presence or absence of gene flow through historical times, coalescent simulations were used in an ABC framework. Our models account for the confounding effect of linked selection (locally reducing the effective population size) and of barriers to gene flow (locally reducing the effective migration rate; Roux et al. 2016; Rougemont and Bernatchez 2018).The two-population models previously developed were extended to a four-population model, representative of the four focal populations studied. It includes ABBA-BABA-related statistics explicitly developed for detecting gene flow at different time scales. A statistical comparison of a model of divergence without gene flow (SI) was performed against allopatric divergence followed by secondary contact (SC) (**Fig S07 A-B**). The best of the two models was then compared against models of sympatric divergence with gene flow, namely Isolation w. Migration (IM) and Ancient Migration (AM) (**Fig S07 C-D**). The SI model assumes that an ancestral population of size *N_ANC_* splits instantaneously at time Tsplit into two daughter populations of constant effective population size *N_12_* and *N_34_* and no gene flow. Afterwards populations continue to diverge and give rise to two subpopulations (*H. h. zuleika* and *H. h melicerta,* of size *N_1_* and *N_2_*) at time T_12_ and two other subpopulations (*H. i. boulleti* and *H. i. clarescens,* of size *N_3_* and *N_4_)* at time T_34_ (**Fig S07**). These subpopulations are connected by continuous gene flow, *M* = *4N_0_.m* with *M*_x←y_ being the number of migrants from population Y into population X at each generation. Under the SC model the two species diverged under a period of strict isolation before undergoing a secondary contact from Tsc until present. All directionalities of secondary contacts were tested, from a single directionality of migration to a set of fully connected populations (**Fig S07 B**) which enabled to test the hypothesis of gene flow either between co-mimics (direction A <-> C in **Fig S07 B)** or between non co-mimics (direction B <-> D in **Fig S07 B**). Under AM the instantaneous split of the ancestral population is followed by a period of gene flow from Tsplit until the split of the two daughter populations. Gene flow is stopped when the first pairs of populations split in two subpopulations (**Fig S07 C)**. As for SI, gene flow was allowed between subpopulations within *H. hecale,* and within *H. ismenius,* respectively. In the IM there is continuous gene flow following the split of the ancestral population into two. Following the split of the subpopulations within species we then allowed for gene flow either between co-mimics (direction A <-> C in **Fig S07 D**) or between the non-co-mimics (direction B <-> D in **Fig S07 D**). In all models, migration priors were sampled independently for each direction, enabling asymmetric introgression between species or populations.

#### Coalescent simulations

For each model, 1 × 10^6^ simulations of datasets matching the sample size of each locus was performed using msnsam (Ross-Ibarra et al. 2008) a modified version of the coalescent simulator ms (Hudson 2002). Selected markers can provide additional information regarding a species history (Bierne et al. 2013; Roux et al. 2013). Therefore, analyses were restricted to the coding sites (CDS) from the *Heliconius melpomene* genome, using a total of 6,600 informative loci falling in those CDS and passing our filtering criterion. Large and uninformative priors were used. Effective population size for all current and ancestral populations were sampled uniformly on independent intervals [0–40,000,000]. Priors for the divergence time (T_split_) were uniformly sampled on the interval [0–5,000,000], so that T_12_, T_34_ and the various T_sc_ were constrained to be chosen within this interval. For the homogeneous version of the migration rate, M was sampled on the interval [0–40] independently for each direction of migrations. We then used beta distribution hyperprior to model the heterogeneity in effective population sizes and migration rate (Roux et al. 2016; Rougemont and Bernatchez 2018). Heterogeneity in migration was modelled using a beta distribution as a hyperprior shaped by two parameters; α randomly sampled on the interval [0–20] and βrandomly sampled on the interval [0–200]. A value was then independently assigned to each locus. For the heterogeneity of Ne, α percent of loci evolved neutrally and share a value uniformly sampled on the interval [0–1]. Therefore 1–α percent of loci were assumed to evolve non neutrally. Their Ne values were thus drawn from a beta distribution defined on the interval [0–20], as defined on the homogeneous version. Ne values were independently drawn for N_anc_, N_1_, and N_2_, N_3_, N_4,_ but shared the same shape parameters (Roux et al. 2016). Priors were generated using a Python version of priorgen software (Ross-Ibarra et al. 2008). A mutation rate of 2.9 × 10^-9^ mutations per bp and generation was used, as estimated for *H. melpomene* (Keightley et al. 2015).

#### Summary statistics

A modified version of mscalc was used to compute summary statistics (Ross-Ibarra et al. 2008; Ross-Ibarra et al. 2009; Roux et al. 2011). This version, coded in pypy for faster computation, provided the average and standard deviation of statistics computed over loci. The number of biallelic sites, the number of fixed differences between species X and Y (sf), the number of exclusively polymorphic positions in species X (sx), the number of shared biallelic positions between species X and Y (ss), the levels population genetic diversity (Tajima’s π and Watterson’s 𝛳) and Tajima’s D were included, as well pair-wise statistics between X and Y populations: levels of population differentiation (*F_ST_*), population absolute and net nucleotide divergence (*D_XY_* and *D_A_*) respectively, the smallest divergence measured between one individual from X and one from Y (minDiv), the highest divergence measured between one individual from X and one from Y (maxDiv) as well as a range of statistics known to be informative with regards to introgression, namely ABBA-BABA’ D statistics (Durand et al. 2011), *f_d_* statistics and D_FOIL_ (Pease and Hahn 2015). Finally, between-population Pearson correlation coefficients in genetic diversity (π and 𝛳) were also computed.

#### Random Forest and Neural Networks model choice

Two approaches were used to perform model choice. First, posterior probabilities were computed using a classic ABC procedure. The “best” 1000 simulations (out of 1 million) were retained, weighted by an Epanechnikov kernel peaking when Sobs = Ssim and a neural network that performs a non-linear multivariate regression where the model is a response variable to be inferred. Fifty feedforward neural networks were used for training with 15 hidden layers in the regression. Ten replicates ABC model choices were performed to obtain the average posterior probability of each model. The R package “abc” (Csilléry et al. 2012) was used for the model choice procedure. Because classical ABC may suffer from the curse of dimensionality, especially for a complex model with many summary statistics as here, a Random Forest approach was also used for model choice (*e.g*. (Rougemont et al. 2016). This approach has the benefit of directly assessing the proportion of correct model choice by the algorithm. A forest made of 500 trees was constructed and a linear discrimination analysis was included as additional summary statistics. Computations were performed using ABCRF package (Pudlo et al. 2016). All available summary statistics were used to construct a confusion matrix, estimate classification error and perform the model choice. The IM and AM models were compared among each other and to other models using only the ABCRF procedure. The whole pipeline is available at https://github.com/QuentinRougemont/ABC_4pop

#### Parameter inference

The posterior probabilities of each parameter value were estimated using the neural network procedure with nonlinear regressions of the parameters on the summary statistics using 50 feed-forwards neural networks and 15 hidden layers after a logit transformation of the parameters. A tolerance of 0.001 was applied. The ABC-RF was also used to estimate demographic parameters of the best models using 1,000 trees to build a forest.

#### Genomic differentiation and divergence analyses to test for linked selection

Divergence among species and populations was measured by estimating nucleotide diversity (π), levels of divergence (*D_XY_*) and of genetic differentiation (*F_ST_*) in windows of 250kb along the genome using Python scripts from Martin et al. (2013). Patterns of heterogeneous genome-wide divergence can be due to genetic hitchhiking of neutral alleles linked to selective sweeps (Smith and Haigh, 1974) or to background selection (BGS, Charlesworth et al. 1993). These combined effects, referred to as linked selection, reduce polymorphism at sites closely linked to advantageous or deleterious variants (Payseur and Nachman 2002). The intensity of selection on linked loci is mostly modulated by variation in local recombination rate and by gene density (Kaplan et al. 1989). Under linked selection, diversity (π, *D_XY_*) and differentiation (*F_ST_*) metrics are expected to be positively and negatively correlated with genome-wide variation in recombination rate respectively (Payseur and Nachman 2002). Correlation among π, *D_XY_* and *F_ST_* were therefore tested among species to test for the conservation of these landscapes (**Fig S04 A-C**). A PCA was then used to obtain a synthetic view across all populations and species of our estimates of π, *D_XY_* and *F_ST_* separately. Next, Spearman correlations based on each (z-transformed) variable projected on the PC1 axis separately were carried out to look at correlation among variables (**Fig S04 D, Fig S05**).

#### Testing for interspecific gene flow

Gene flow was evaluated between species (*H. hecale* vs. *H. ismenius*), between co-mimics (*H. h. melicerta* vs. *H. i. boulleti*) and between non co-mimics (*H. h. zuleika* vs. *H. i. clarescens*) using the same ABBA-BABA test as implemented in our ABC pipeline, namely we calculated *D* over the whole genome (Green et al. 2010; Durand et al. 2011) and *f_d_* (Martin et al. 2013) over the whole genome and in windows. Given three taxa *P1*, *P2*, *P3* where *P1* and *P2* are sister taxa, and an outgroup *O* with the following topology [(*P1*, *P2*), *P3*]*, O* we tested for gene flow between *P2* (the target population) and *P3* the donor population. *H. melpomene* was used as outgroup and two configurations were tested to quantify gene flow between the co-mimics *H. h. melicerta* and *H. i. boulleti* on the one hand, and between the non-mimetic pair *H. h. zuleika* and *H. i. clarescens* on the other hand. In addition, other more distant P1 species were tested, namely *H. pardalinus* and *H. numata* as well as *H. ismenius telchinia.* A 1Mb block jack-knifing approach was implemented to calculate the mean and variance of *D* and *f_d_*, and to test whether *D* differed significantly from zero. *f_d_* statistics were calculated in 10kb sliding windows along the genome using custom scripts developed by Martin et al. (2013).

To investigate relationships among individuals at regions showing strong *f_d_* values, local phylogenies using RAxML-NG (Kozlov et al. 2019) were built. A GTRCAT model and 100 bootstraps were used.

Analyses were complemented with the D*_FOIL_* statistics (Pease and Hahn, 2015), that enable testing the topology [(P1,P2)(P3,P4)],O), with a potential higher power to test the directionality of introgression than the above test. In addition, this test enables the direct consideration of the four co-mimics and non co-mimics species in a single test. As for our ABC pipeline,the original D*_FOIL_* statistics was modified to work on allele frequency of multiple individuals instead of the sequence of a single sample.

Finally, the different topology relationships between co-mimics and non co-mimics were quantified using the tree weighting approach implemented in *twisst* (Martin and Van Belleghem 2017). The proportion of each of the 15 different possible topologies was computed (detailed in **Fig S16**), with a particular focus on topologies reflecting gene flow between the co-mimic species (*H. h. melicerta* and *H. h. boulleti*), or gene flow between the non co-mimics (*H. h. zuleika vs H. i. clarescens*). The complete method, which considers all subtrees, was used to estimate the exact weighting to the full tree. Topologies were then classified according to whether they reflect: the true species trees (1 topology), partial tree with incongruence in *H. hecale* only (2 topologies), partial tree with incongruence in *H. ismenius* only (2 topologies) or perfect geography (Geographic perfect (*H. i. b*oulleti + *H. h. melicerta* AND *H. i. clarescens* + *H. h. zuleika* together): 1 topology), Geographic partial East (*H. i. boulleti* + *H. h. melicerta* together NOT *H. i. clarescens* + *H. h. zuleika*): 2 topologies corresponding to higher gene flow between co-mimics, Geographic partial West (*H. i. clarescens* + *H. h. zuleika* together NOT bou+mel): 2 topologies, Incongruent (boul+zul AND/OR cla+mel): 5 topologies (but see **Fig S16**). Window sizes of 100, 50, 25, 15 and 10 SNPs were tested.

#### ARG reconstruction at *optix*

The Ancestral Recombination Graph (ARG) was estimated in order to investigate variation in population history of *optix* (population size, topology, allele age and gene flow) using ARGweaver (Rasmussen et al. 2014). A constant mutation rate was used along the genome of 2.9 × 10e^-9^ mutations per bp and generation, estimated in *H. melpomene* (Keightley et al. 2015) and recombination rate of 1e^-8^ per bp/generation. Utility scripts provided in Hubisz & Siepel (2020) were modified to plot the resulting trees as well as the TMRA, RTH, π levels and population size. A large window around *optix* (660 kb to 750 kb, with *optix* being located between 705,604 and 706,407 bp) was used to provide further context. Dataset included both variant and invariant sites.

### Behavioural and Chemistry analyses

#### Courtship description

To define the sequence and prevalence of behavioural steps during courtship and mating in *H. hecale* butterflies, behaviour was video-recorded until mating happened (successful courtships, n=4) or until at least one whole sequence including all steps but mating had been registered (unsuccessful courtships, n=13). Our observations were supported by other behavioural experiments in this (detailed below) and previous studies concerning other species in the subfamily Heliconiinae (Klein and Araújo 2010; Crane 1955, 1957; Rutowski and Schaefer, 1984; Mega and Araújo, 2010).

#### Inter- and intraspecific sexual interactions

To unravel the signals involved in species recognition between *H. hecale melicerta* and *H. ismenius boulleti*, no-choice interspecific encounter experiments were performed. Three mature (>5 days old) conspecific males of one species were put in a 2x2x2m cage with one newly-emerged (a few hours old) heterospecific female. All male courtship events and female responses occurring during 15-min trials were recorded on video. Main courtship steps (**Fig 6**) were registered for 9 trials with *H. h. melicerta* males and 7 with *H. i. boulleti* males. Intraspecific no-choice trials were run as controls using *H. hecale* (n=24), and involved two mature males and two virgin females of each of the two races *H. h. melicerta* and *H. h. zuleika*.

#### Male choice using female wing models

To determine the importance of female wing cues in courtship decisions by males, choice experiments were performed where males of one species were presented with female wing models of both species. *Localisation* events (close approach to models) and *hovering* events (sustained stationary flight close to the models for >=3 seconds) were registered. Sexually mature, reared (n=44) or field-caught (n=33) males were tested multiple times, on different days, to obtain independent behavioural registrations for each individual. A total of 38 *H. h*. *melicerta* and 32 *H. i. boulleti* were characterised by three registrations, plus 4 *H. h*. *melicerta* and 3 *H. i. boulleti* males had only two registrations each. Models were made with real unwashed female wings, thus including visual and chemical signals. Wings were connected to a plastic wire and moved in a fluttering motion inside a 2x2x2 m cage. Trials were run for 5 minutes, presenting two models simultaneously, and switching model positions in the middle. *Localisation* and *hovering* were analysed separately. The relative probability of males courting *boulleti* female models rather than *melicerta* models was estimated using a likelihood function as follows (Edwards 1972; Jiggins et al. 2001):

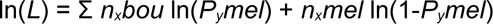

where *P_y_mel* denotes the probability of courtship by male *y* towards *melicerta* wings, *n_x_bou* the total number of events of male *x* directed towards *boulleti* models, and *n_x_mel* towards *melicerta* models. The solver option in Excel (Microsoft) was used to estimate the probabilities of male courtship by numerically searching for values of *P_y_mel* that maximised ln(*L*). Support limits (asymptotically equivalent to 95% confidence intervals) were assessed by looking for values that decreased ln(*L*) by two units (Edwards, 1972). To compare courtship across species, a model with equal relative probabilities (*Pmel*=*Pbou*) was compared to a model with probabilities calculated separately for each species (*Pmel*≠*Pbou*), using a likelihood ratio test assuming a *χ*^2^-distribution with one degree of freedom for the test statistic G=2Δln(*L*).

#### Female choice based on colour

To assess whether visual cues are important in mating acceptance by *Heliconius* females, experiments were performed involving a virgin *H. hecale* female (subspecies *melicerta* or *zuleika*) and two mature males of their own subspecies with two different colouration treatments (**Fig 7C**). *Treated* males were painted with black ink, using a *Sharpie* pen, following Kemp (2007), to cover the white and yellow patches of dorsal and ventral forewings. *Sham* males, or controls, were painted on the black region of the forewing, covering a similar surface as for *treated* males but avoiding pattern modification. Experiments were run in 1x2x2m cages. In this case females were over one day old, thought to be better able to reject courting males than just-emerged females. A total of 26 trials were performed (13 with *H. h. zuleika* females and 13 with *melicerta* females) resulting in 20 successful courtships. Direct observation of sexual behaviour was carried out until mating occurred or for a maximum period of 3 hours, and all main courting events were registered.

#### Chemical analyses

To evaluate differences in chemical signalling between *H. h. melicerta* and *H. i. boulleti*, chemical compounds were extracted from three different tissues. In males, wing regions where forewing and hindwing overlap (*OW*), and abdominal glands (claspers, *C*) have been proposed to contain pheromone-producing glands (androconia) releasing volatile signals by *Heliconius* males (Emsley 1963; Brown 1981; Schulz et al. 2008; Estrada et al. 2011; Darragh et al. 2017). Similarly, male’s genital claspers are often open during courtship, putatively diffusing volatile signals perceived by the female (Schulz et al. 2008; Estrada et al. 2011). Finally, female rejection might involve the evagination of abdominal glands, which in mated females contain volatile anti-aphrodisiacs compounds (*e.g.* Rutowski and Schaefer 1984, Schulz et al. 2008). Our description of courtship behaviour suggests they might be used both in *H. hecale* and *H. ismenius*. In virgin females, abdominal glands (*G*) were analysed given their outstanding role in producing sex pheromones in several species of Lepidoptera. Additionally, the cuticle of the body close to the base of the wings (*B*), was analysed because of the presence of small bristles which might be involved in the production of chemical signals (i. e. hairpencils, Boppré, 1978; Bacquet et al. 2015) diffused by the wings. Tissues from 3-7 individuals were extracted separately from virgin females and mature males of both species, right after sacrificing the individuals. Pieces of tissue were immersed into 130-200 µl of hexane containing dodecane 100 ng/µl as internal standard. Extractions were not stopped. Elution samples were analysed with gas chromatography coupled to mass spectrometry (GC-MS) with a Bruker Autosampler SCION SQ 436-GC, using a non-polar Rtx-5 MS fused silica capillary column (0.32 mm i.d., 30 m long, 0.25 µm thick film) and helium as the gas carrier. Volumes of 0.5 µL were injected automatically in a split-less mode with the injector temperature at 250°C (see Mérot et al. 2015 for more details). Peak area was used as a measure of compound abundance relative to the concentration of dodecane in the sample. Since chemical receptor sensitivity is unknown for these butterflies, low-concentration peaks were not excluded from our descriptions. However, peaks that were too low for quantification were treated as missing data for a given extract. Quantification values were log-transformed [log_10_(x+1)], allowing comparison of peaks with large differences in abundance (Lecocq et al. 2013).

#### Statistical analysis of chemical data

Chemical similarities and differences between species and sexes were visualised with heatmap graphs on the relative abundance of the compounds present in each tissue separately, using *Matlab R2012a Student Version*. Since chemical bouquets are likely to be perceived as composite signals, multivariate analyses were performed on the concentration of all compounds present in each tissue separately, where each compound is treated as a variable. For each tissue, analysis of compositional similarity of chemical cocktails between species was performed with sexes pooled as well as separating by sex, using non-supervised non-metric multidimensional scaling (NMDS) ordination implemented in the *vegan* (Oksanen et al. 2017) packages in *R*. Prior to NMDS, the Bray-Curtis distance matrix of the chemical composition was computed. The appropriate number of dimensions was chosen for each dataset by plotting a “scree diagram” (stress versus number of dimensions). The significance of chemical differences between species or sexes was assessed by a discriminant analysis on the Bray-Curtis similarity matrix, using a permutational multivariate analysis of variance (perMANOVA; Anderson, 2001) implemented in the *vegan* package (Oksanen et al. 2017), with 1000 permutations. Whenever categories were significantly different, partial least squares discriminant analysis (PLS-DA) was applied to the dataset using *R* packages *mixOmics* and *DiscriMiner* to identify variables (*i.e*. compounds) most diagnostic of a given category. The *indval* index, which gives the probability of association between a compound and a given category, was also estimated by computing indicator values that suggest which compounds are indicative of a given category. For that purpose, the *indval* function in the *R* package *labdsv* (Roberts 2013) was used, following Heuskin et al. (2014). Finally, a comparison of the per-compound concentration was performed with a Kruskal-Wallis test coupled with a pairwise Wilcoxon rank-sum test, and Bonferroni correction for multiple testing. To facilitate the identification of the compounds, linear retention indices (LRI) were calculated for each compound relative to the retention times of a mix of alkane standards (C10-C26). So far, a partial number of compounds were tentatively identified by comparison of retention indices and mass spectra with those of authentic reference standards (Sigma-Aldrich Corporation), with published databases (Wiley Registry of Mass Spectral data-7^th^ edition and NIST Chemistry WebBook; NIST Chemistry WebBook 2015) and with compounds previously identified (Estrada et al. 2011; Schulz et al. 1998; Schulz et al. 2008). Other peaks of interest remain to be identified and the current identification of some of the compounds remains to be confirmed before manipulative assays of their role in species recognition between *H. hecale* and *H. ismenius* can be envisioned.

## Data Availability

Raw data are available on NCBI (see details in **Table S01**). Filtered vcf will be deposited on Zenodo. Scripts to reproduce all genomics analyses are available on the first author github page.

