## supplementary Materials for "Subtle introgression footprints at the end of the speciation continuum in a clade of *Heliconius* butterflies"

Table S14. Compositional similarities and differences between the chemical blends extracted from abdominal glands of males (*C*) and females (*AG*) of *H. hecale melicerta* and *H. ismenius boulleti*.53

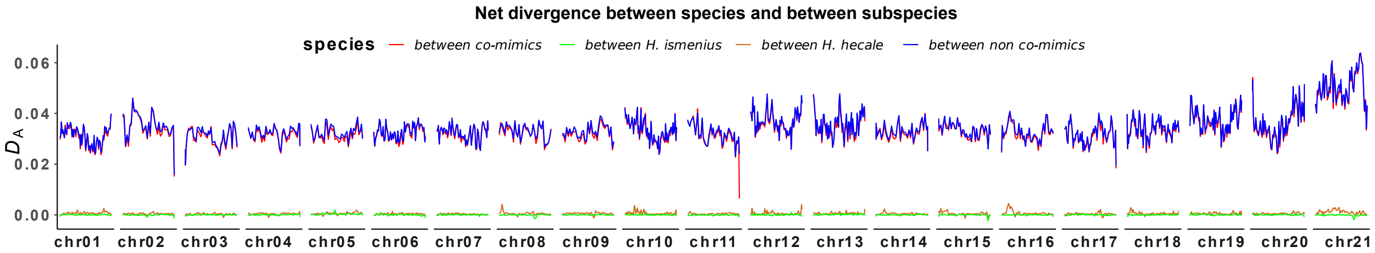

**Figure S01. Net divergence ($D_A$) between species.**

Landscape along each chromosome between sympatric co-mimics (*H. h. melicerta* vs *H. i. boulleti*, red line) and between non co-mimic species (*H. h. zuleika vs H. i. clarescens*, blue line). Genetic differentiation between subspecies of each species is also shown (*H. h. melicerta* vs *H. h. zuleika*, chocolate; *H. i. boulleti* vs *H. i. clarescens*, green). Computation averaged in 250 kb windows.

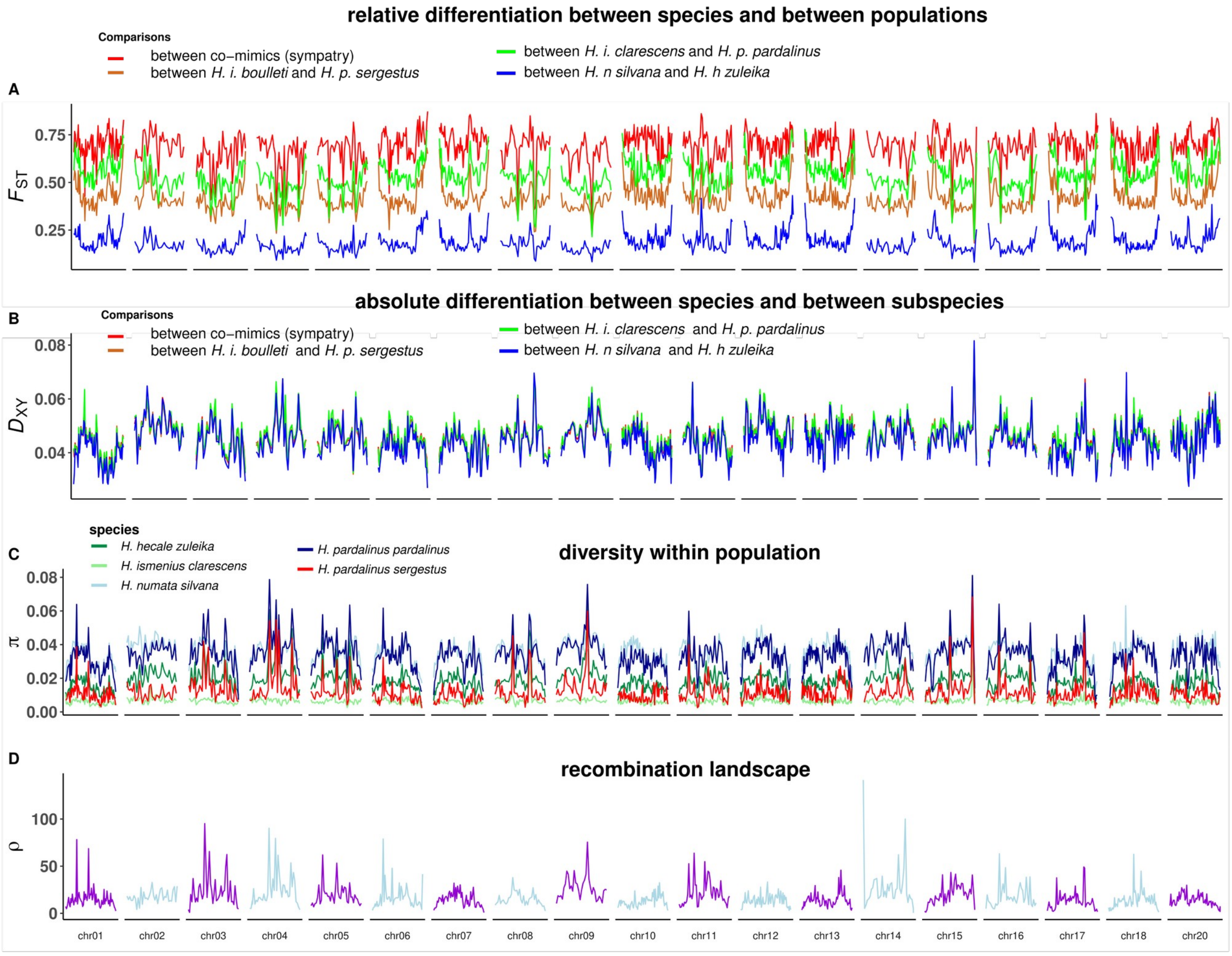

**Figure S02. Conserved divergence landscape across *Heliconius* butterflies from the silvaniform clade.**

Variation along the whole genome is displayed along the x-axis. Variation in each summary statistic is displayed along the y-axis and computed in 250 kb windows to show broad scale variation among species, with **A**) $F_{ST}$ among sets of different species pairs, **B**) $D_{XY}$ among species pairs, **C**) distribution of nucleotide diversity (π), and **D**) landscape of population scale recombination (ρ = 4Ne*r). Each colour represents a species or a pairwise comparison in the case of $D_{XY}$ and $F_{ST.}$

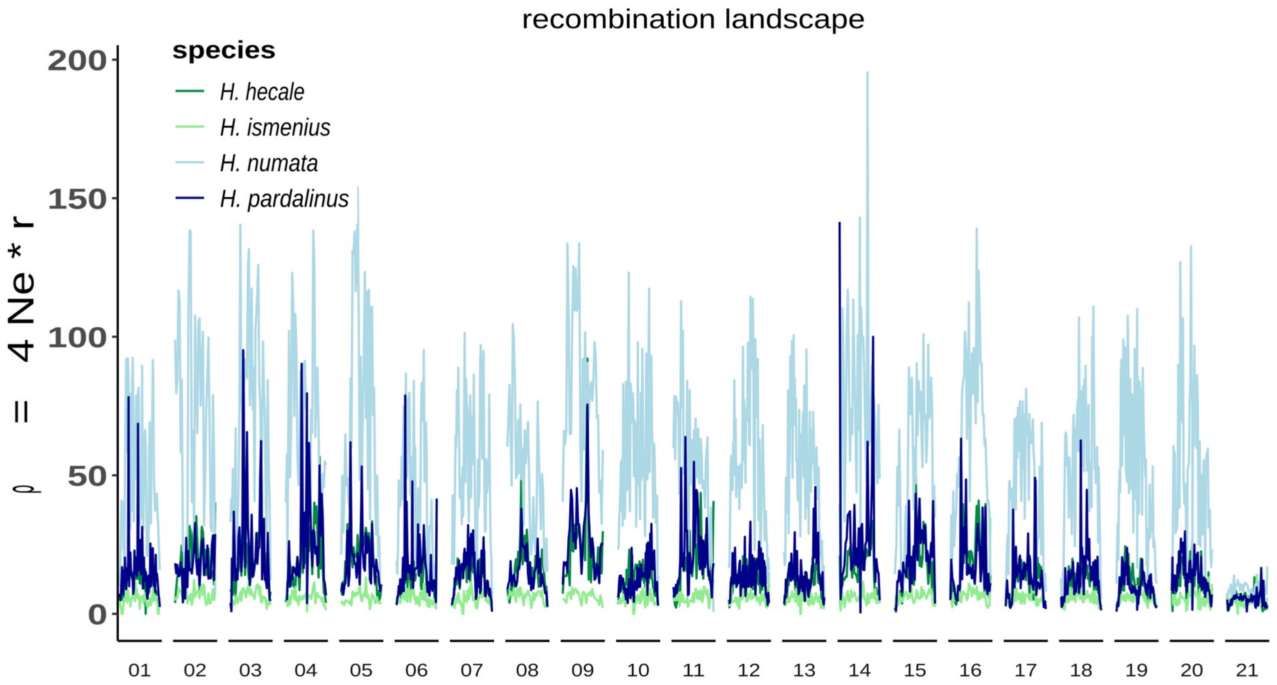

**Figure S03. Conserved landscape of population scale recombination at a broad genome spatial scale among multiple species.**

Each colour represents a species. The whole genome is represented on the x-axis and variation in recombination by chromosome is displayed along the y-axis. Species with higher levels of nucleotide diversity (see Figure 2 in main text), a proxy of effective population size, display higher population-scale recombination.

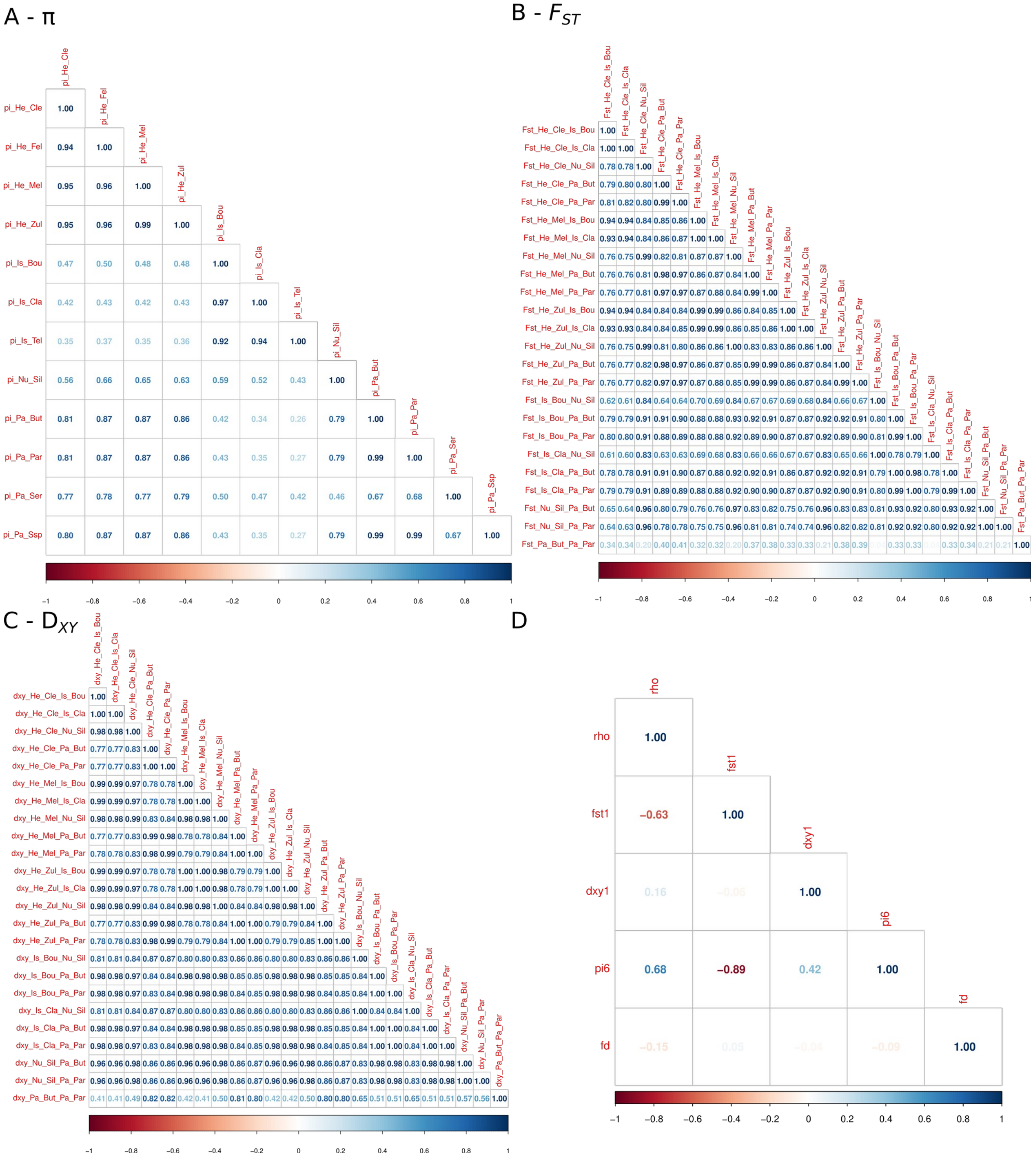

**Figure S04. Covariation among different metrics highlight the conserved landscape of divergence among species.**

Each cell displays the correlation coefficient between pairs of variables. Darker colours depicts to stronger correlation (red = negative, blue = positive). **A**) Correlation in nucleotide diversity among species and populations. **B**) Correlation in genetic differentiation ($F_{ST}$) between each pair of species included in this study. **C**) Correlation in genetic divergence ($D_{XY}$) between each pair of species included in this study. **D**) Correlation among the first PC-axis of each summary statistic (i.e. rhi, pi, fst, Dxy and Fd). PC-axes summarise genomic variation in each statistic observed across all species (see details in methods).

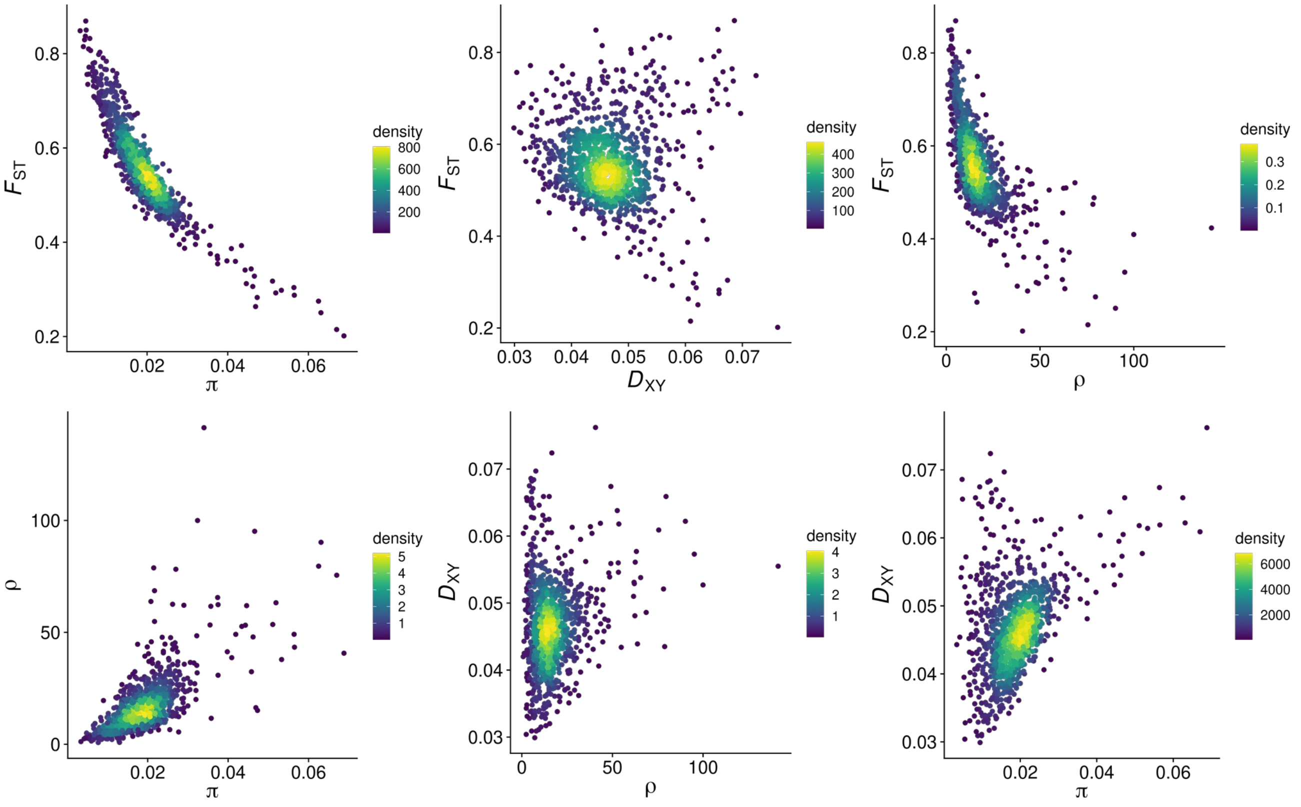

**Figure S05. Conserved landscape arise likely due to linked selection.**

Displayed are pairwise correlation between summary statistics of diversity (i.e. nucleotide diversity (π) and between species diversity ($D_{XY}$)), and differentiation ($F_{ST}$) measures along the genome of each species in 250 kb windows. Under linked selection, both π and $D_{XY}$ should correlate positively with recombination rate while $F_{ST}$ should correlate negatively with genome-wide variation in recombination rate (ρ), respectively, and with the density of targets (e.g. genes, regulatory regions) that are subject to selection (Payseur & Nachman, 2002, Charlesworth et al. 1993; Kaplan et al. 1989; Nordborg et al. 1996).

Charlesworth B, Morgan MT, Charlesworth D. The effect of deleterious mutations on neutral molecular variation. Genetics. 1993;134: 1289–1303.

Kaplan NL, Hudson RR, Langley CH. The “hitchhiking effect” revisited. Genetics. 1989;123: 887–899.

Nordborg M, Charlesworth B, Charlesworth D. The effect of recombination on background selection. Genet Res. 1996;67: 159–174.

Payseur BA, Nachman MW. Natural selection at linked sites in humans. Gene. 2002;300: 31–42.

Hudson RR, Kaplan NL. Deleterious Background Selection with Recombination. Genetics. 1995;141: 1605–1617.

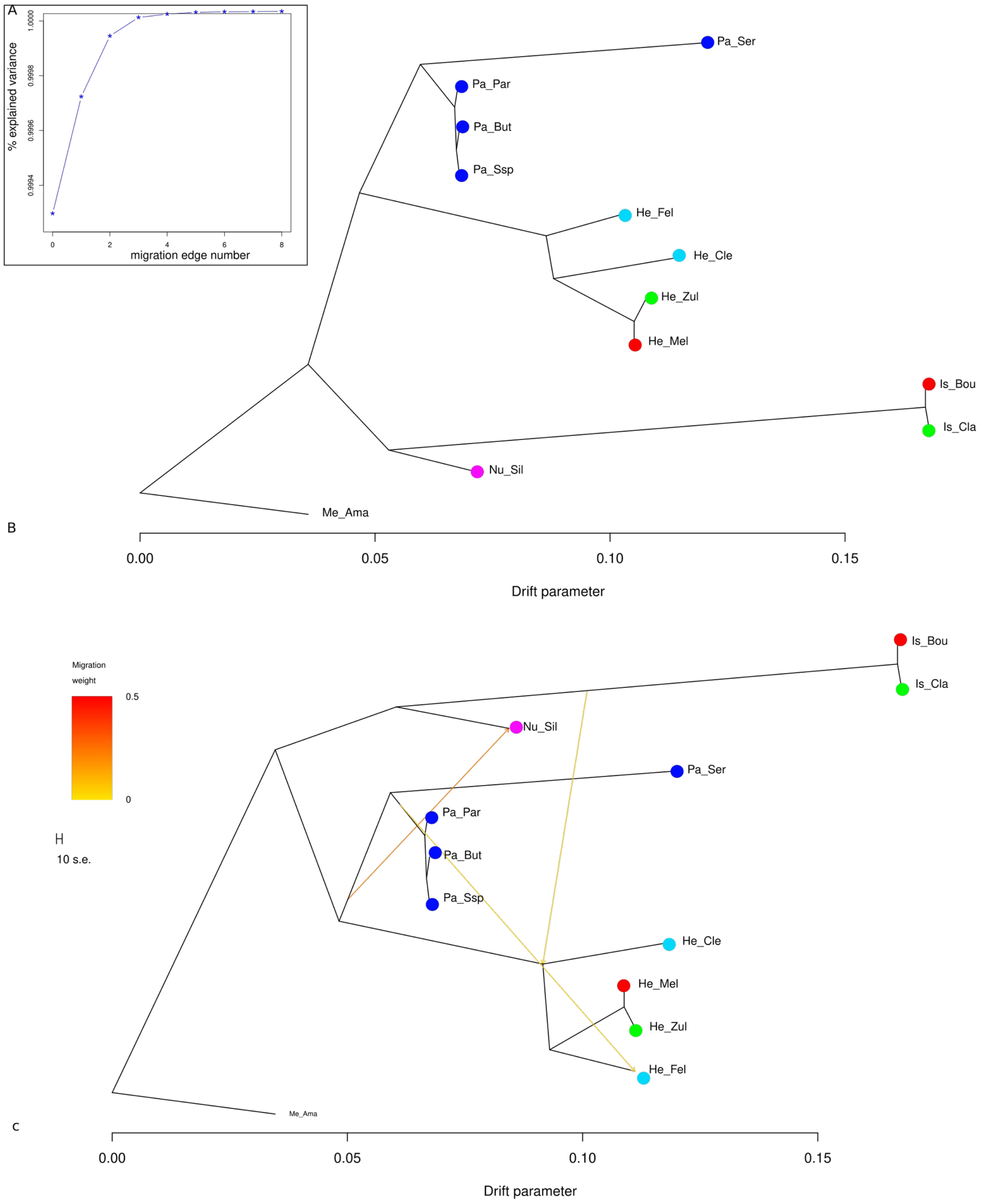

**Figure S06. Treemix inference of migration event along branches of the tree.**

**A**) Proportion of variance explained in the covariance matrix of allelic frequency by treemix as migration edges (i.e. events of migrations) are added to the tree. **B**) Model without gene flow and **C**) model with three-migration events, capturing potentially ancestral migration. Each coloured dot represents a genus except for red and green dots which represent co-mimics and non-comimcs, respectively. *ismenius* displayed the highest drift. Abbreviation: Pa = *H. pardalinus* (Par = subspecies *pardalinus*, But = *butleri*, Ssp = *Ssp. Nov*). Is = *H. ismenius* (Cla = *clarescens*, Bou = *boulleti*), He = h*ecale* (Mel = *melicerta*, Zul = *zuleika*, Fel = *felix*), Nu_Sil = *H. numata silvana*.

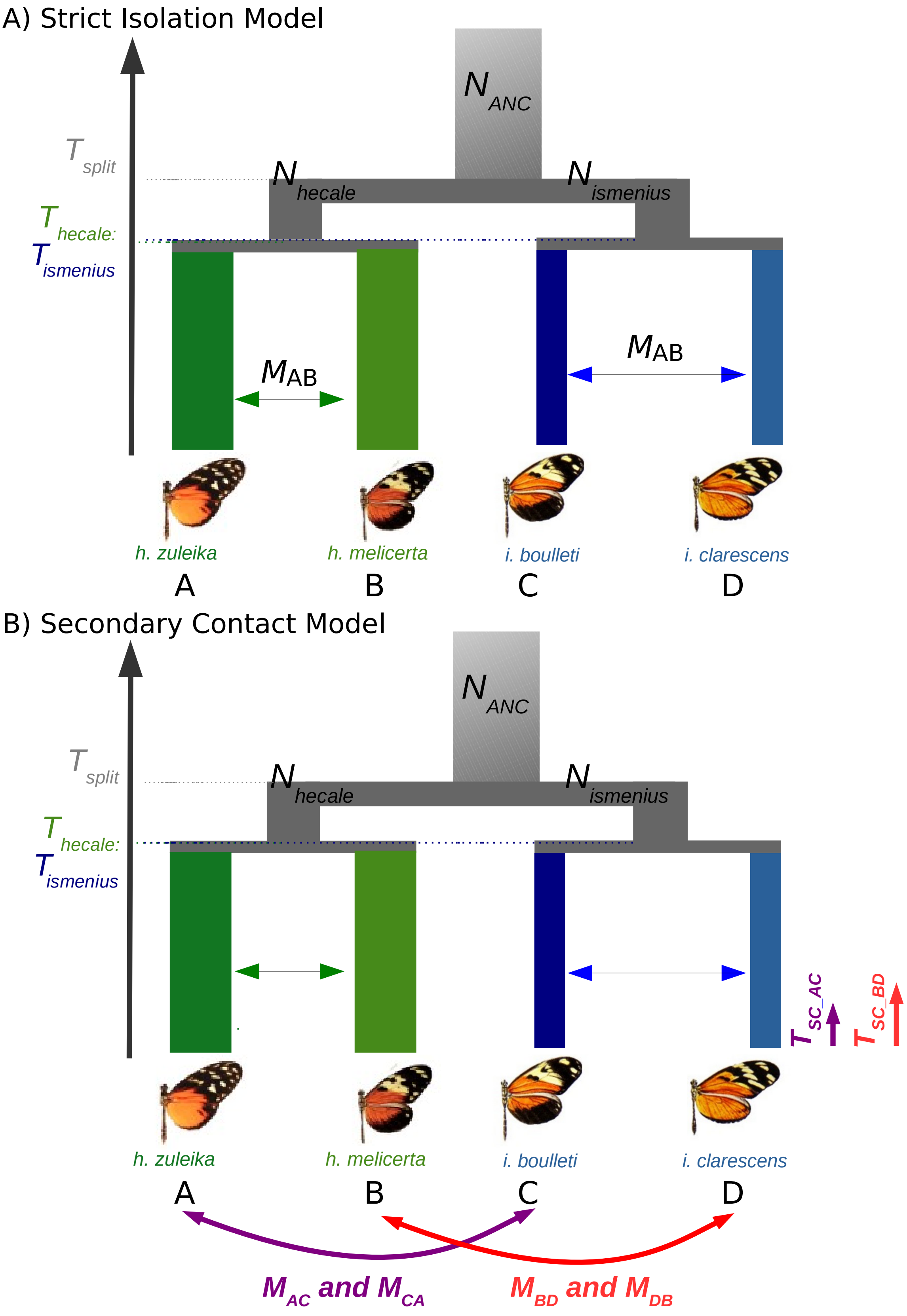
A) Strict Isolation Model
N ANC
T split
N hecale
N ismenius
T hecale:
T ismenius
MAB
MAB
h. zuleika
h. melicerta
i. boulleti
i. clarescens
A
B
C
D
B) Secondary Contact Model
N ANC
T split
N hecale
N ismenius
T hecale:
T ismenius
T SC_AC
T SC_BD
h. zuleika
h. melicerta
i. boulleti
i. clarescens
A
B
C
D
M AC and M CA
M BD and M DB

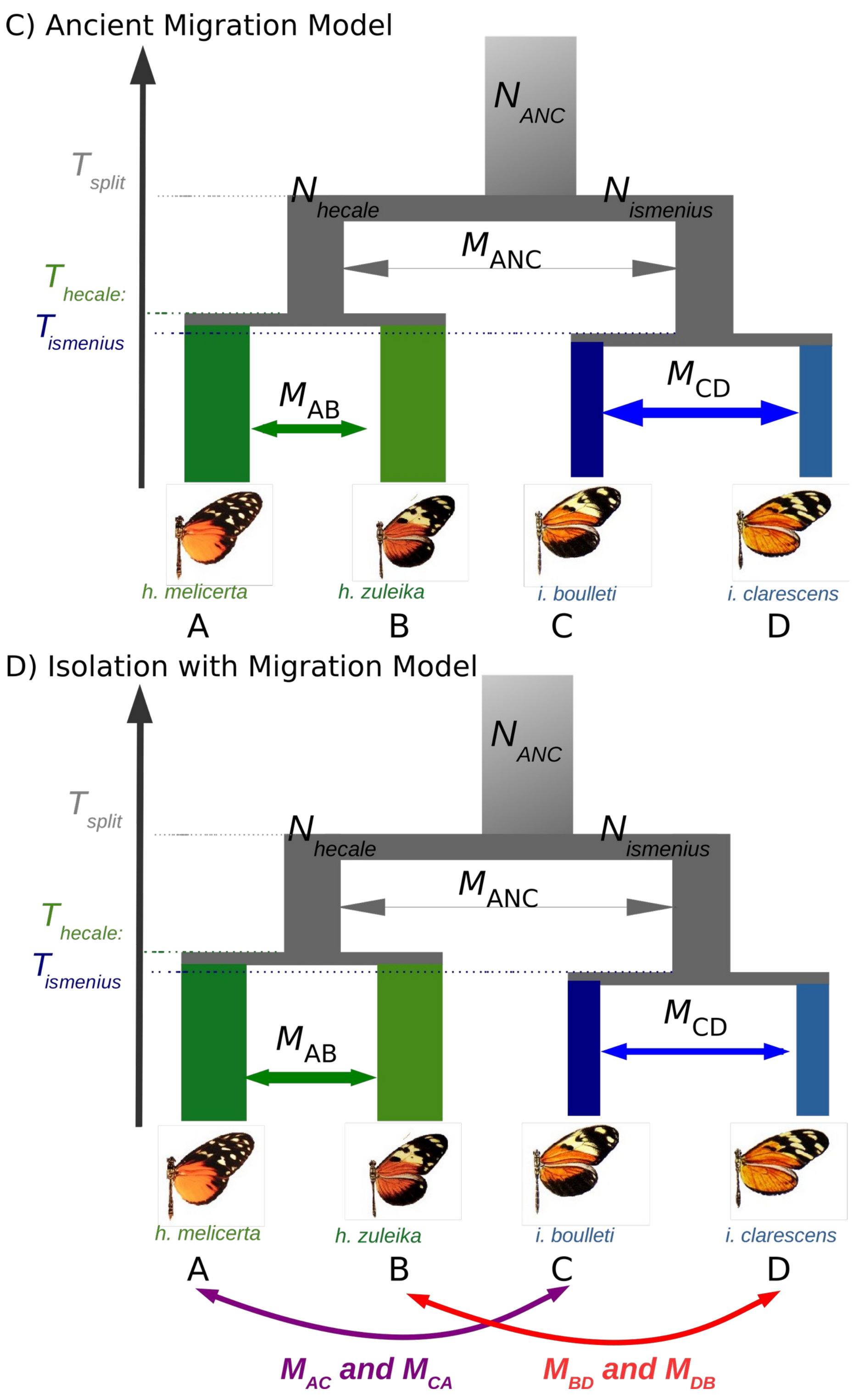

**Figure S07. Simplified representation of the major demographic models being tested.**

**A**) Model of strict isolation and **B**) model of secondary contact. Each model accounts for linked selection (see Methods). Seven directions of migration were tested in the SC model:
(1) A ← C, (2) C ← A, (3) A ↔ C, (4) B ← D, (5) D ← B, (6) B ↔ D, (7) AC ↔ BD.
The major demographic parameters are displayed through simplified representation. C) Model of Ancient Migration with gene flow in the ancestral populations of *hecale* and *ismenius;* and D) model of Isolation with Migration. In the IM models, two versions testing gene flow either between A ↔ C or B ↔ D were tested to test the hypothesis of differential gene flow with bidirectional asymmetric introgression

A)

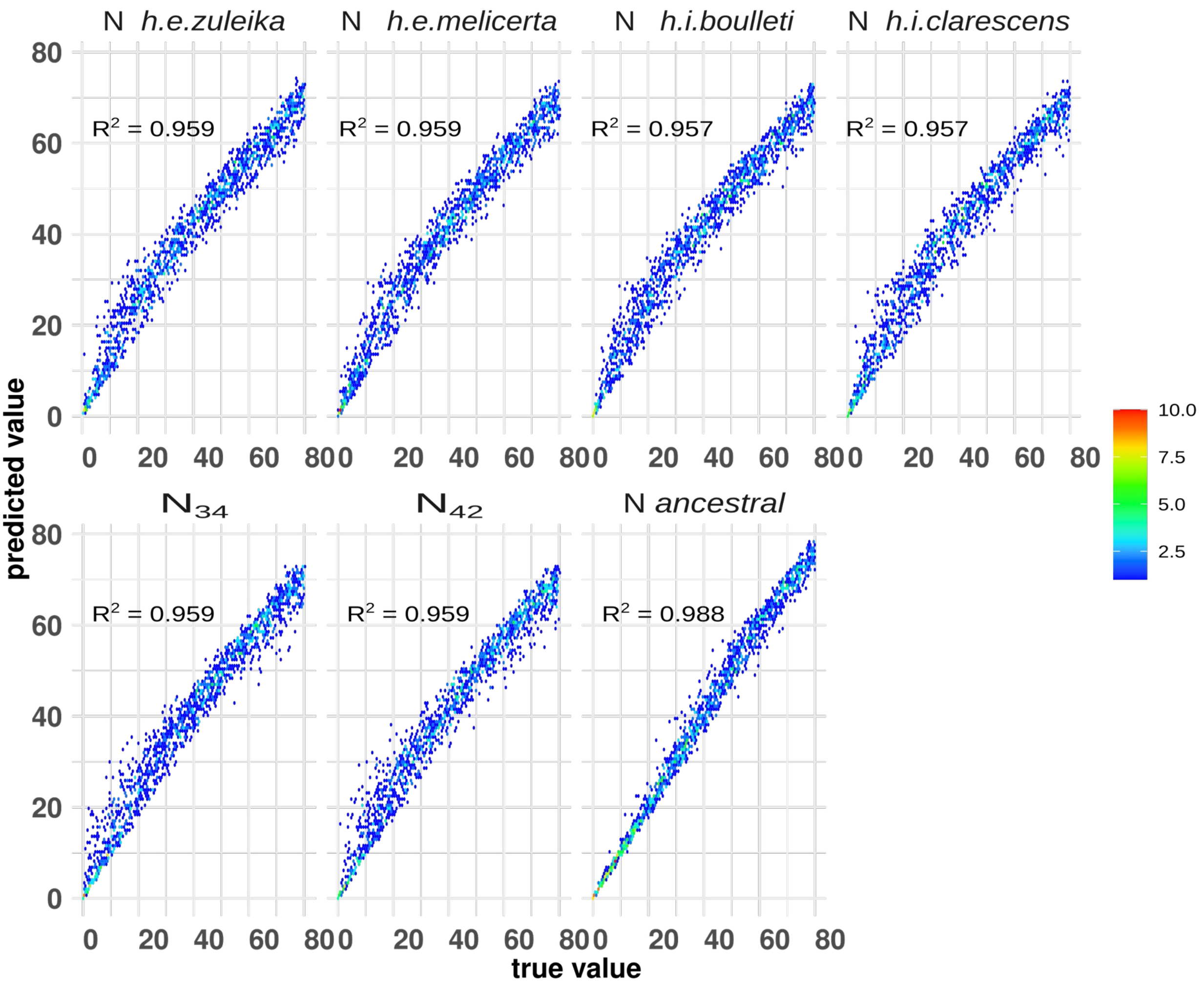
N h.e.zuleika
N h.e.meliçerta
N h.i.boulleti
N h.i.clarescens
$N_{34}$
$N_{42}$
N ancestral
R² = 0.959
R² = 0.959
R² = 0.957
R² = 0.957
R² = 0.959
R² = 0.959
R² = 0.988
predicted value
true value
10.0
7.5
5.0
2.5

B)

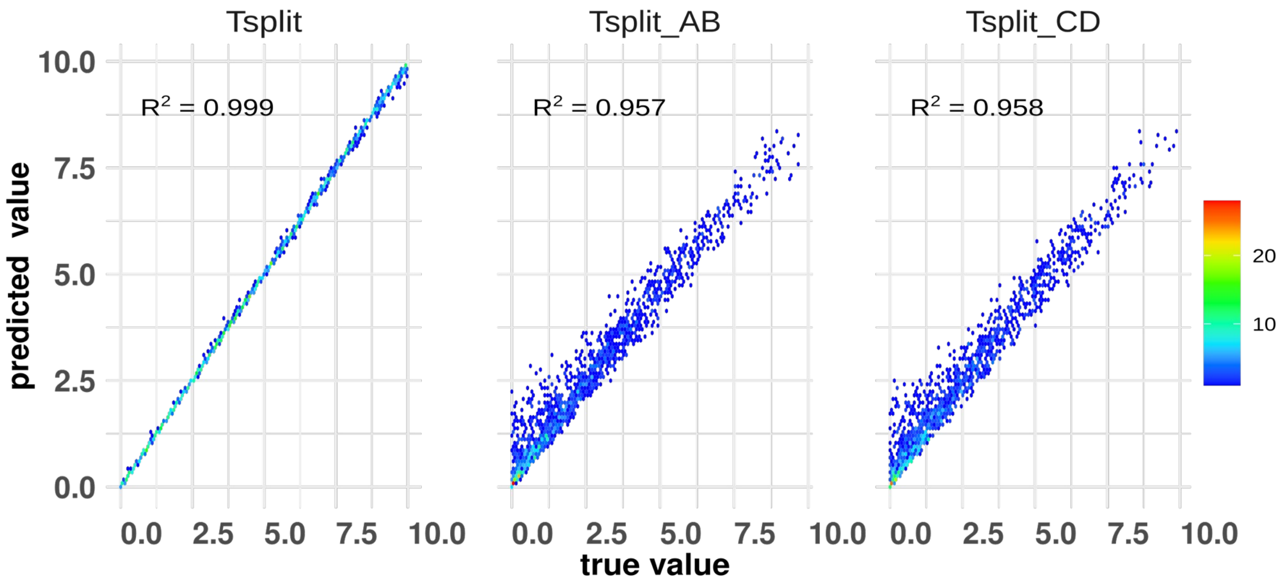
Tsplit
Tsplit_AB
Tsplit_CD
R² = 0.999
R² = 0.957
R² = 0.958
predicted value
true value
20
10

**C)**

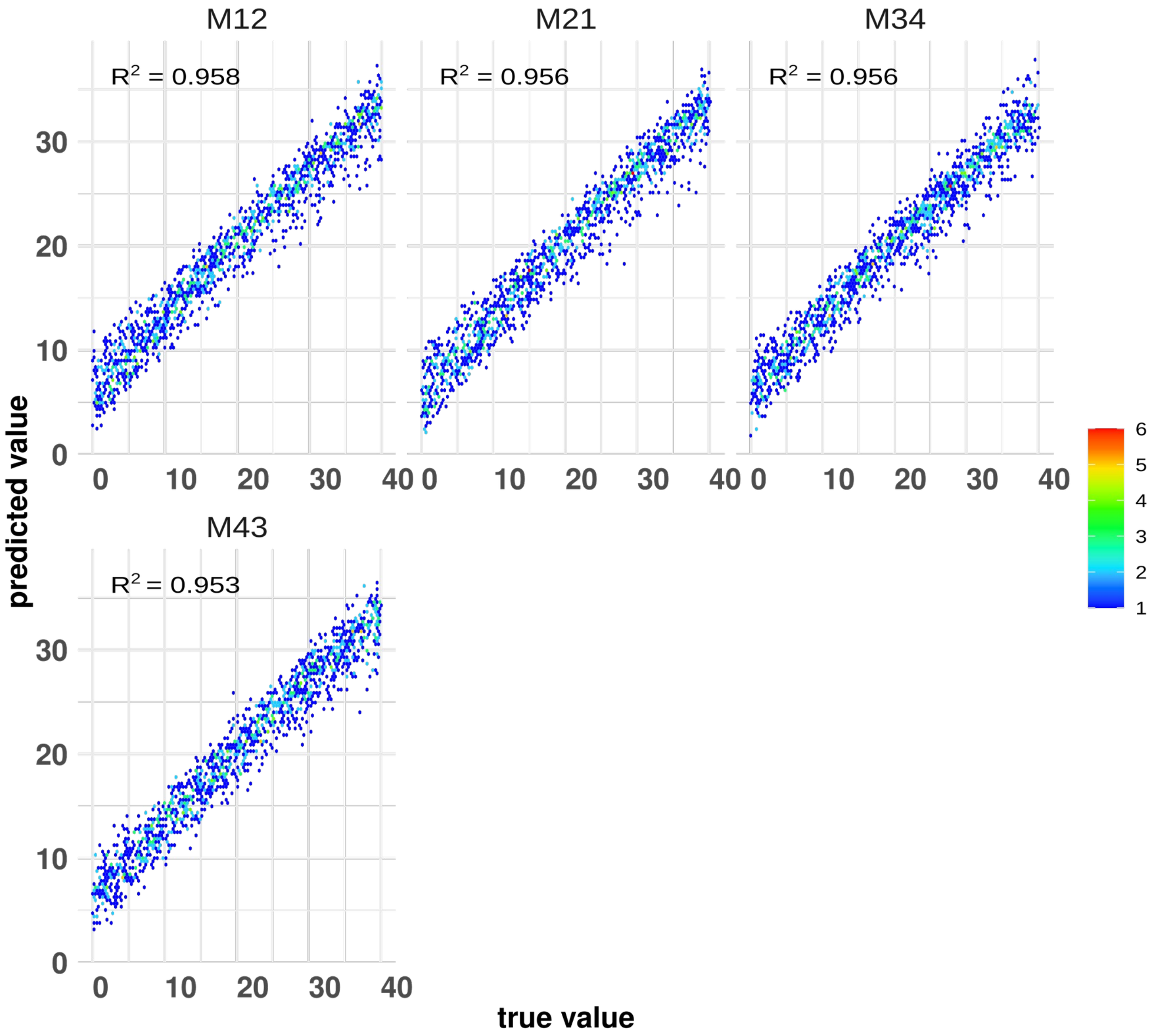

**Figure S08. ABC-RF prediction accuracy under the model of strict isolation (except within *hecale* and within *ismenius*).**

Showed are the correlations between each observed value from simulated data (i.e. true value x-axis) and the predicted value (y-axis) by the RF algorithm. Values provided in “coalescent units” based on 1,000 pseudo-observed datasets.

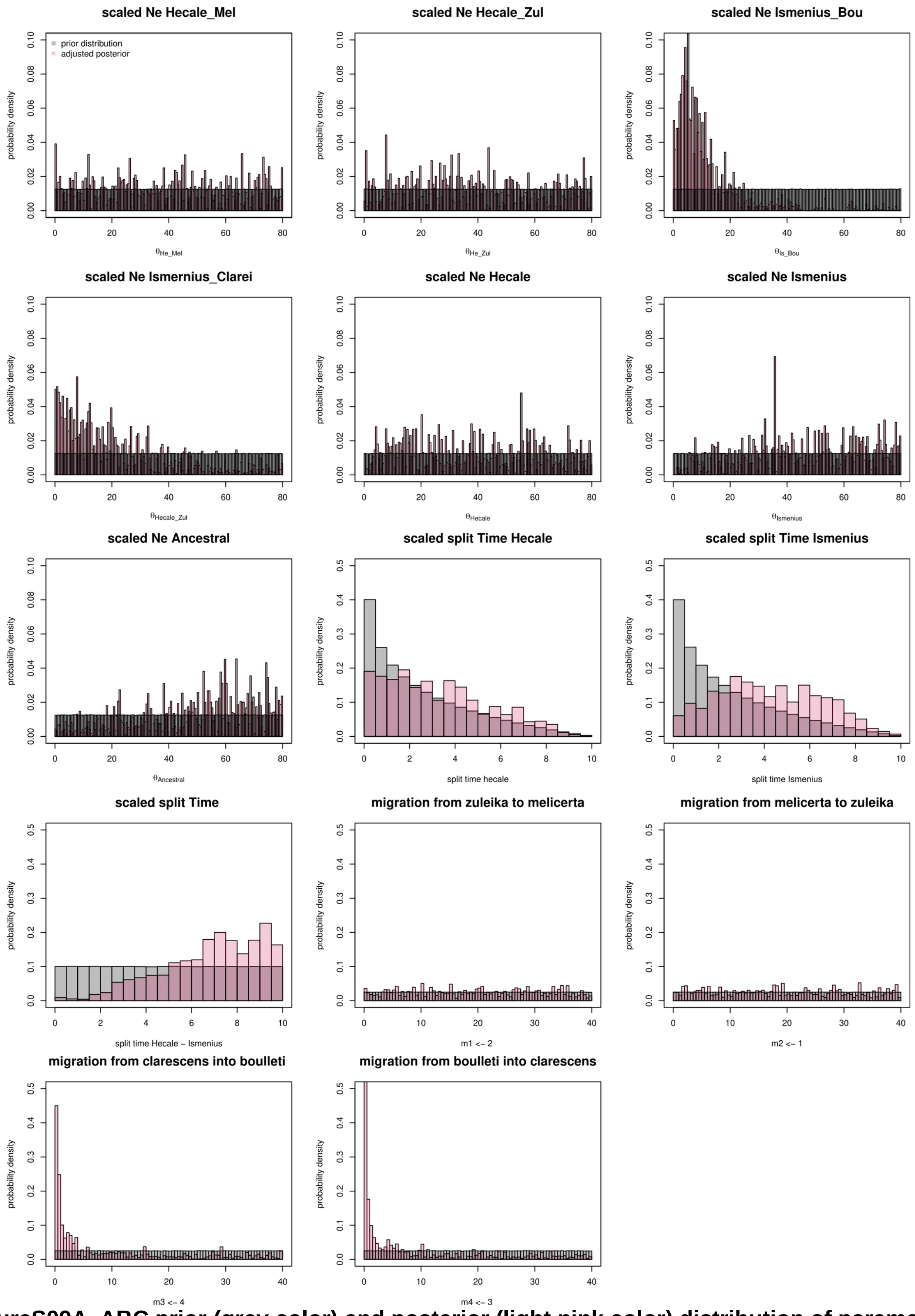

**FigureS09A. ABC prior (grey color) and posterior (light pink color) distribution of parameter estimates under the model of strict isolation among species.**

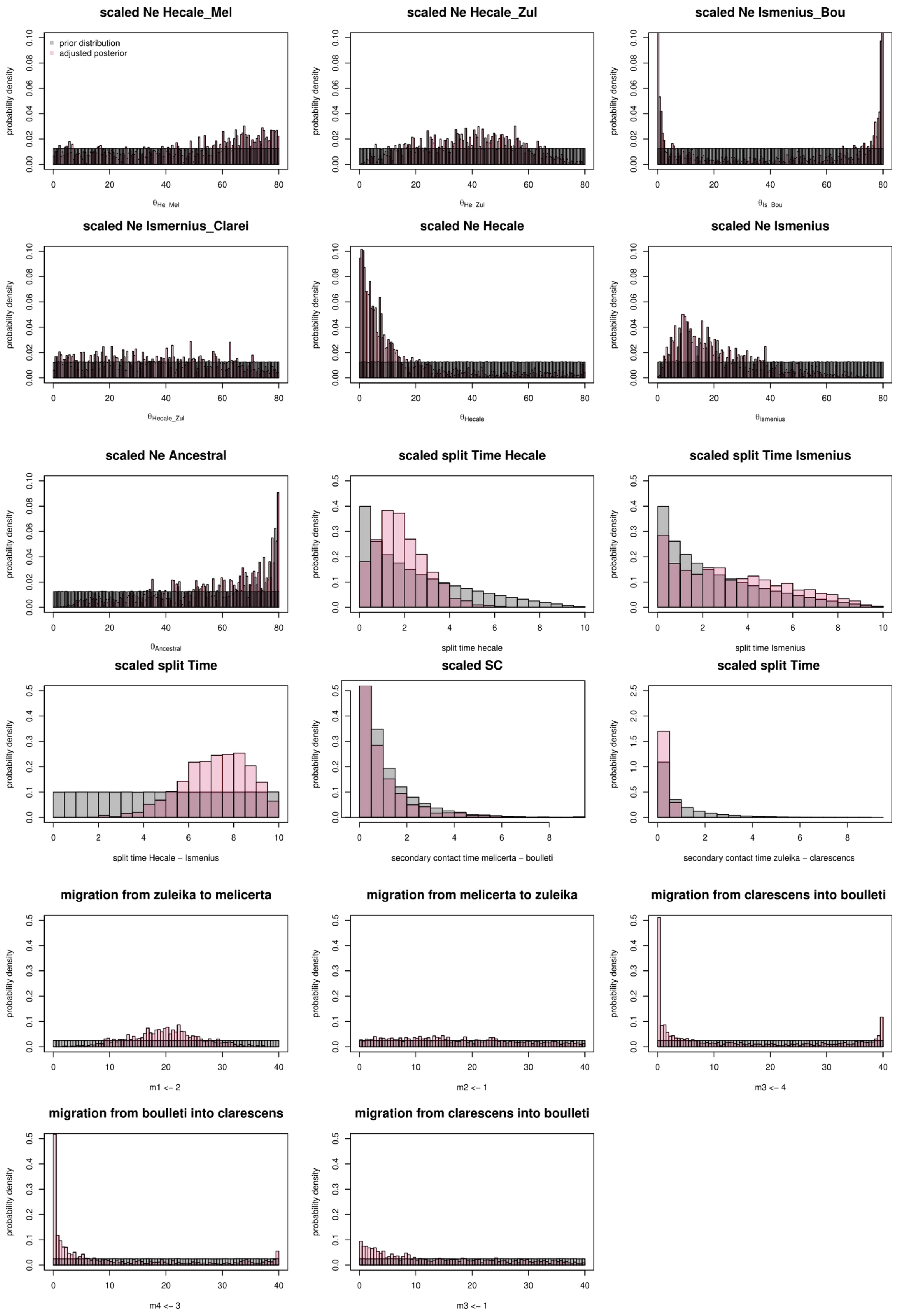

**Figure S09B. ABC (grey color) and posterior (light pink color) distribution of parameter estimates under the model of Secondary Contact among species.**

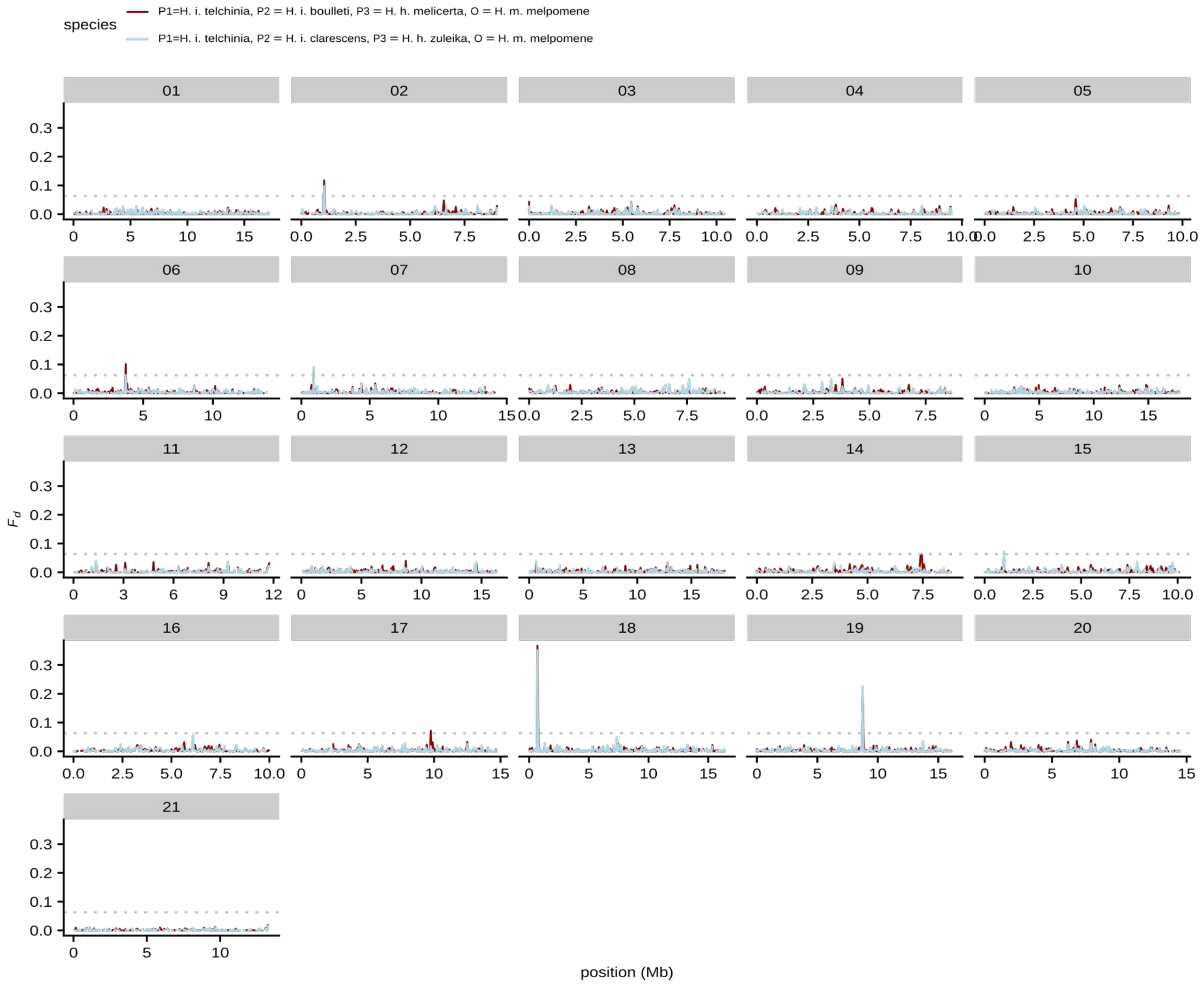

**Figure S10. ABBA-BABA landscape ($f_d$ values) along the genome considering *ismenius telchinia* as P1.**

Gene flow between *H. ismenius boulleti* (=P2) and *H. hecale melicerta (=P3) (co-mimics)* is displayed in red. Gene flow between *H. ismenius clarescens* (=P2) and *H. hecale zuleika (=P3) (non co-mimics)* is displayed in lightblue. Each chromosome is displayed separately with its number provided in the grey box. Results are summarised in 10kb windows with a 1000 bp step. The strongest signals are observed on CHR18 (around *optix*) and CHR19, and are shared in both directions.

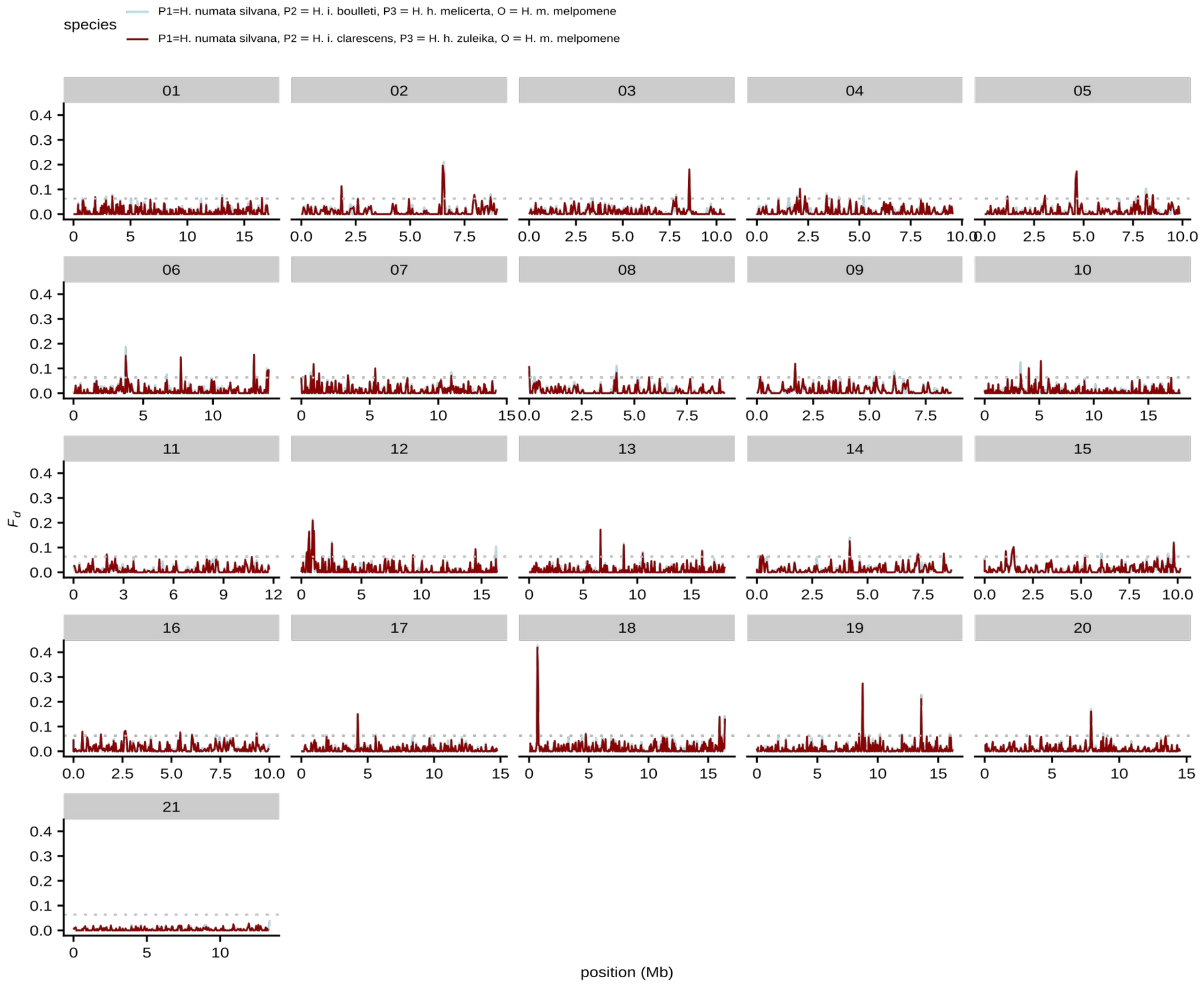

**Figure S11. ABBA-BABA landscape ($f_d$ values) along the genome considering *H. numata* as P1.**

Gene flow between *H. ismenius boulleti* (=P2) and *H. hecale melicerta (=P3) (co-mimics)* is displayed in red. Gene flow between *H. ismenius clarescens* (=P2) and *H. hecale zuleika (=P3) (non co-mimics)* is displayed in lightblue. Each chromosome is displayed separately with its number provided in the grey box. Results are summarised in 10kb windows with a 1000 bp step. The strongest signals are observed on CHR18 (around *optix*) and CHR19, and are shared in both directions.

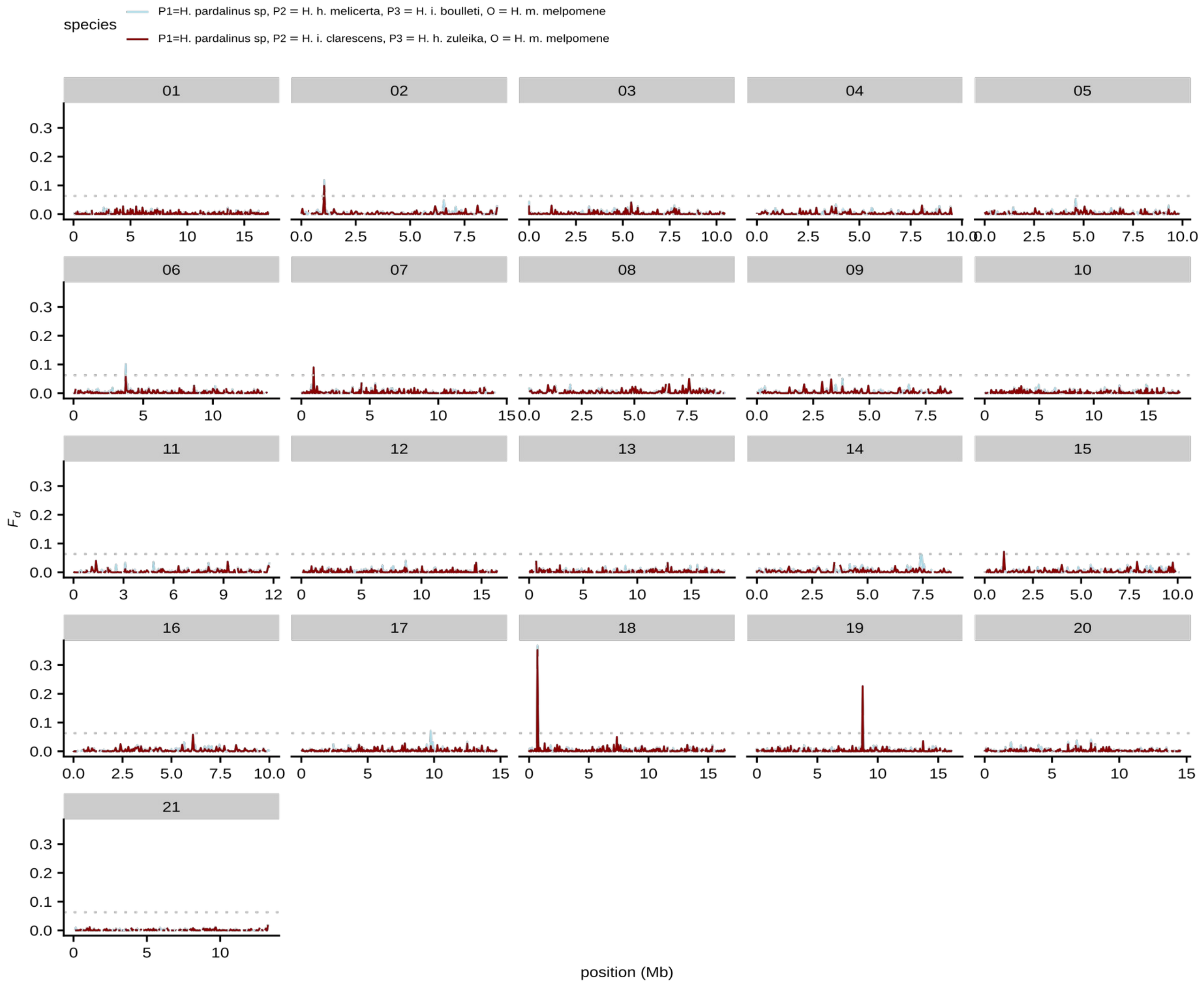

**Figure S12. ABBA-BABA landscape ($f_d$ values) along the genome considering *H. pardalinus* as P1.**

Gene flow between *H. ismenius boulleti* (=P2) and *H. hecale melicerta (=P3) (co-mimics)* is displayed in red. Gene flow between *H. ismenius clarescens* (=P2) and *H. hecale zuleika (=P3) (non co-mimics)* is displayed in lightblue. Each chromosome is displayed separately with its number provided in the grey box. Results are summarised in 10kb windows with a 1000 bp step. The strongest signals are observed on CHR18 (around *optix*) and CHR19, and are shared in both directions.

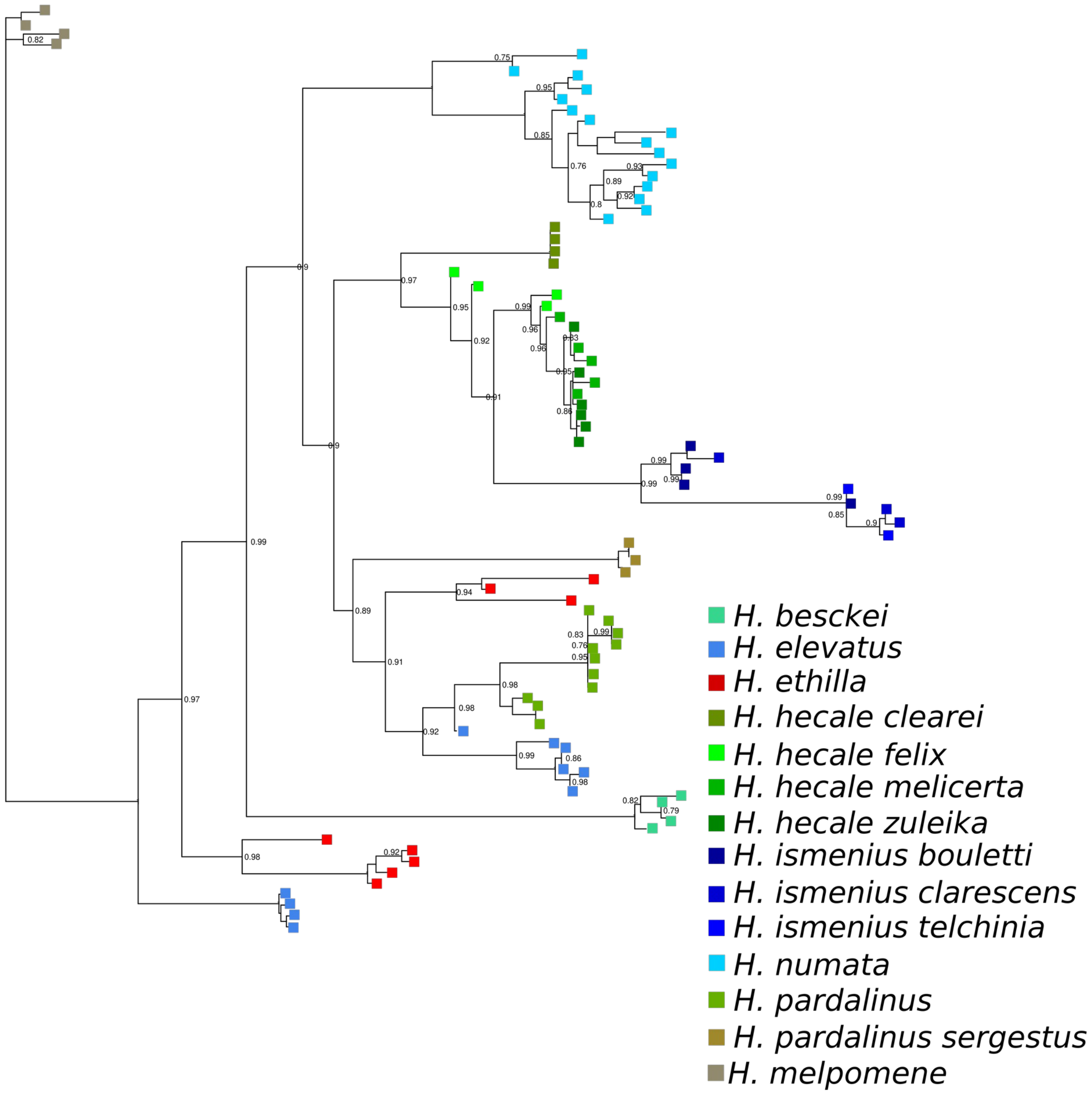

**Figure S13. Phylogenetic tree inferred at the four locally introgressed genes in *H. ismenius* on chromosome 19.**

Phylogenetic tree obtained with Raxml. Bootstrap support obtained with 100 replicate trees.

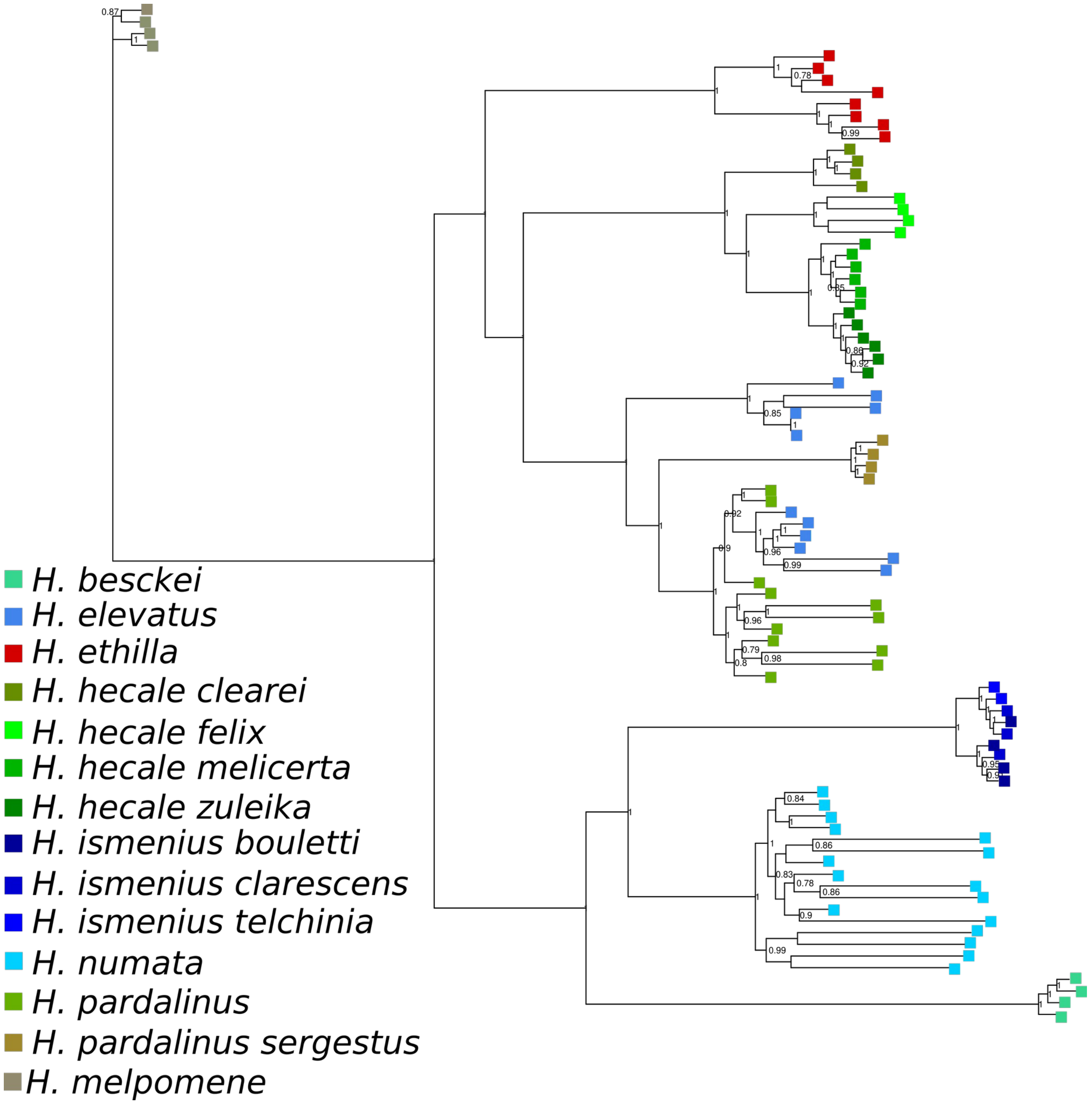

**Figure S14. Phylogenetic tree inferred using the sex chromosome.**

Phylogenetic tree obtained with Raxml. Bootstrap support obtained with 400 replicate trees.

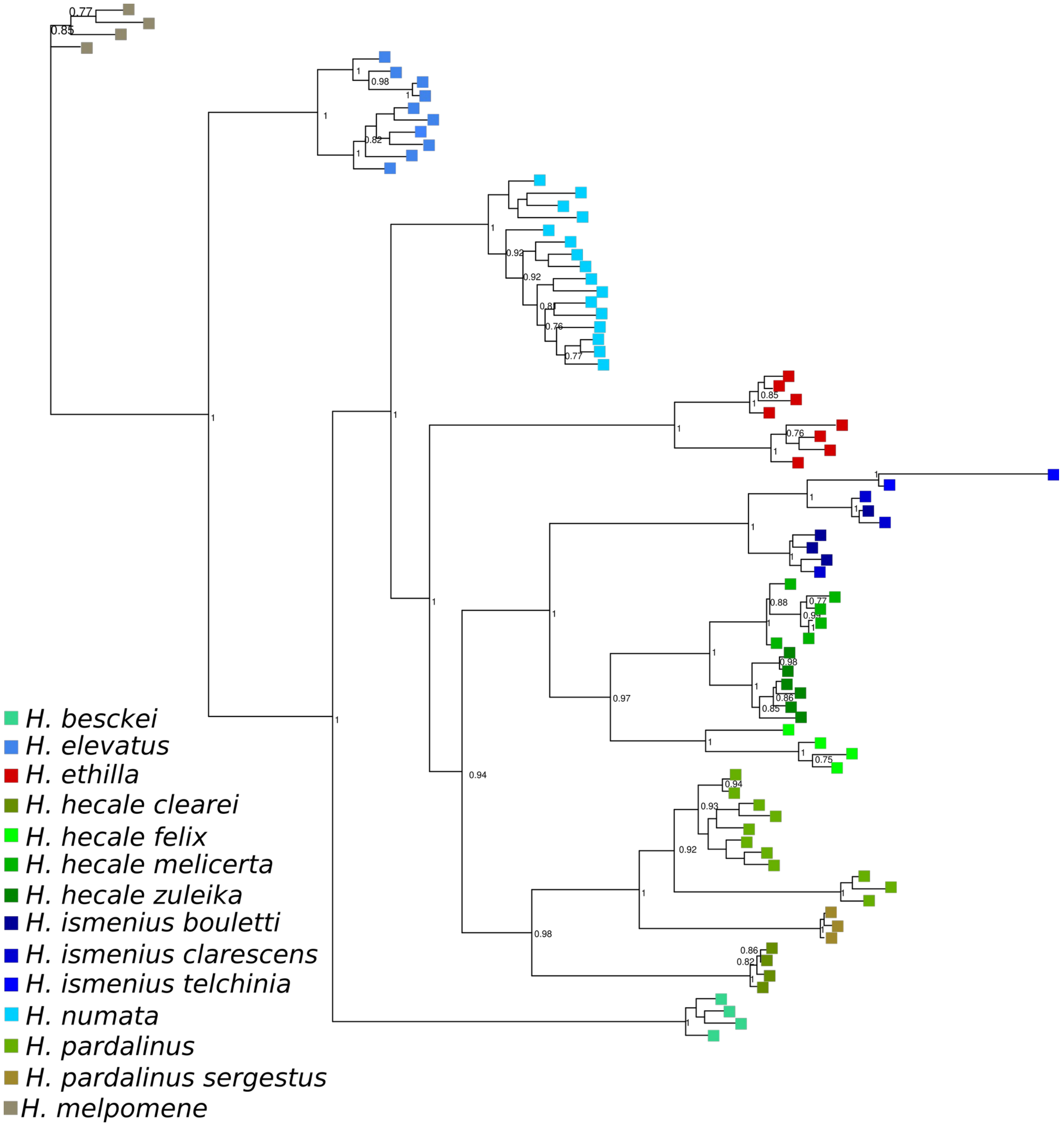

**Figure S15. Phylogenetic tree inferred using a 100 kb region around *optix* gene on chromosome 18.**

Phylogenetic tree obtained with Raxml. Bootstrap support obtained with 100 replicate trees.

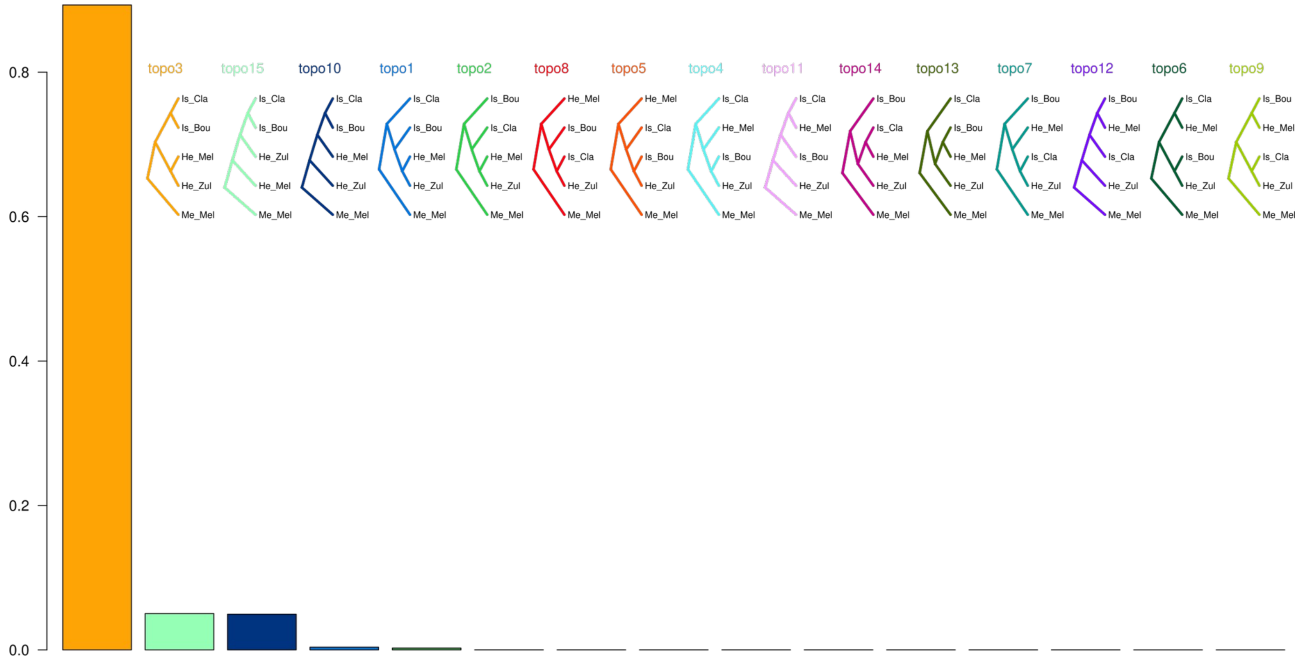

**Figure S16. Shape of the 15 topologies being tested and weight support for each 15 topologies.**

Topo3: Species tree (congruent phylogeny)
Topo10 & 15: Partial species tree where *hecale* is paraphyletic
Topo1 & 2: Partial species tree where *ismenius* is paraphyletic
Topo8 & 7: Gene flow between non-comimics (higher than gene flow between co-mimics)
Topo12 & 13: Gene flow due to mimicry (higher than gene flow due to non-mimicry)
Topo9: Gene flow in sympatry (i.e. between co-mimics on one side and non-comimics on the other)
Topo4,5,6,11,14: Incongruent topologies
Each topology is coloured differently. The species tree received the highest support.

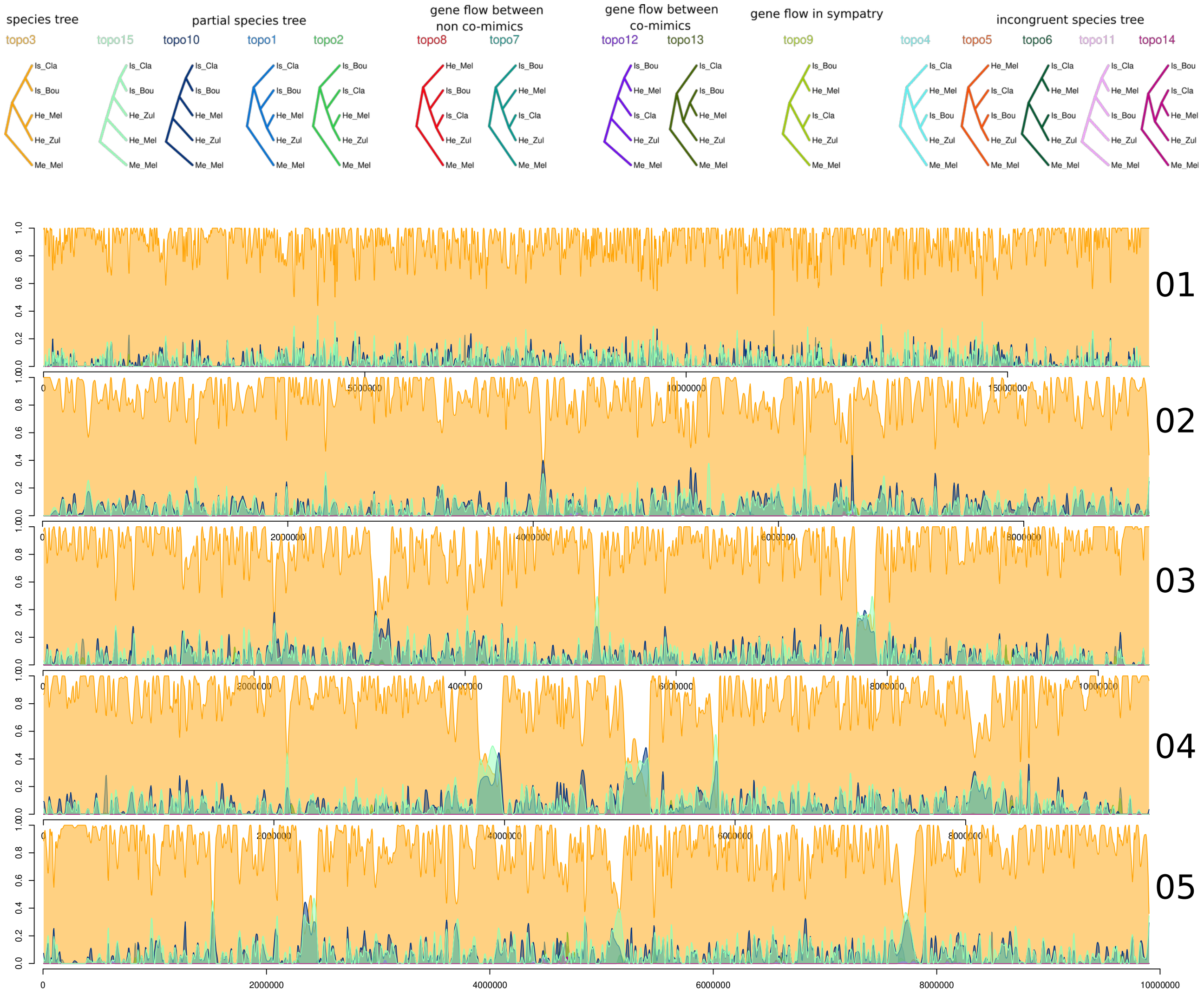

**Figure S17A. Twisst variation in topology support along the genome for chromosomes 01 to 05.**

Each colour represents a topology and its support is depicted along the y-axis.

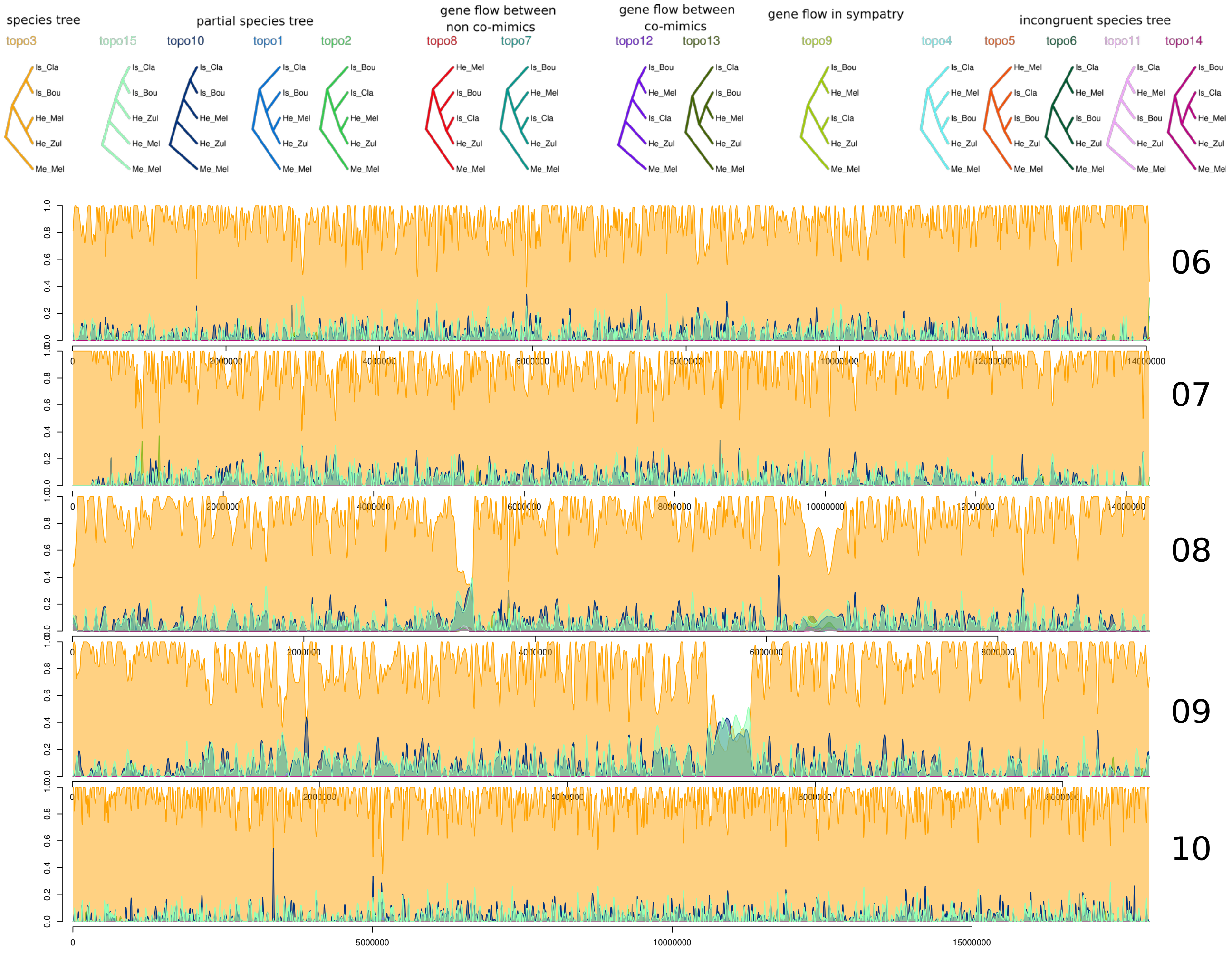

**Figure S17B. Twisst variation in topology support along the genome for chromosomes 06 to 10.**

Each colour represents a topology and its support is depicted along the y-axis.

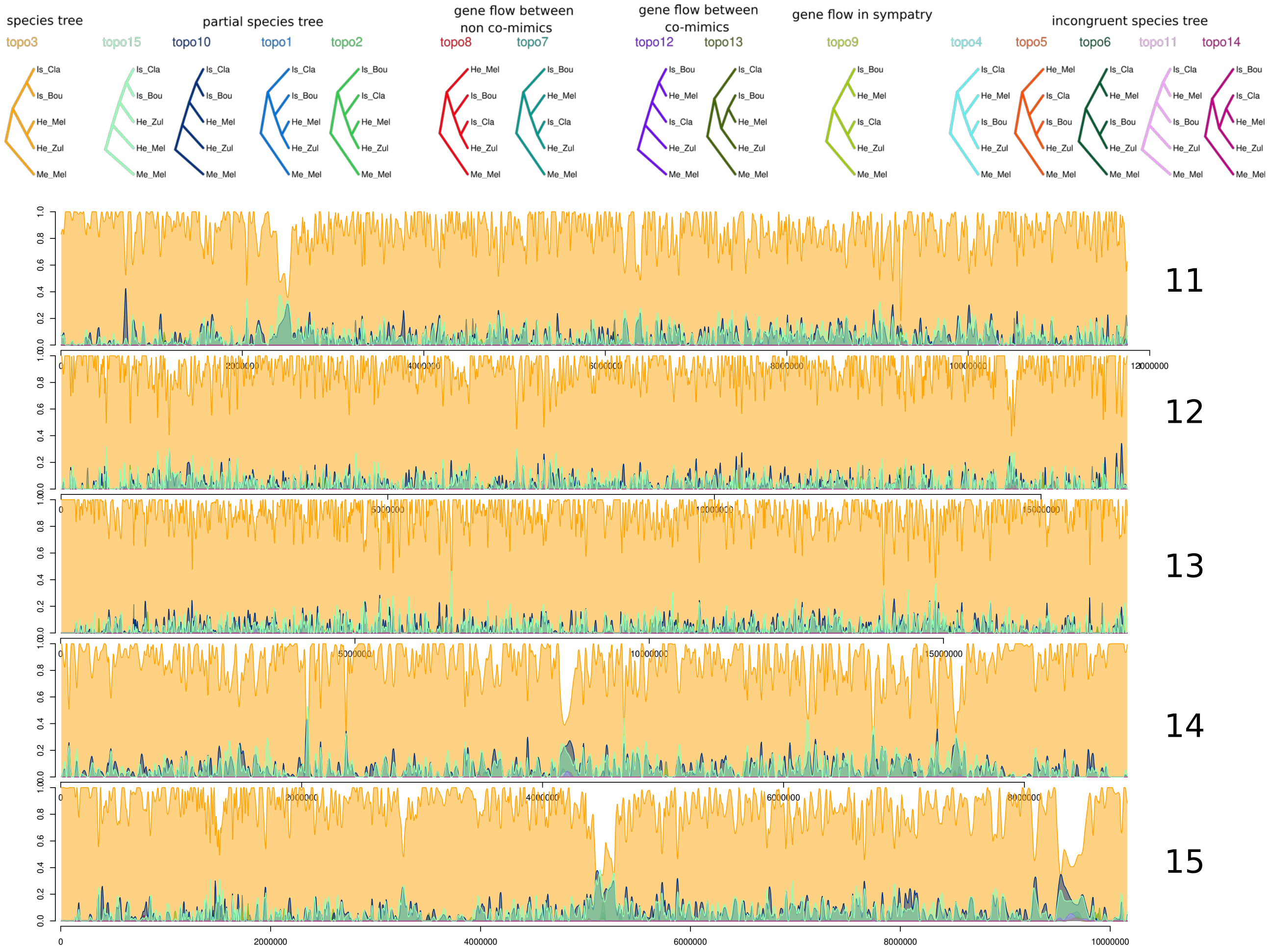

**Figure S17C. Twisst variation in topology support along the genome for chromosomes 11 to 15.**

Each colour represents a topology and its support is depicted along the y-axis.

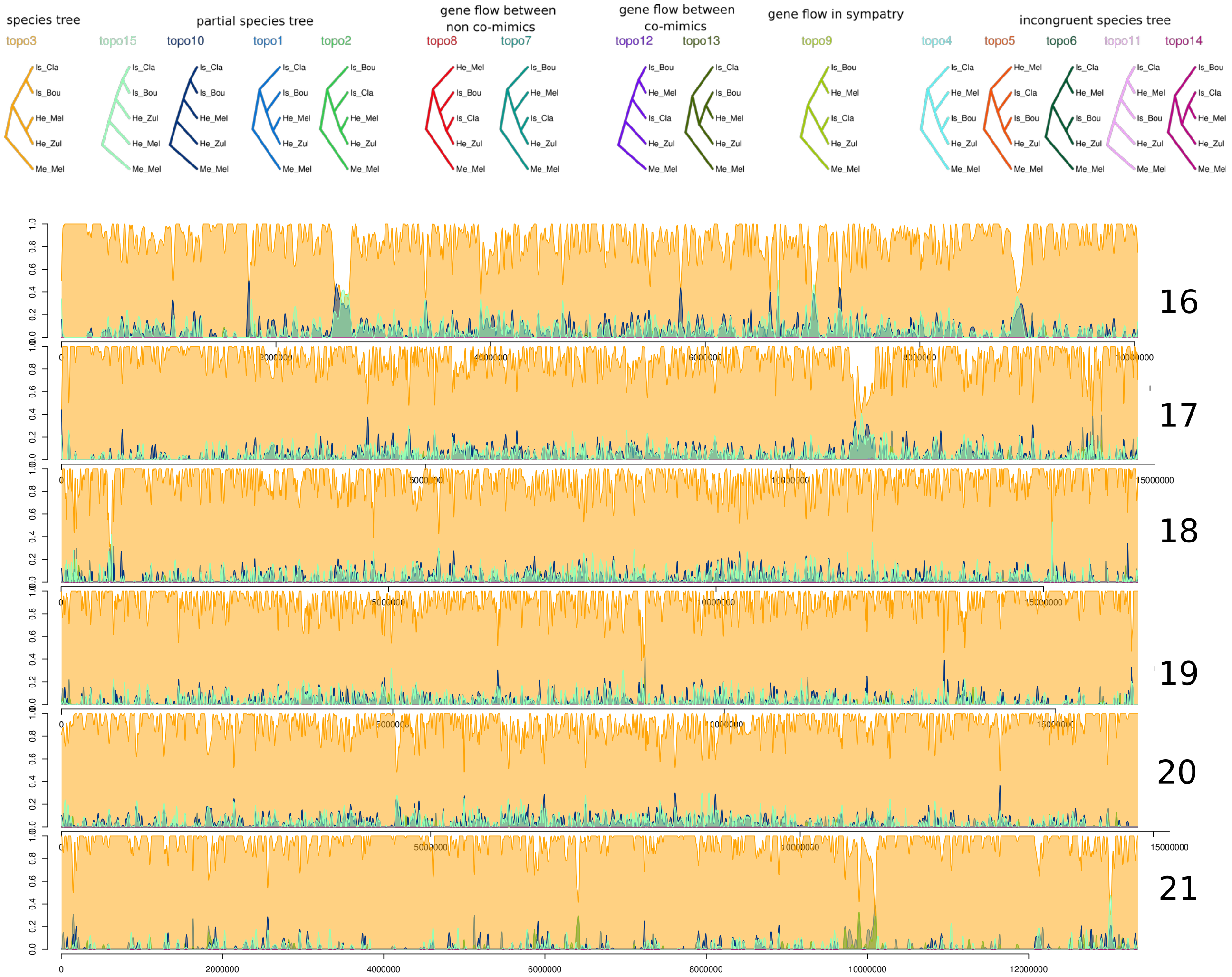

**Figure S17D. Twisst variation in topology support along the genome for chromosomes 16 to 21.**

Each colour represents a topology and its support is depicted along the y-axis.

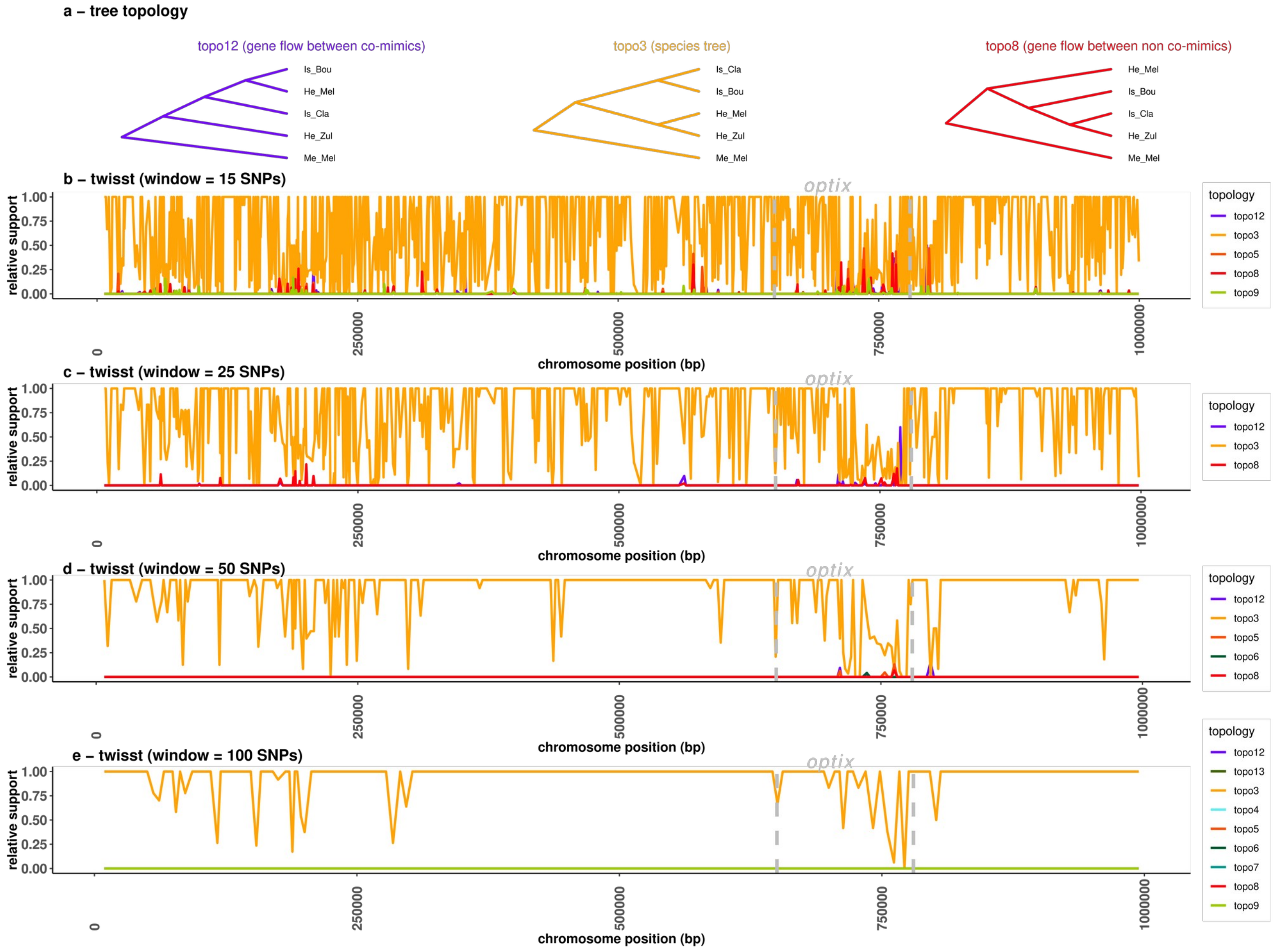

**Figure S18. Effect of window size on detection of introgression signals around *optix* on chromosome 18.**

Topology 3 corresponds to the species tree. Topology 12 and 13 are expected under gene flow between co-mimics and topology 8 is expected under gene flow between non co-mimics. Topology 9 is expected under introgression due to sympatry (i.e. introgression between both co-mimics and non co-mimics).

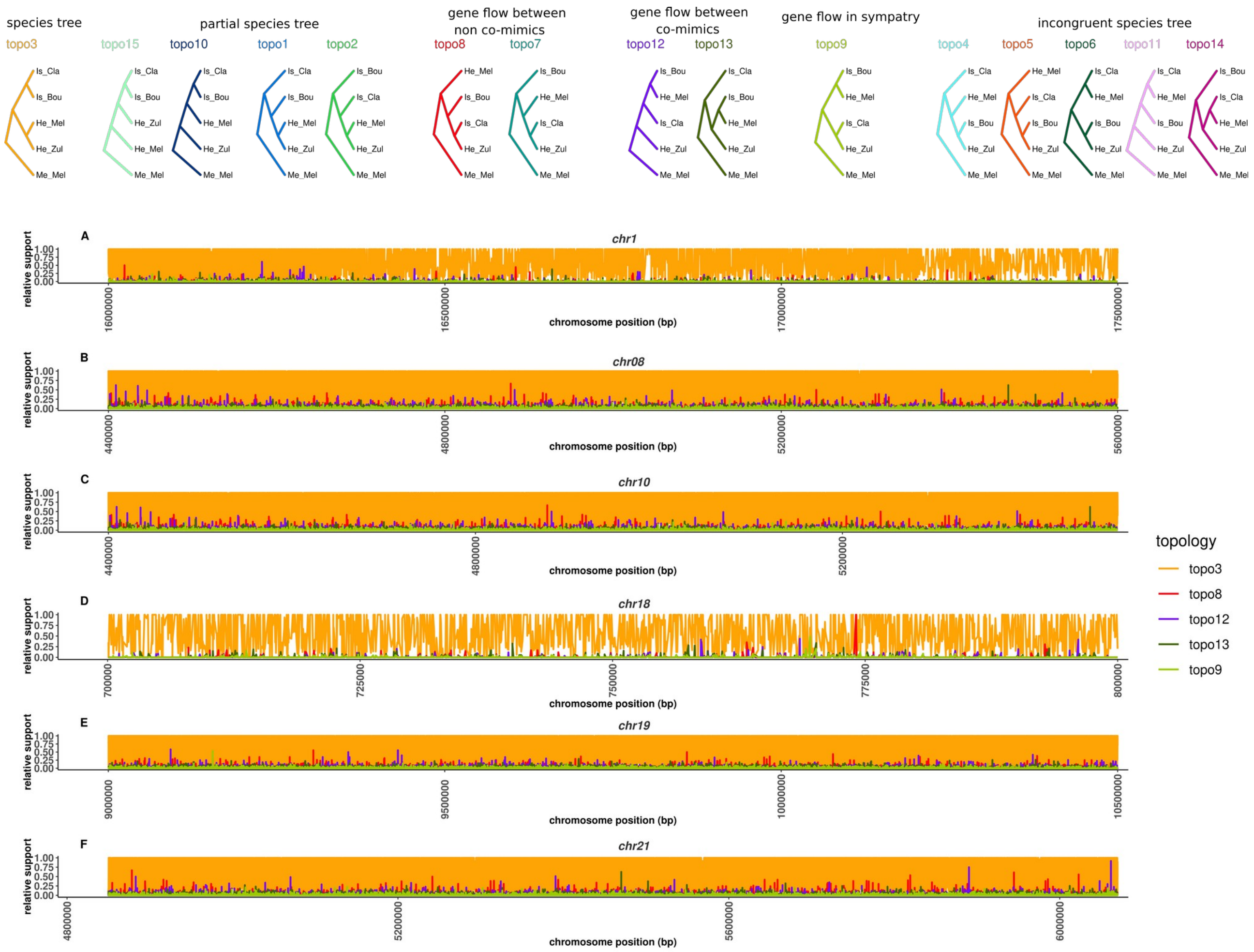

**Figure S19: Introgressed windows have been broken down by recombination into small and localised windows.**

Figure displaying the most salient introgression peaks along the genome. Topology 3 corresponds to the species tree. Topology 12 and 13 are expected under gene flow between co-mimics and topology 8 is expected under gene flow between non co-mimics. Topology 9 is expected under introgression due to sympatry (i.e. introgression between both co-mimics and non co-mimics).

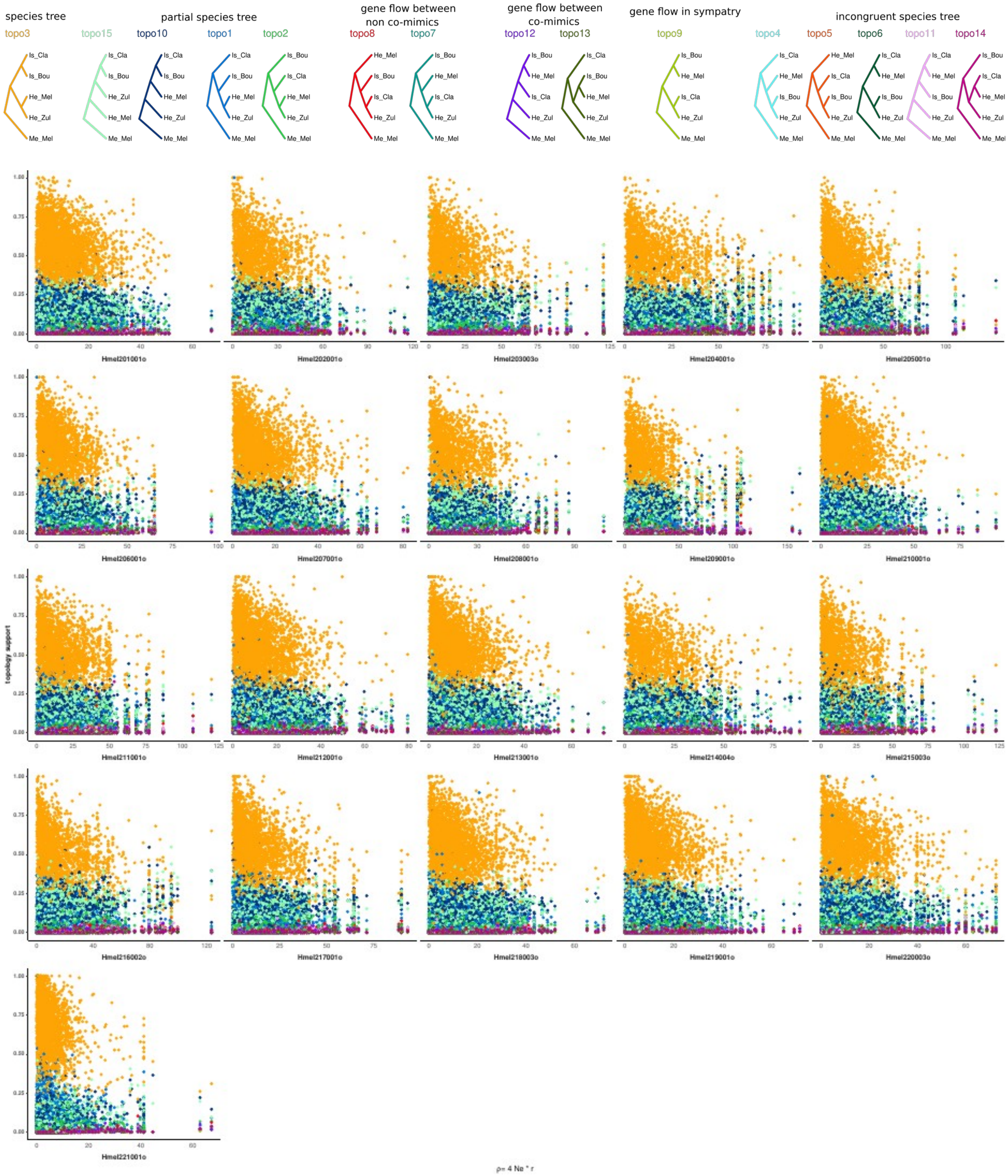

**Figure S20. Support for the species tree decreases with increasing recombination.**

Showed is the relative topology support for each window (Twisst window size = 10 SNPs) on the y-axis as a function of population scale recombination rate (x-axis) estimated with LDhat and averaged on 5 kb windows.

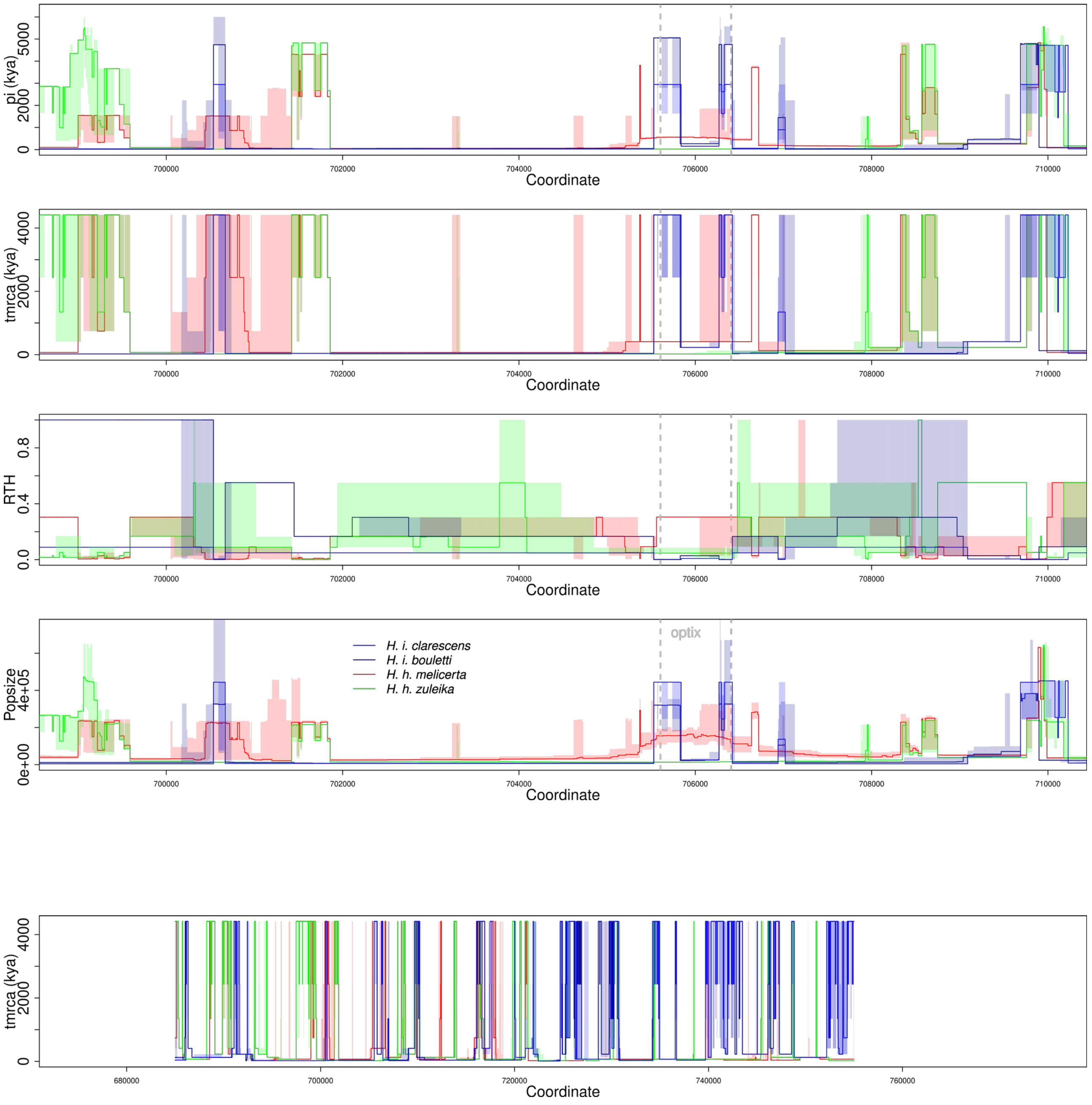

**Figure S21a. Inference of the ancestral recombination graph (ARG) from ARGWeaver for each species along the region surronding the *Optix* gene.**

The dotted grey line displays the limits of optix.
The bottom panel display local variation in the TMRCA in a broader context along the chromosome 18.
TMRCA (Time to the Most Recent Common Ancestor) and RTH (relative TMRCA half-time) along part of the chromosome 18, including the gene *Optix* (position = 704 - 705 Kb). The RTH describes the minimum time for half of the lineages to coalesce, normalised by the TMRCA.

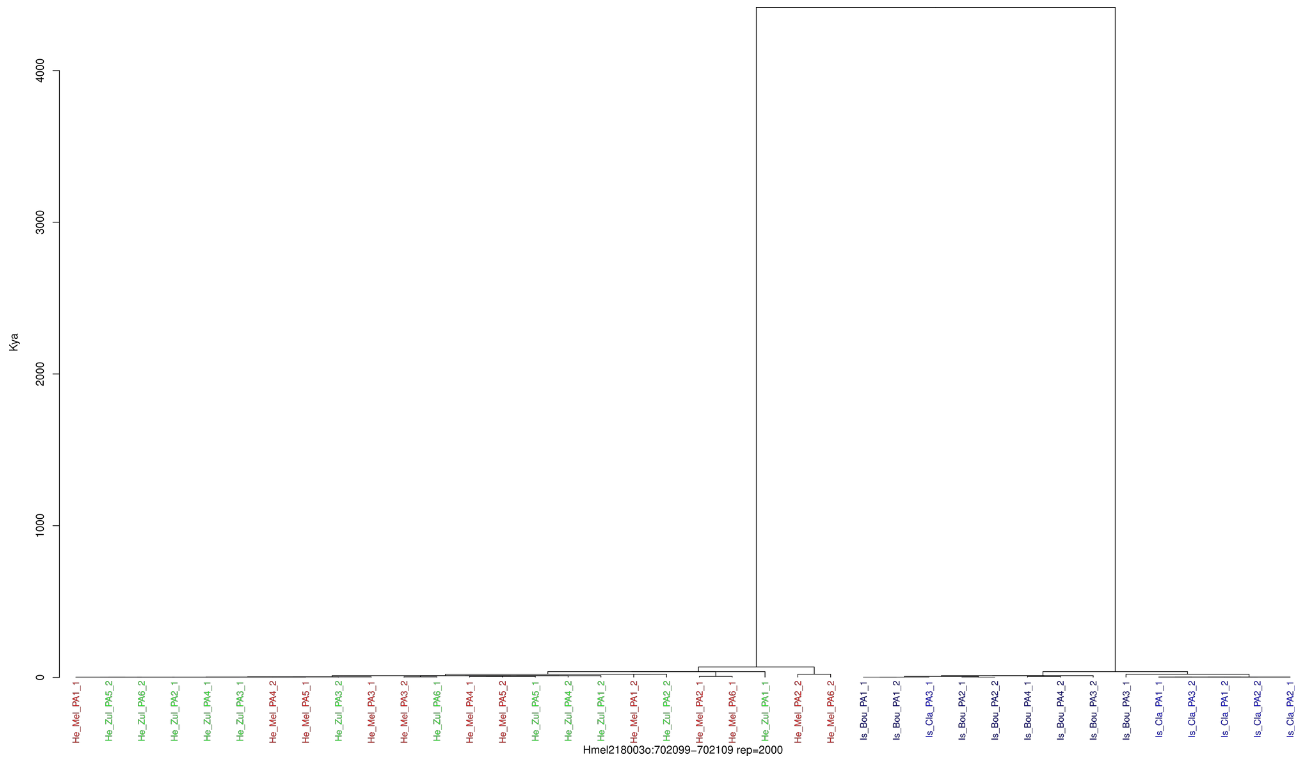

**Figure S21b. Examples of local trees infered from the ancestral recombination graph (ARG) using ARGWeaver for each species along the region surrounding the *Optix* gene.**

Example of the most commonly observed tree with a deep TMRCA, compatible with the inffered species tree and divergence time from our ABC analyses.

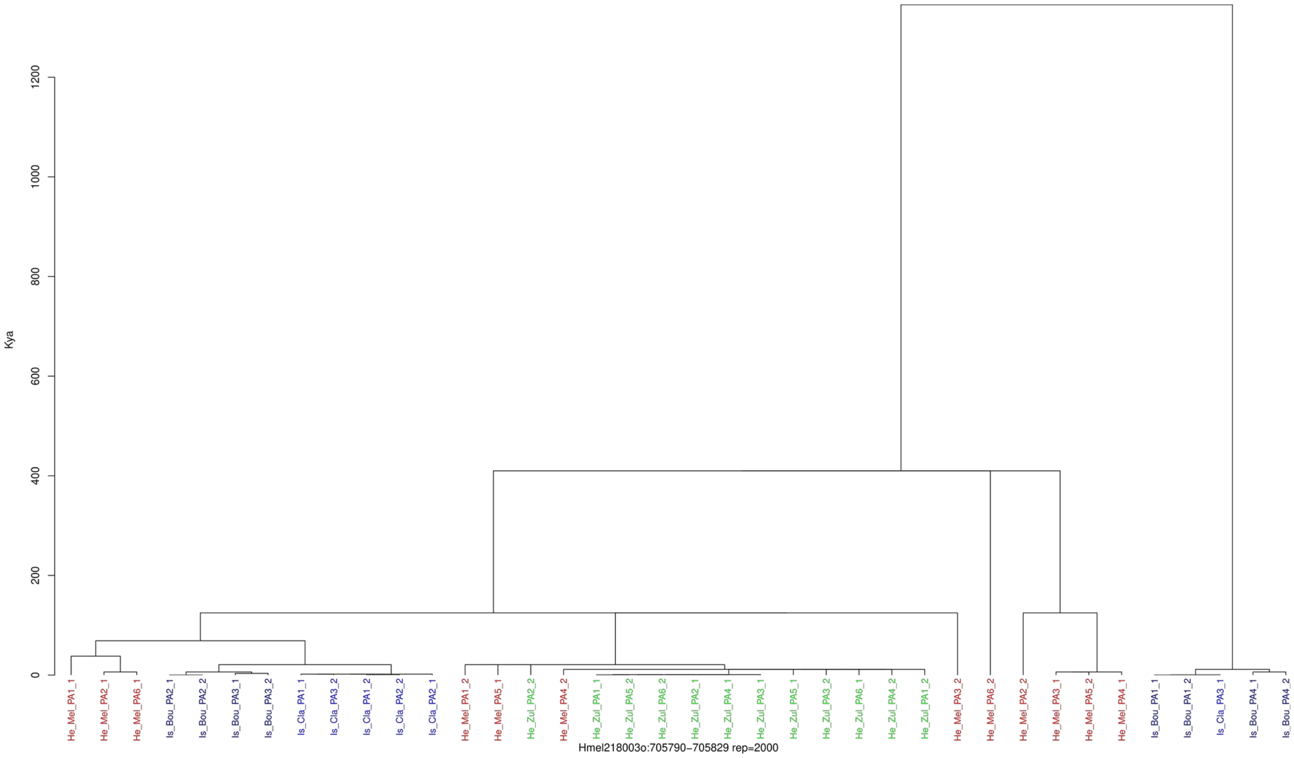

Example tree with a recent TMRCA (within the optix gene) suggestive of a recent divergence or gene flow within *Optix* gene

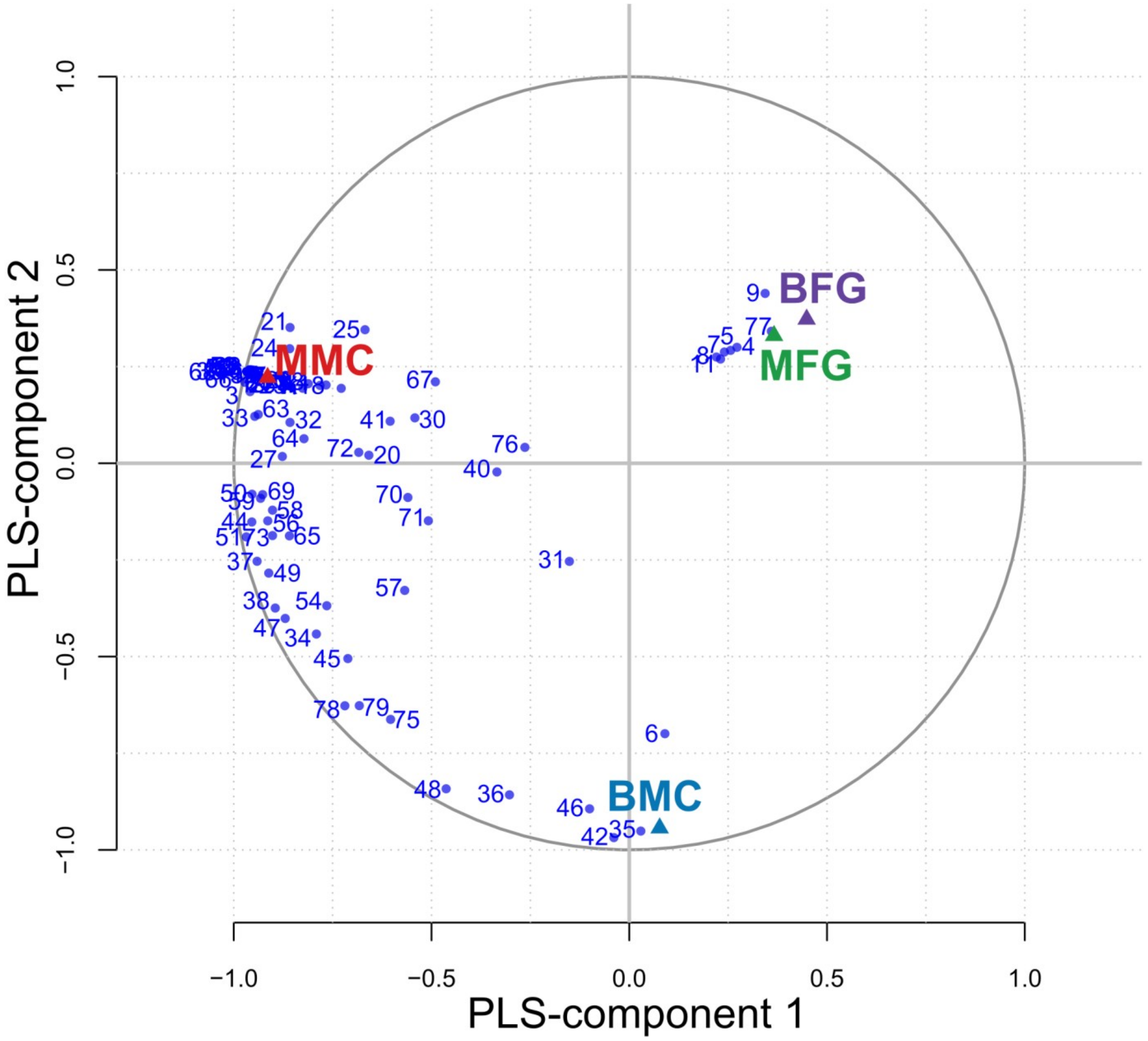

**Figure S22. Comparison of chemical blends of abdominal glands.**

Partial least squares discriminant analysis (PLS-DA) based on comparisons of the chemical blends composition of abdominal glands. A correlation circle is plotted between the compounds (variables; blue dots and labels) and the components. The variables being indicative of the groups are strongly correlated with the groups being separated and close to the concentric circle of radius 1. See Tables S14 and S17 for details about the compounds

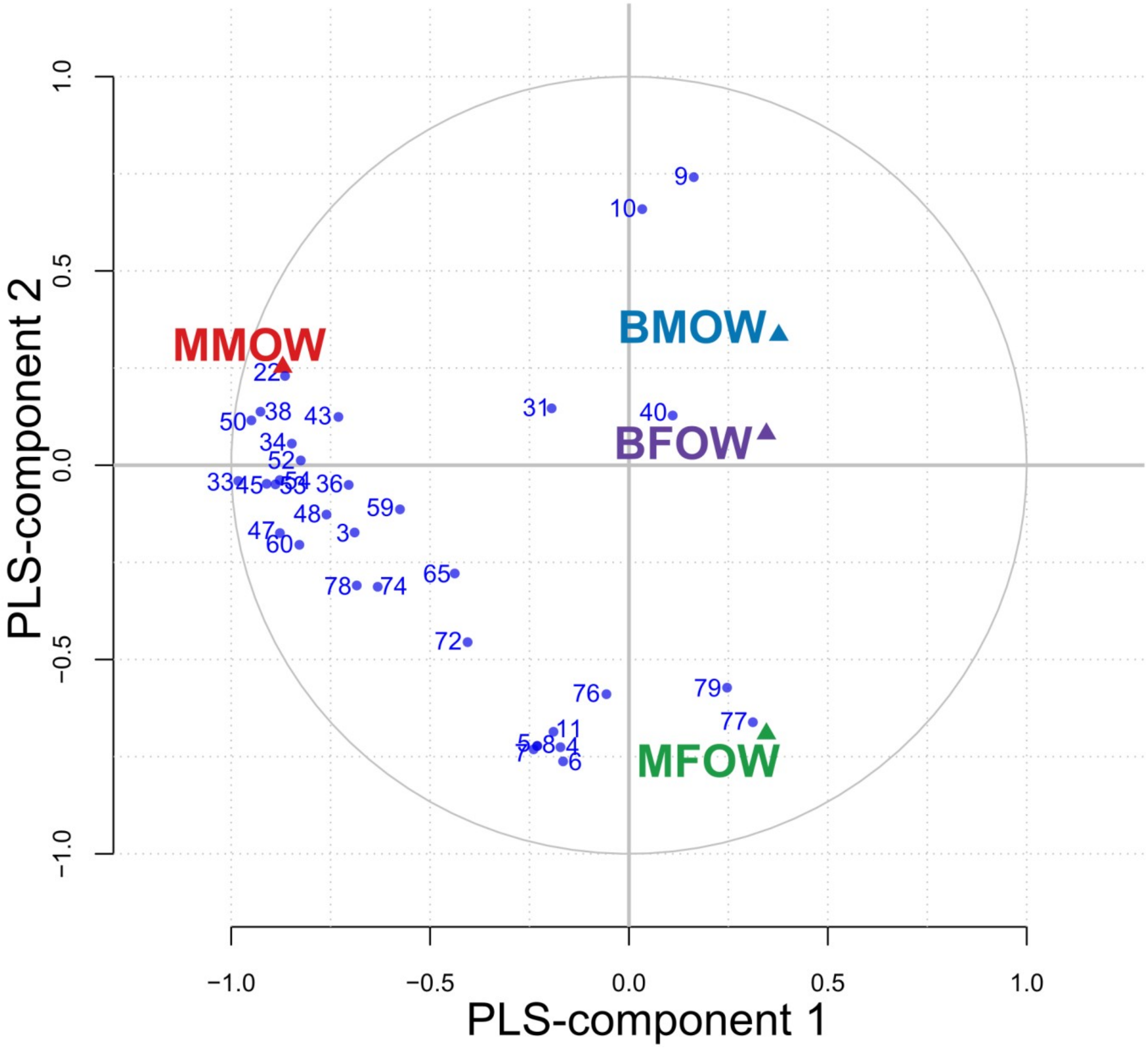

**Figure S23. Comparisons of chemical blends of the wings among species.**

Partial least squares discriminant analysis (PLS-DA) based on comparisons of the chemical blends composition of the wings for *H. h. melicerta* males (MMOW) and females (MFOW), and *H. i. boulleti* males (BMOW) and females (BFOW). A correlation circle is plotted between the compounds (variables; blue dots and labels) and the components. The variables being indicative of the groups are strongly correlated with the groups being separated and close to the concentric circle of radius 1. See Tables S15 and S17 for details about the compounds.

**Table S01. Details of sampling design, including country of sampling, accession numbers and sample mean depth of sequencing.**

| SampleID | Taxon | Country | Accession | MeanDepth |
|---|---|---|---|---|
| He Cle VE1 | *H. hecale clearei* | Venezuela | NA | 26.423608 |
| He Cle VE2 | *H. hecale clearei* | Venezuela | NA | 28.182284 |
| He Cle VE3 | *H. hecale clearei* | Venezuela | NA | 22.782875 |
| He Cle VE4 | *H. hecale clearei* | Venezuela | NA | 31.84672 |
| He Fel PR4 | *H. hecale felix* | Peru | ERP002440 | 33.979015 |
| He Hec R1 | *H. hecale felix* | Peru | ERP009041 | 24.100849 |
| He Hec R2 | *H. hecale felix* | Peru | ERP009041 | 25.337944 |
| He Hec R3 | *H. hecale felix* | Peru | ERP009041 | 26.592065 |
| He Mel PA1 | *H. hecale melicerta* | Panama | SRP145614 | 18.82234 |
| He Mel PA2 | *H. hecale melicerta* | Panama | NA | 18.340344 |
| He Mel PA3 | *H. hecale melicerta* | Panama | NA | 13.345133 |
| He Mel PA4 | *H. hecale melicerta* | Panama | NA | 18.643497 |
| He Mel PA5 | *H. hecale melicerta* | Panama | NA | 22.939988 |
| He Mel PA6 | *H. hecale melicerta* | Panama | NA | 18.894109 |
| He Zul PA1 | *H. hecale zuleika* | Panama | SRP145614 | 23.594665 |
| He Zul PA2 | *H. hecale zuleika* | Panama | NA | 14.521932 |
| He Zul PA3 | *H. hecale zuleika* | Panama | NA | 14.494293 |
| He Zul PA4 | *H. hecale zuleika* | Panama | NA | 26.303984 |
| He Zul PA5 | *H. hecale zuleika* | Panama | NA | 20.927219 |
| He Zul PA6 | *H. hecale zuleika* | Panama | NA | 20.328594 |
| Is Bou PA1 | *H. ismenius boulleti* | Panama | SRP145614 | 18.018185 |
| Is Bou PA2 | *H. ismenius boulleti* | Panama | NA | 26.064319 |
| Is Bou PA3 | *H. ismenius boulleti* | Panama | NA | 14.250511 |
| Is Bou PA4 | *H. ismenius boulleti* | Panama | NA | 20.284426 |
| Is Cla PA1 | *H. ismenius clarescens* | Panama | NA | 16.820047 |
| Is Cla PA2 | *H. ismenius clarescens* | Panama | NA | 19.316856 |
| Is Cla PA3 | *H. ismenius clarescens* | Panama | NA | 24.659618 |
| Is Tel ME1 | *H. ismenius telchinia* | Panama | SRP145614 | 15.47225 |
| Is Tel PA1 | *H. ismenius telchinia* | Panama | SRP145614 | 28.654071 |
| Me Agl PR1 | *H. melpomene aglaope* | Peru | ERP002440 | 27.820661 |
| Me Agl PR2 | *H. melpomene aglaope* | Peru | ERP002440 | 30.949998 |
| Me Agl PR3 | *H. melpomene aglaope* | Peru | ERP002440 | 34.701024 |
| Me Agl PR4 | *H. melpomene aglaope* | Peru | ERP002440 | 29.137085 |
| Me Ama PR1 | *H. melpomene amaryllis* | Peru | ERP002440 | 25.612402 |
| Me Ama PR2 | *H. melpomene amaryllis* | Peru | ERP002440 | 33.86252 |
| Me Ama PR3 | *H. melpomene amaryllis* | Peru | ERP002440 | 40.756403 |
| Me Ama PR4 | *H. melpomene amaryllis* | Peru | ERP002440 | 42.131738 |

**Table S01 *(continuation)***

| SampleID | Taxon | Country | Accession | MeanDepth |
|---|---|---|---|---|
| Me Mel FG1 | *H. melpomene melpomene* | Guiana | ERP002440 | 29.090925 |
| Me Mel FG2 | *H. melpomene melpomene* | Guiana | ERP002440 | 19.921848 |
| Me Mel FG3 | *H. melpomene melpomene* | Guiana | ERP002440 | 18.828091 |
| Me Mel FG4 | *H. melpomene melpomene* | Guiana | ERP002440 | 28.390443 |
| Nu Sil FG2 | *H. numata silvana* | Guiana | ERP123741 | 24.682775 |
| Nu Sil FG3 | *H. numata silvana* | Guiana | ERP123741 | 40.09978 |
| Nu Sil PR1 | *H. numata silvana* | Peru | ERP014239 | 18.378539 |
| Nu Sil PR10 | *H. numata silvana* | Peru | Novaseq | 32.998323 |
| Nu Sil PR11 | *H. numata silvana* | Brazil | Novaseq | 27.367769 |
| Nu Sil PR12 | *H. numata silvana* | Brazil | Novaseq | 22.223587 |
| Nu Sil PR13 | *H. numata silvana* | Brazil | Novaseq | 29.115604 |
| Nu Sil PR14 | *H. numata silvana* | Brazil | Novaseq | 20.796895 |
| Nu Sil PR2 | *H. numata silvana* | Peru | ERP014239 | 21.735388 |
| Nu Sil PR3 | *H. numata silvana* | Peru | ERP014239 | 17.755915 |
| Nu Sil PR4 | *H. numata silvana* | Peru | ERP123741 | 23.674833 |
| Nu Sil PR5 | *H. numata silvana* | Peru | Novaseq | 21.013074 |
| Nu Sil PR6 | *H. numata silvana* | Peru | Novaseq | 25.87696 |
| Nu Sil PR7 | *H. numata silvana* | Peru | Novaseq | 20.375161 |
| Nu Sil PR8 | *H. numata silvana* | Peru | Novaseq | 30.01565 |
| Nu Sil PR9 | *H. numata silvana* | Peru | Novaseq | 28.786442 |
| Nu Sil FG1 | *H. numata silvana* | Guiana | ERP014239 | 25.851975 |
| Pa But PR1 | *H. pardalinus butleri* | Peru | SRP068426 | 23.909991 |
| Pa But PR2 | *H. pardalinus butleri* | Peru | SRP145614 | 29.622154 |
| Pa But PR3 | *H. pardalinus butleri* | Peru | SRP145614 | 28.498145 |
| Pa But PR4 | *H. pardalinus butleri* | Peru | SRP145614 | 27.339839 |
| Pa Par EC1 | *H. pardalinus pardalinus* | Ecuador | SRP145614 | 29.757004 |
| Pa Par EC2 | *H. pardalinus pardalinus* | Ecuador | SRP145614 | 31.460933 |
| Pa Par EC3 | *H. pardalinus pardalinus* | Ecuador | SRP145614 | 21.781465 |
| Pa Ser PR1 | *H. pardalinus sergestus* | Peru | ERP009041 | 26.89481 |
| Pa Ser PR2 | *H. pardalinus sergestus* | Peru | ERP002440 | 30.220326 |
| Pa Ser PR3 | *H. pardalinus sergestus* | Peru | ERP009041 | 24.425723 |
| Pa Ser PR4 | *H. pardalinus sergestus* | Peru | ERP009041 | 28.859658 |
| Pa Ssp PR1 | *H. pardalinus ssp nov* | Peru | ERP002440 | 26.891751 |
| Pa Ssp PR2 | *H. pardalinus ssp nov* | Peru | ERP009041 | 30.747527 |
| Pa Ssp PR3 | *H. pardalinus ssp nov* | Peru | ERP009041 | 26.917481 |
| Pa Ssp PR4 | *H. pardalinus ssp nov* | Peru | ERP009041 | 24.456507 |

**Table S02. Summary statistics computed among the four species using the coding sites for ABC inferences.**

Summary statistics computed with *mscalc*.
A = *H. hecale melicerta*
B = *H. hecale zuleika*
C = *H. ismenius boulleti*
D = *H. ismenius clarescens*

| Summary statistic | Value |
|---|---|
| bialsites avg | 189.17553 |
| bialsites std | 170.28117 |
| sfAB avg | 0.06734 |
| sfAB std | 0.12062 |
| sfAC avg | 0.11888 |
| sfAC std | 0.15989 |
| sfAD avg | 0.17487 |
| sfAD std | 0.1912 |
| sfBC avg | 0.12161 |
| sfBC std | 0.15993 |
| sfBD avg | 0.17191 |
| sfBD std | 0.18963 |
| sfCD avg | 0.14317 |
| sfCD std | 0.18278 |
| sxA avg | 0.00226 |
| sxA std | 0.00243 |
| sxB avg | 0.00267 |
| sxB std | 0.0027 |
| sxC avg | 0.00109 |
| sxC std | 0.00151 |
| sxD avg | 0.00087 |
| sxD std | 0.00131 |
| ssAB avg | 0.00505 |
| ssAB std | 0.0056 |
| ssAC avg | 0.00019 |
| ssAC std | 0.00068 |
| ssAD avg | 0.00014 |
| ssAD std | 0.00052 |
| ssBC avg | 0.00019 |
| ssBC std | 0.0007 |
| ssBD avg | 0.00015 |
| ssBD std | 0.00054 |

| Summary statistic | Value |
|---|---|
| piA avg | 0.00263 |
| piA std | 0.00247 |
| piB avg | 0.00273 |
| piB std | 0.0025 |
| piC avg | 0.00102 |
| piC std | 0.00117 |
| piD avg | 0.001 |
| piD std | 0.00121 |
| divAB avg | 0.07053 |
| divAB std | 0.12099 |
| divAC avg | 0.12284 |
| divAC std | 0.15969 |
| divAD avg | 0.1788 |
| divAD std | 0.19099 |
| divBC avg | 0.12567 |
| divBC std | 0.15975 |
| divBD avg | 0.17594 |
| divBD std | 0.18944 |
| divCD avg | 0.14447 |
| divCD std | 0.18294 |
| netdivAB avg | 0.06785 |
| netdivAB std | 0.12142 |
| netdivAC avg | 0.12102 |
| netdivAC std | 0.15994 |
| netdivAD avg | 0.17699 |
| netdivAD std | 0.19128 |
| netdivBC avg | 0.12379 |
| netdivBC std | 0.16001 |
| netdivBD avg | 0.17407 |
| netdivBD std | 0.18974 |
| netdivCD avg | 0.14346 |
| netdivCD std | 0.18312 |

| | |
|---|---|
| ssCD avg | 0.00113 |
| ssCD std | 0.00184 |
| pearson r pi AB | 0.89336 |
| pearson r pi AC | 0.3421 |
| pearson r pi AD | 0.31638 |
| pearson r pi BC | 0.34752 |
| pearson r pi BD | 0.32511 |
| pearson r pi CD | 0.72048 |
| thetaA avg | 0.00272 |
| thetaA std | 0.00226 |
| thetaB avg | 0.00291 |
| thetaB std | 0.00237 |
| thetaC avg | 0.00113 |
| thetaC std | 0.00118 |
| thetaD avg | 0.00107 |
| thetaD std | 0.00119 |
| pearson r theta AB | 0.86058 |
| pearson r theta AC | 0.37495 |
| pearson r theta AD | 0.35064 |
| pearson r theta BC | 0.37486 |
| pearson r theta BD | 0.34352 |
| pearson r theta CD | 0.64371 |
| DtajA avg | -0.22338 |
| DtajA std | 0.91617 |
| DtajB avg | -0.33066 |
| DtajB std | 0.89231 |
| DtajC avg | -0.4352 |
| DtajC std | 0.89405 |
| DtajD avg | -0.33302 |
| DtajD std | 0.87502 |
| minDivAB avg | 0.06765 |
| minDivAB std | 0.12117 |
| minDivAC avg | 0.12001 |
| minDivAC std | 0.16008 |
| minDivAD avg | 0.17609 |
| minDivAD std | 0.19143 |
| minDivBC avg | 0.1228 |
| minDivBC std | 0.16016 |
| minDivBD avg | 0.17321 |
| minDivBD std | 0.18988 |
| minDivCD avg | 0.1433 |

| | |
|---|---|
| FST AB avg | 0.35972 |
| FST AB std | 0.45621 |
| FST AC avg | 0.82266 |
| FST AC std | 0.19376 |
| FST AD avg | 0.85856 |
| FST AD std | 0.1826 |
| FST BC avg | 0.82319 |
| FST BC std | 0.19345 |
| FST BD avg | 0.8543 |
| FST BD std | 0.18478 |
| FST CD avg | 0.55827 |
| FST CD std | 0.48872 |
| D AB C avg | -0.02178 |
| D AB C std | 0.46399 |
| D negOne AB C | 243 |
| D posOne AB C | 175 |
| fd AB C avg | -0.01422 |
| fd AB C std | 0.57516 |
| fhom AB C avg | -2.6286 |
| fhom AB C std | 101.7061 |
| D AB D avg | 0.00275 |
| D AB D std | 0.41771 |
| D negOne AB D | 208 |
| D posOne AB D | 159 |
| fd AB D avg | 0.0009 |
| fd AB D std | 0.29696 |
| fhom AB D avg | -0.72753 |
| fhom AB D std | 24.0897 |
| D BA C avg | 0.02178 |
| D BA C std | 0.46399 |
| D negOne BA C | 175 |
| D posOne BA C | 243 |
| fd BA C avg | -0.00324 |
| fd BA C std | 0.99236 |
| fhom BA C avg | -0.34454 |
| fhom BA C std | 149.99488 |
| D BA D avg | -0.00275 |
| D BA D std | 0.41771 |
| D negOne BA D | 159 |
| D posOne BA D | 208 |
| fd BA D avg | -0.02667 |

| | |
|---|---|
| minDivCD std | 0.18295 |
| maxDivAB avg | 0.07502 |
| maxDivAB std | 0.12028 |
| maxDivAC avg | 0.12644 |
| maxDivAC std | 0.15903 |
| maxDivAD avg | 0.18219 |
| maxDivAD std | 0.19025 |
| maxDivBC avg | 0.12935 |
| maxDivBC std | 0.15905 |
| maxDivBD avg | 0.17941 |
| maxDivBD std | 0.18867 |
| maxDivCD avg | 0.14642 |
| maxDivCD std | 0.18253 |
| GminAB avg | 0.36187 |
| GminAB std | 0.46539 |
| GminAC avg | 0.81529 |
| GminAC std | 0.22388 |
| GminAD avg | 0.86402 |
| GminAD std | 0.1964 |
| GminBC avg | 0.81832 |
| GminBC std | 0.22058 |
| GminBD avg | 0.86099 |
| GminBD std | 0.1977 |
| GminCD avg | 0.5607 |
| GminCD std | 0.48891 |
| GmaxAB avg | 2.47282 |
| GmaxAB std | 1.73936 |
| GmaxAC avg | 1.27462 |
| GmaxAC std | 0.51575 |
| GmaxAD avg | 1.19891 |
| GmaxAD std | 0.41111 |
| GmaxBC avg | 1.27107 |
| GmaxBC std | 0.49254 |
| GmaxBD avg | 1.20673 |
| GmaxBD std | 0.42272 |
| GmaxCD avg | 2.14661 |
| GmaxCD std | 1.65553 |

| | |
|---|---|
| fd BA D std | 0.88561 |
| fhom BA D avg | -1.62343 |
| fhom BA D std | 73.12984 |
| D CD A avg | -0.16218 |
| D CD A std | 0.60606 |
| D negOne CD A | 498 |
| D posOne CD A | 396 |
| fd CD A avg | -0.1556 |
| fd CD A std | 0.46788 |
| fhom CD A avg | -30222255514 |
| fhom CD A std | 25314923256491 |
| D CD B avg | -0.14364 |
| D CD B std | 0.60518 |
| D negOne CD B | 463 |
| D posOne CD B | 391 |
| fd CD B avg | -0.15215 |
| fd CD B std | 1.23975 |
| fhom CD B avg | -1.01907 |
| fhom CD B std | 291.78617 |
| D DC A avg | 0.16218 |
| D DC A std | 0.60606 |
| D negOne DC A | 396 |
| D posOne DC A | 498 |
| fd DC A avg | 0.13567 |
| fd DC A std | 0.5296 |
| fhom DC A avg | -6.21522 |
| fhom DC A std | 401.23425 |
| D DC B avg | 0.14364 |
| D DC B std | 0.60518 |
| D negOne DC B | 391 |
| D posOne DC B | 463 |
| fd DC B avg | 0.10683 |
| fd DC B std | 0.79867 |
| fhom DC B avg | 271693992961 |
| fhom DC B std | 21419069644265 |

**TableS03. Model choice among SI models.** SI = Strict Isolation. Results are provided for the Random Forest and ABC neural-network algorithm.

Random Forest:

| | Confusion among homogeneous and heterogeneous models | | |
|---|---|---|---|
| | classified as: | | |
| true models | ***SI Nhomo*** | ***SI Nhetero*** | Error Rate |
| ***SI Nhomo*** | 67336 | 1864 | 0.02693642 |
| ***SI Nhetero*** | 7773 | 61427 | 0.11232659 |
| **Out-of-bag prior error rate: 6.96%** | | | |

| | Posterior probabilities among models | | |
|---|---|---|---|
| best model | ***SI Nhomo*** | ***SI Nhetero*** | **Posterior probability** |
| ***SI Nhetero*** | 193 | 307 | 0.62 |

ABC:

| ABC-neural net results | | |
|---|---|---|
| **P(SI Nhomo)** | **P(SI Nhetero)** | sd |
| 0.003 | 0.997 | 2.67E-08 |

**TableS04. Model choice among SC models.**

SC = Secondary Contact. Results are provided for the Random Forest and ABC neural-network algorithm.

**Random Forest results:**

| | **Confusion among homogeneous and heterogeneous models** | | | | |
|---|---|---|---|---|---|
| | classified as: | | | | |
| **true models** | ***SC Mhomo Nhomo*** | ***SC Mhetero Nhomo*** | ***SC Mhomo Nhetero*** | ***SC Mhetero Nhetero*** | class.error |
| ***SCMhomoNhomo*** | 80274 | 4479 | 32848 | 2199 | 0.32993 |
| ***SCMheteroNhomo*** | 9811 | 76731 | 4672 | 28586 | 0.35950 |
| ***SCMhomoNhetero*** | 33404 | 1820 | 78619 | 5957 | 0.34374 |
| ***SCMheteroNhetero*** | 3889 | 28029 | 11040 | 76842 | 0.3585 |

**Out-of-bag prior error rate: 12.6737%**

| | **Posterior probabilities among models** | | | | |
|---|---|---|---|---|---|
| **best model** | **SC1M1N** | **SC2M1N** | **SC1M2N** | **SC2M2N** | **Post.proba** |
| **SC2M2N** | 46 | 153 | 75 | 226 | 0.5626 |

| | **confusion matrix among SC directionalities (ABC-RF)** | | | | | | | |
|---|---|---|---|---|---|---|---|---|
| | **SC$_{A \leftarrow C}$** | **SC$_{B \leftarrow D}$** | **SC$_{C \leftarrow A}$** | **SC$_{D \leftarrow B}$** | **SC$_{A \leftrightarrow c}$** | **SC$_{C \leftrightarrow D}$** | **SC$_{AC \leftrightarrow BD}$** | class.error |
| **SCA ← c** | 209513 | 808 | 6644 | 2189 | 375 | 11444 | 1830 | 0.100 |
| **SCB ← D** | 779 | 209609 | 1987 | 6701 | 12129 | 274 | 1324 | 0.100 |
| **SCC ← A** | 4886 | 1085 | 210954 | 2320 | 368 | 11443 | 1747 | 0.094 |
| **SCD ← B** | 1100 | 2865 | 2102 | 221332 | 3803 | 268 | 1333 | 0.049 |
| **SCA ↔ c** | 400 | 29056 | 769 | 20553 | 180325 | 191 | 1509 | 0.225 |
| **SCC ↔ D** | 28205 | 371 | 26035 | 750 | 313 | 174937 | 2192 | 0.249 |
| **SCAC ↔ BD** | 2316 | 2293 | 2369 | 2194 | 3507 | 3706 | 216418 | 0.070 |

**Out-of-bag prior error rate: 12.6737%**

| | **Posterior probabilities among SC models (ABC-RF)** | | | | | | | |
|---|---|---|---|---|---|---|---|---|
| | **SC$_{A \leftarrow C}$** | **SC$_{B \leftarrow D}$** | **SC$_{C \leftarrow A}$** | **SC$_{D \leftarrow B}$** | **SC$_{A \leftrightarrow c}$** | **SC$_{C \leftrightarrow D}$** | **SC$_{ABCD}$** | |
| selectedModel | vote | vote | vote | vote | vote | vote | vote | **post.proba** |
| **SCC ← A** | 114 | 90 | **280** | 45 | 23 | 100 | 98 | **0.6153** |

**Table S04** *(continuation)*

| | **Posterior probabilities among SC models (abc neural-net)** | | | | | | |
|---|---|---|---|---|---|---|---|
| **round1** | **p($SC_{A \leftarrow C}$)** | **p($SC_{B \leftarrow D}$)** | **p($SC_{C \leftarrow A}$)** | **p($SC_{D \leftarrow B}$)** | **p($SC_{A \leftrightarrow C}$)** | **p($SC_{C \leftrightarrow D}$)** | **p($SC_{ABCD}$)** |
| | 0.1079 | 0.1184 | **0.2742** | **0.3856** | 0.0610 | 0.0432 | 0.0097 |

| | **Posterior proba. among SC models (abc neural-net)** | | | |
|---|---|---|---|---|
| **round2** | **p($SC_{A \leftarrow C}$)** | **p($SC_{B \leftarrow D}$)** | **p($SC_{C \leftarrow A}$)** | **p($SC_{D \leftarrow B}$)** |
| | 0.1243 | 0.1164 | **0.5963** | 0.1631 |

| | **Posterior probabilities among models (abc neural-net)** | | |
|---|---|---|---|
| **round3** | | **p($SC_{B \leftarrow D}$)** | **p($SC_{C \leftarrow A}$)** |
| | | 0.1232 | **0.8768** |

**TableS05. Model choice among the best alternative models.**

Results are provided for the Random Forest algorithm.

**Random Forest results:**

| ABC-RF | | | ABC-RF Confusion matrix | | | |
|---|---|---|---|---|---|---|
| | **p(SI)** | **p(SCc)** | | **SI** | **SCC** | class.error |
| **probability** | 0.1 | 0.9 | **SI** | 977736 | 45864 | 0.0448 |
| **votes** | 215 | 1285 | **SCC** | 15698 | 1007902 | 0.0153 |

**Comparison against alternative models with ongoing gene flow (IM) or Ancient Gene Flow (AM):**

| ABC-RF | | | ABC-RF Confusion matrix | | | |
|---|---|---|---|---|---|---|
| | **p(IM AC)** | **p(IM BD)** | | **IM AC** | **IM BD** | **class.error** |
| **probability** | 0.75 | 0.25 | **IM AC** | 98471 | 529 | 0.0053 |
| **votes** | 505 | 295 | **IM BD** | 846 | 98154 | 0.0085 |

| ABC-RF | | | ABC-RF Confusion matrix | | | |
|---|---|---|---|---|---|---|
| | **p(AM)** | **p(IM AC)** | | **AM** | **IM AC** | **class.error** |
| **probability** | 0.83 | 0.17 | **AM** | 49320 | 680 | 0.013 |
| **votes** | 434 | 366 | **IM AC** | 361 | 49639 | 0.007 |

| ABC-RF | | | ABC-RF Confusion matrix | | | |
|---|---|---|---|---|---|---|
| | **p(SCc)** | **p(AM)** | | **SCC** | **AM** | **class.error** |
| **probability** | 0.87 | 0.13 | **SCC** | 47555 | 2445 | 0.049 |
| **votes** | 613 | 187 | **AM** | 776 | 49224 | 0.015 |

**Table S06. Parameter Estimation for the two best models using ABC neural net algorithm**

| **Posterior distribution of parameters under SI2N (Tolerance: 0.001)** | | | | | | | |
|---|---|---|---|---|---|---|---|
| ABC nnet | **$Ne_{mel}$** | **$Ne_{zul}$** | **$Ne_{bou}$** | **$Ne_{cla}$** | **$Ne_{hec}$** | **$Ne_{ism}$** | **Nanc** |
| 2.5%CI | 895502 | 996674 | 782199 | 392657 | 2200894 | 3774124 | 2760118 |
| Median | 20506642 | 19456473 | 6098801 | 5971332 | 17952494 | 24282776 | 25063570 |
| Mean | 19894677 | 20099544 | 8170522 | 8698639 | 18722580 | 23398548 | 23630369 |
| 97.5%CI | 38950337 | 39053040 | 32128732 | 34757146 | 38881304 | 39401852 | 39382868 |

| **Posterior distribution of parameters under SI2N (Tolerance: 0.001)** | | | | | | | |
|---|---|---|---|---|---|---|---|
| **ABC nnet** | ***Tsplit H.h melicerta H. h. zuleika*** | **Tsplit *H.i. boulleti H. i.* clarescens** | **Tsplit *H. hecale H. ismenius*** | **M12** | **M21** | **M34** | **M43** |
| 2.5%CI | 74673 | 218370 | 1184065 | 0.000002 | 0.000003 | 0.000000 | 0.000000 |
| Median | 1440158 | 1996364 | 3703290 | 0.000041 | 0.000039 | 0.000006 | 0.000008 |
| Mean | 1654576 | 2058700 | 3509361 | 0.000040 | 0.000040 | 0.000019 | 0.000017 |
| 97.5%CI | 4116594 | 4313305 | 4926088 | 0.000077 | 0.000077 | 0.000073 | 0.000072 |

| **Posterior distribution of parameters under SC2N2Mc (tolerance:0.001)** | | | | | | | |
|---|---|---|---|---|---|---|---|
| ABC nnet | **$Ne_{mel}$** | **$Ne_{zul}$** | **$Ne_{bou}$** | **$Ne_{cla}$** | **$Ne_{hec}$** | **$Ne_{ism}$** | **Nanc** |
| 2.5%CI | 1116554 | 2954990 | 30 | 877668 | 157167 | 1437435 | 6302684 |
| Median | 26775666 | 19630180 | 25814226 | 17787244 | 3688436 | 8889244 | 30255792 |
| Mean | 23829847 | 19315926 | 21247082 | 18117535 | 7625900 | 10896872 | 27877287 |
| 97.5%CI | 39470546 | 34808362 | 39999852 | 37889597 | 35338598 | 30095761 | 39890591 |

| **Posterior distribution of parameters under SC2N2Mc (tolerance:0.001)** | | | | | | | | | |
|---|---|---|---|---|---|---|---|---|---|
| ABC nnet | ***Tsplit H.h melicerta H. h. zuleika*** | **Tsplit *H.i. boulleti H. i.* clarescens** | **Tsplit *H. hecale H. ismenius*** | **$T_{SC}$ *H. i. boulleti H. h. melicerta*** | **M12** | **M21** | **M34** | **M43** | **M31 bou ← mel** |
| 2.5%CI | 120608 | 30064 | 2031518 | 1436 | 0.000015 | 0.000002 | 0.000000 | 0.000000 | 0.000001 |
| Median | 861639 | 1438985 | 3664595 | 131045 | 0.000040 | 0.000031 | 0.000012 | 0.000008 | 0.000022 |
| Mean | 942204 | 1626975 | 3593545 | 339174 | 0.000040 | 0.000035 | 0.000026 | 0.000021 | 0.000028 |
| 97.5%CI | 2237601 | 4186128 | 4771047 | 2014549 | 0.000064 | 0.000076 | 0.000080 | 0.000079 | 0.000074 |

**Table S07. Accuracy of parameter estimation for the ABC-RF algorithm, XGBoost method and ABC neural net method.**

A subset of parameters were tested for the ABC neural net method given its high computing time.

| **Model: SI2N** | | | | **Model: SC2M2Nc** | |
|---|---|---|---|---|---|
| **Parameter** | **$R^2$ – ABCRF** | **$R^2$ – Xgboost** | **$R^2$ – abc** | **Parameter** | **$R^2$- *ABCRF*** |
| N popA | 0.959 | 0.731 | 0.11 | M12 | 0.953 |
| N popB | 0.959 | 0.728 | NA | M21 | 0.953 |
| N popC | 0.957 | 0.679 | NA | M31 | 0.947 |
| N popD | 0.957 | 0.675 | NA | M34 | 0.965 |
| Na AB | 0.959 | 0.684 | NA | M43 | 0.951 |
| Na CD | 0.959 | 0.677 | NA | Na | 0.981 |
| Na AB | 0.988 | 0.932 | NA | Na AB | 0.954 |
| Tsplit | 1 | 0.999 | 0.24 | Na CD | 0.957 |
| Tsplit AB | 0.958 | 0.691 | NA | N popA | 0.957 |
| Tsplit CD | 0.956 | 0.974 | NA | N popB | 0.957 |
| M12 | 0.955 | 0.302 | NA | N popC | 0.957 |
| M21 | 0.955 | 0.296 | NA | N popD | 0.955 |
| M34 | 0.957 | 0.259 | NA | T SC AC | 0.953 |
| M43 | 0.955 | 0.256 | NA | T SC BD | 0.946 |
| ***average*** | *0.962* | *0.635* | *0.175* | Tsplit | 0.977 |
| | | | | Tsplit AB | 0.958 |
| | | | | Tsplit CD | 0.95 |
| | | | | ***average*** | *0.957* |

**Table S08. Summary of ABBA-BABA and related summary statistics results, testing for gene flow at the genome scale.**

Err = standard error obtained through genome bootstraping. Fd and FdM error were zero and are not displayed here.

Abreviation:
*H.h. = Heliconius hecale*
*H.i. = Heliconius ismenius*
*H.n. = Heliconius numata*
*H.p. = Heliconius pardalinus*

| P1 | P2 | P3 | nABBA | nBABA | D | D_Z | D_err | f | f_err | fd | fdM |
|---|---|---|---|---|---|---|---|---|---|---|---|
| *H. h. felix* | *H. h. melicerta* | *H. i. boulleti* | 22,027 | 20,928 | 0.026 | 5.154 | 0.005 | 0.002 | 0.000 | 0.007 | 0.002 |
| *H. h. felix* | *H. h. zuleika* | *H. i. clarescens* | 22,153 | 20,968 | 0.027 | 5.256 | 0.005 | 0.002 | 0.000 | 0.011 | 0.002 |
| *H. h. melicerta* | *H. h. zuleika* | *H. i. clarescens* | 15,082 | 14,985 | 0.003 | 1.183 | 0.003 | 0.000 | 0.000 | 0.005 | 0.000 |
| *H. h. zuleika* | *H. h. melicerta* | *H. i. boulleti* | 14,982 | 15,072 | -0.003 | -1.088 | 0.003 | 0.000 | 0.000 | 0.005 | 0.000 |
| *H. i. boulleti* | *H. i. clarescens* | *H. h. zuleika* | 2,181 | 2,178 | 0.001 | 0.090 | 0.007 | 0.000 | 0.000 | 0.003 | 0.000 |
| *H.i. clarescens* | *H. i. boulleti* | *H. h. melicerta* | 2,176 | 2,171 | 0.001 | 0.166 | 0.007 | 0.000 | 0.000 | 0.003 | 0.000 |
| *H. i. telchninia* | *H. i. clarescens* | *H. h. zuleika* | 2,161 | 2,084 | 0.018 | 1.798 | 0.010 | 0.000 | 0.000 | 0.004 | 0.000 |
| *H. n. silvana* | *H. i. boulleti* | *H. h. melicerta* | 35,516 | 36,589 | -0.015 | -2.383 | 0.006 | -0.006 | 0.002 | 0.014 | -0.001 |
| *H. n. silvana* | *H. i. clarescens* | *H. h. zuleika* | 35,587 | 36,583 | -0.014 | -2.193 | 0.006 | -0.005 | 0.002 | 0.014 | -0.001 |
| *H. p. butleri* | *H. h. melicerta* | *H. i. boulleti* | 43,376 | 37,710 | 0.070 | 12.208 | 0.006 | 0.011 | 0.001 | 0.019 | 0.007 |
| *H. p. butleri* | *H. h. zuleika* | *H. i. clarescens* | 43,492 | 37,760 | 0.071 | 12.083 | 0.006 | 0.011 | 0.001 | 0.019 | 0.007 |
| *H. p. sergestus* | *H. h. melicerta* | *H. i. boulleti* | 43,541 | 37,921 | 0.069 | 11.291 | 0.006 | 0.011 | 0.001 | 0.021 | 0.005 |
| *H. p. sergestus* | *H. h. zuleika* | *H. i. clarescens* | 43,626 | 37,920 | 0.070 | 11.277 | 0.006 | 0.011 | 0.001 | 0.021 | 0.005 |
| *H. p. ssp. Nov* | *H. h. melicerta* | *H. i. boulleti* | 43,479 | 37,834 | 0.069 | 12.085 | 0.006 | 0.011 | 0.001 | 0.019 | 0.007 |
| *H. p. ssp. Nov* | *H. h. zuleika* | *H. i. clarescens* | 43,568 | 37,862 | 0.070 | 12.001 | 0.006 | 0.011 | 0.001 | 0.019 | 0.007 |
| *H. n. silvana* | *H. i. clarescens* | *H. h. zuleika* | 35,656 | 36,504 | -0.012 | -1.830 | 0.006 | -0.004 | 0.002 | 0.014 | 0.000 |
| *H. n. silvana* | *H. i. boulleti* | *H.h. melicerta* | 35,591 | 36,518 | -0.013 | -2.029 | 0.006 | -0.005 | 0.002 | 0.014 | 0.000 |

**Table S09. Summary of *Dfoil* statistics and count of the total number of significant tests (column "sum").**

Count obtained by summing over all tested windows and all possible comparisons.

| | DFO | DIL | DFI | DOL | sum |
|---|---|---|---|---|---|
| **None** | 0 | 0 | 0 | 0 | 506567 |
| **P 1 ⇒ P 3** | + | + | + | 0 | 204 |
| **P 3 ⇒ P 1** | + | 0 | + | + | 277 |
| **P 1 ⇒ P 4** | - | - | 0 | + | 134 |
| **P 4 ⇒ P 1** | - | 0 | + | + | 136 |
| **P 2 ⇒ P 3** | + | + | - | 0 | 106 |
| **P 3 ⇒ P 2** | 0 | + | - | - | 220 |
| **P 2 ⇒ P 4** | - | - | 0 | - | 79 |
| **P 4 ⇒ P 2** | 0 | - | - | - | 139 |
| **P 12 ⇒ P 3** | + | + | 0 | 0 | 47721 |
| **P 3 ⇒ P 12** | + | + | 0 | 0 | 0 |
| **P 12 ⇒ P 4** | - | - | 0 | 0 | 22851 |
| **P 4 ⇒ P 12** | - | - | 0 | 0 | 0 |
| **P 1 ⇔ P 2** | 0 | 0 | 0 | 0 | 0 |
| **P 3 ⇔ P 4** | 0 | 0 | 0 | 0 | 0 |

**Table S10. *Twisst* windows with support for gene flow higher than 0.5.**

Showed are windows considering the sum of topologies 12 and 13 (indicating high gene flow between co-mimics) and topology 9 (indicating gene flow in sympatry). Sum is the sum of topology9 + topology12 + topology13.

| Scaffold | Start | Species tree | Sum | topo9 | topo12 | topo13 |
|---|---|---|---|---|---|---|
| Hmel210001o | 4434865 | 0.021 | 0.729 | 0.122 | 0.608 | 0.000 |
| Hmel210001o | 5469724 | 0.292 | 0.625 | 0.000 | 0.000 | 0.625 |
| Hmel210001o | 8507290 | 0.111 | 0.556 | 0.000 | 0.556 | 0.000 |
| Hmel211001o | 8004111 | 0.125 | 0.531 | 0.000 | 0.375 | 0.156 |
| Hmel212001o | 5390271 | 0.000 | 0.510 | 0.000 | 0.510 | 0.000 |
| Hmel212001o | 7528163 | 0.056 | 0.611 | 0.000 | 0.611 | 0.000 |
| Hmel212001o | 11806306 | 0.292 | 0.583 | 0.000 | 0.000 | 0.583 |
| Hmel212001o | 16228060 | 0.306 | 0.667 | 0.019 | 0.611 | 0.037 |
| Hmel215003o | 355468 | 0.313 | 0.521 | 0.326 | 0.087 | 0.109 |
| Hmel216002o | 5071065 | 0.111 | 0.582 | 0.000 | 0.556 | 0.026 |
| Hmel216002o | 7915534 | 0.111 | 0.576 | 0.000 | 0.556 | 0.021 |
| Hmel218003o | 87840 | 0.278 | 0.566 | 0.000 | 0.556 | 0.010 |
| Hmel218003o | 768867 | 0.375 | 0.583 | 0.194 | 0.194 | 0.194 |
| Hmel218003o | 770020 | 0.034 | 0.536 | 0.201 | 0.000 | 0.335 |
| Hmel218003o | 3200048 | 0.000 | 0.547 | 0.000 | 0.000 | 0.547 |
| Hmel218003o | 10668188 | 0.041 | 0.535 | 0.000 | 0.446 | 0.089 |
| Hmel219001o | 9092355 | 0.417 | 0.583 | 0.000 | 0.583 | 0.000 |
| Hmel219001o | 9155009 | 0.010 | 0.535 | 0.535 | 0.000 | 0.000 |
| Hmel219001o | 9429389 | 0.111 | 0.556 | 0.000 | 0.556 | 0.000 |
| Hmel219001o | 10373736 | 0.139 | 0.528 | 0.003 | 0.418 | 0.106 |
| Hmel201001o | 6786967 | 0.306 | 0.611 | 0.000 | 0.611 | 0.000 |
| Hmel201001o | 9430107 | 0.111 | 0.556 | 0.000 | 0.556 | 0.000 |
| Hmel201001o | 16269414 | 0.032 | 0.535 | 0.000 | 0.356 | 0.178 |
| Hmel201001o | 17126855 | 0.000 | 0.656 | 0.109 | 0.438 | 0.109 |
| Hmel220003o | 4401141 | 0.163 | 0.608 | 0.000 | 0.389 | 0.219 |
| Hmel220003o | 6011509 | 0.229 | 0.512 | 0.130 | 0.130 | 0.252 |

**Table S10** (Continuation)

| Scaffold | Start | Species_tree | Sum | topo9 | topo12 | topo13 |
|---|---|---|---|---|---|---|
| Hmel221001o | 2699593 | 0.191 | 0.688 | 0.000 | 0.573 | 0.115 |
| Hmel221001o | 4882846 | 0.000 | 0.542 | 0.000 | 0.500 | 0.042 |
| Hmel221001o | 5890120 | 0.250 | 0.750 | 0.000 | 0.750 | 0.000 |
| Hmel221001o | 6061731 | 0.000 | 0.917 | 0.000 | 0.917 | 0.000 |
| Hmel221001o | 7242243 | 0.130 | 0.530 | 0.106 | 0.419 | 0.004 |
| Hmel221001o | 11520453 | 0.111 | 0.505 | 0.000 | 0.333 | 0.171 |
| Hmel202001o | 1503076 | 0.035 | 0.583 | 0.003 | 0.288 | 0.292 |
| Hmel202001o | 2081909 | 0.049 | 0.535 | 0.000 | 0.535 | 0.000 |
| Hmel203003o | 5069681 | 0.347 | 0.583 | 0.097 | 0.486 | 0.000 |
| Hmel206001o | 11144177 | 0.306 | 0.611 | 0.000 | 0.611 | 0.000 |
| Hmel208001o | 4408609 | 0.208 | 0.634 | 0.000 | 0.625 | 0.009 |
| Hmel208001o | 5534112 | 0.306 | 0.509 | 0.000 | 0.417 | 0.093 |

**Table S11. Ethogram of the male-female interactions during courtship and mating in *Heliconius hecale***

| Behavioural unit* | Alternative names | Some other butterfly species for which this behaviour has been reported | Description |
|---|---|---|---|
| **Male-related** | | | |
| **Main and minor steps** | | | |
| ***Localisation (A)*** | *Inspection (B), nudging (E)* | *H. melpomene (G), H. erato (B, E), Dryas iulia (C), Bicyclus anynana (A)*, several other species in which males fly almost constantly in search of females (patrolling species, see *F*). | The male approaches the alighted female |
| ***Flickering (A)*** | *Hovering (B, C), fluttering of wings (D), fanning (E)* | *H. erato (B, D, E), Dryas iulia (C)* | The male flights sustainedly over the alighted female (mostly over her head) moving his wings very fast. The forewings and hindwings are very distant from each other, the androconia are exposed. The male can come really close to the female during this step, almost touching her sometimes |
| *Alighting* | *Parallel alighting (C)* | | Male alights very closely besides the female, always keeping forewings and hindwings separated |
| *Slow flapping (B)* | *Wing clapping (C)* | | Standing beside the female, the male opens and closes the wings slowly. Generally linked to *androconia exposition* and *orientation* behaviours |
| *Claspers opening* | | | The male opens his claspers during *slow flapping* and *orientation* |
| *Androconia exposition (B)* | *Silvery friction surfaces exposition (D)* | | The male increases the distance between the forewings and hindwings, exposing the silverish overlapping region between them (androconia) |
| ***Attempting (A)*** | *Abdomen bending (B)* | | Alighted, next to the female, the male bends his abdomen towards hers. His head can be very close to the female's one |
| **Atypical steps** | | | |
| *Alighting on wings (B, C)* | | *H. erato phyllis (B), Dryas iulia (C)* | The male lands on the female wings briefly or sustainedly |
| *Close contact while alighting* | | | The male lands on the basal part of the wings and the thorax of the female, touching her legs with his legs |
| *Orientation* | | | The male changes the angle between his body and the female's one, flapping slowly his wings and exposing androconia. At the end of |

| | | | |
|---|---|---|---|
| | | | this step, the male generally recovers a parallel position in relation to the female |
| **Table S11** (continuation) | | | |
| *Facing* | | *H. erato hydara (D)* | The male faces the female, so that both heads are very close to each other. The antennae seemingly make contact |
| *Touching with proboscis (B, C)* | | *H. erato phyllis (B), Dryas iulia (C), H. erato hydara (D)* | The male extends his proboscis and touches different parts of the female with it |
| **Female-related** | | | |
| **Main and minor steps** | | | |
| ***Female rejection*** | *Mate-refusal posture (J), Posterior vibration (B), abdomen raising and/or wing pressure (C)* | *H. melpomene (G), H. erato phyllis (B), Dryas iulia (C), Agraulis vanilla (H),* several species in the Pieridae family (this behaviour has been called the Pieridae rejection posture; for review see *F*) | The female raises the abdomen to place it above the level of the wings and extrudes the abdominal glands. Simultaneously, her wings are entirely opened and pressed against the ground |
| *Female acceptance* | *Wing shutting (C), contraction(B)* | *H. erato phyllis (B), Dryas iulia (C), Pieris napi (I), Eurema lisa (K), Colias philodice (L)* | The female shuts her wings, descends her abdomen under the hindwings line and remains motionless |
| **Male and female-related** | | | |
| **Main and minor steps** | | | |
| *Flight pursuit (B)* | *Flight (E)* | *H. erato phyllis (B),* several species in the subfamily Heliconiinae *(E)* | The male chases the female while both are flying |
| *Genital contact (A)* | | All species | The male claspers come into contact with the tip of the abdomen of the female, which may precede or not mating |
| ***Success*** | *Copulation (A, D)* | All species | The male grabs the tip of the female's abdomen with his claspers, and they stay attached for a long time (between 15 minutes and many hours) |
| *Rotation* | *Coupled rotation (C)* | | After copulation the male turns the angle of its body, pointing with its head on the opposite direction than the female's one |

*Behaviours that were always recorded during a complete courtship sequence (starting with *localisation* and ending with *success* or *female rejection*) were referred to as *main* events and are highlighted in bold script. We called *minor* steps those behaviours that were less conspicuous but linked to the *main* events, and therefore always present in the courtship sequence. Finally, we described *atypical* steps as those rarely repeated across replicates.

The references from which the information is taken are shown in brackets. *A*: Nieberding *et al.*, 2008; *B:* Mega and Araújo, 2010*;C:* Klein and Araújo, 2010*; D*: Crane, 1955; *E:* Crane, 1957; *F:* Scott, 1972; *G*: Schulz *et al.*, 2008; *H*: Rutowski and Schaefer, 1984; *I*: Forsberg and Wiklund, 1989; *J*: Obara, 1964; *K*: Rutowski, 1977; *L*: Silberglied and Taylor, 1978.

**Table S12. Number of *localisation* events directed toward *H. hecale melicerta* (*melicerta*) and *H. ismenius boulleti* (*boulleti*) female wing models by males of both species.**

(n=42 *melicerta* and n=35 *boulleti*)

| Male | Female | | Male | Female | |
|---|---|---|---|---|---|
| | *melicerta* | *boulleti* | | *melicerta* | *boulleti* |
| *melicerta* 1 | 15 | 20 | *boulleti* 1 | 7 | 3 |
| *melicerta* 2 | 33 | 27 | *boulleti* 2 | 40 | 35 |
| *melicerta* 3 | 3 | 5 | *boulleti* 3 | 4 | 5 |
| *melicerta* 4 | 6 | 5 | *boulleti* 4 | 56 | 64 |
| *melicerta* 5 | 19 | 29 | *boulleti* 5 | 17 | 11 |
| *melicerta* 6 | 11 | 8 | *boulleti* 6 | 7 | 10 |
| *melicerta* 7 | 15 | 11 | *boulleti* 7 | 15 | 9 |
| *melicerta* 8 | 15 | 20 | *boulleti* 8 | 13 | 7 |
| *melicerta* 9 | 15 | 28 | *boulleti* 9 | 11 | 18 |
| *melicerta* 10 | 24 | 37 | *boulleti* 10 | 15 | 23 |
| *melicerta* 11 | 3 | 9 | *boulleti* 11 | 10 | 8 |
| *melicerta* 12 | 14 | 23 | *boulleti* 12 | 13 | 21 |
| *melicerta* 13 | 9 | 9 | *boulleti* 13 | 11 | 18 |
| *melicerta* 14 | 7 | 6 | *boulleti* 14 | 7 | 13 |
| *melicerta* 15 | 13 | 17 | *boulleti* 15 | 3 | 4 |
| *melicerta* 16 | 8 | 6 | *boulleti* 16 | 15 | 13 |
| *melicerta* 17 | 9 | 9 | *boulleti* 17 | 9 | 8 |
| *melicerta* 18 | 25 | 17 | *boulleti* 18 | 3 | 8 |
| *melicerta* 19 | 6 | 7 | *boulleti* 19 | 9 | 6 |
| *melicerta* 20 | 23 | 16 | *boulleti* 20 | 44 | 31 |
| *melicerta* 21 | 11 | 2 | *boulleti* 21 | 18 | 15 |
| *melicerta* 22 | 11 | 11 | *boulleti* 22 | 30 | 28 |
| *melicerta* 23 | 6 | 11 | *boulleti* 23 | 29 | 30 |
| *melicerta* 24 | 10 | 9 | *boulleti* 24 | 12 | 10 |
| *melicerta* 25 | 33 | 33 | *boulleti* 25 | 31 | 32 |
| *melicerta* 26 | 15 | 24 | *boulleti* 26 | 34 | 28 |
| *melicerta* 27 | 8 | 13 | *boulleti* 27 | 45 | 42 |
| *melicerta* 28 | 12 | 20 | *boulleti* 28 | 24 | 33 |
| *melicerta* 29 | 5 | 4 | *boulleti* 29 | 23 | 37 |
| *melicerta* 30 | 22 | 28 | *boulleti* 30 | 20 | 27 |
| *melicerta* 31 | 17 | 19 | *boulleti* 31 | 27 | 35 |
| *melicerta* 32 | 27 | 35 | *boulleti* 32 | 22 | 37 |
| *melicerta* 33 | 33 | 27 | *boulleti* 33 | 24 | 31 |
| *melicerta* 34 | 9 | 9 | *boulleti* 34 | 23 | 36 |
| *melicerta* 35 | 26 | 31 | *boulleti* 35 | 4 | 6 |
| *melicerta* 36 | 11 | 12 | | | |
| *melicerta* 37 | 6 | 12 | | | |
| *melicerta* 38 | 18 | 21 | | | |
| *melicerta* 39 | 37 | 25 | | | |
| *melicerta* 40 | 14 | 12 | | | |
| *melicerta* 41 | 37 | 46 | | | |
| *melicerta* 42 | 47 | 46 | | | |

**Table S13. Number of *hovering* events directed toward *H. hecale melicerta* (*melicerta*) and *H. ismenius boulleti* (*boulleti*) female wing models by males of both species.**

(n=42 *melicerta* and n=35 *boulleti*)

| **Male** | **Female** | | **Male** | **Female** | |
|---|---|---|---|---|---|
| | *melicerta* | *boulleti* | | *melicerta* | *boulleti* |
| *melicerta* 1 | 5 | 2 | *boulleti* 1 | 0 | 0 |
| *melicerta* 2 | 1 | 0 | *boulleti* 2 | 1 | 0 |
| *melicerta* 3 | 0 | 1 | *boulleti* 3 | 1 | 1 |
| *melicerta* 4 | 3 | 3 | *boulleti* 4 | 3 | 2 |
| *melicerta* 5 | 6 | 6 | *boulleti* 5 | 0 | 1 |
| *melicerta* 6 | 7 | 2 | *boulleti* 6 | 0 | 0 |
| *melicerta* 7 | 3 | 3 | *boulleti* 7 | 1 | 0 |
| *melicerta* 8 | 6 | 3 | *boulleti* 8 | 0 | 0 |
| *melicerta* 9 | 0 | 3 | *boulleti* 9 | 0 | 2 |
| *melicerta* 10 | 3 | 0 | *boulleti* 10 | 0 | 0 |
| *melicerta* 11 | 1 | 1 | *boulleti* 11 | 0 | 2 |
| *melicerta* 12 | 4 | 1 | *boulleti* 12 | 0 | 2 |
| *melicerta* 13 | 2 | 0 | *boulleti* 13 | 1 | 1 |
| *melicerta* 14 | 4 | 2 | *boulleti* 14 | 1 | 0 |
| *melicerta* 15 | 0 | 1 | *boulleti* 15 | 0 | 0 |
| *melicerta* 16 | 0 | 0 | *boulleti* 16 | 0 | 1 |
| *melicerta* 17 | 0 | 0 | *boulleti* 17 | 0 | 0 |
| *melicerta* 18 | 0 | 0 | *boulleti* 18 | 3 | 3 |
| *melicerta* 19 | 0 | 0 | *boulleti* 19 | 1 | 2 |
| *melicerta* 20 | 5 | 3 | *boulleti* 20 | 0 | 2 |
| *melicerta* 21 | 1 | 0 | *boulleti* 21 | 1 | 0 |
| *melicerta* 22 | 2 | 1 | *boulleti* 22 | 8 | 6 |
| *melicerta* 23 | 0 | 0 | *boulleti* 23 | 0 | 4 |
| *melicerta* 24 | 0 | 1 | *boulleti* 24 | 3 | 0 |
| *melicerta* 25 | 6 | 0 | *boulleti* 25 | 1 | 1 |
| *melicerta* 26 | 0 | 1 | *boulleti* 26 | 2 | 0 |
| *melicerta* 27 | 0 | 0 | *boulleti* 27 | 0 | 1 |
| *melicerta* 28 | 0 | 1 | *boulleti* 28 | 0 | 1 |
| *melicerta* 29 | 0 | 0 | *boulleti* 29 | 1 | 3 |
| *melicerta* 30 | 0 | 1 | *boulleti* 30 | 0 | 1 |
| *melicerta* 31 | 0 | 0 | *boulleti* 31 | 0 | 0 |
| *melicerta* 32 | 0 | 1 | *boulleti* 32 | 0 | 2 |
| *melicerta* 33 | 3 | 0 | *boulleti* 33 | 0 | 2 |
| *melicerta* 34 | 3 | 1 | *boulleti* 34 | 0 | 0 |
| *melicerta* 35 | 4 | 0 | *boulleti* 35 | 0 | 1 |
| *melicerta* 36 | 4 | 2 | | | |
| *melicerta* 37 | 0 | 1 | | | |
| *melicerta* 38 | 1 | 1 | | | |
| *melicerta* 39 | 2 | 0 | | | |
| *melicerta* 40 | 0 | 0 | | | |
| *melicerta* 41 | 7 | 3 | | | |
| *melicerta* 42 | 10 | 1 | | | |

**Table S14. Compositional similarities and differences between the chemical blends extracted from abdominal glands of males (*C*) and females (*AG*) of *H. hecale melicerta* and *H. ismenius boulleti***

| | | | | Relative concentration (mean)[e] | | | | Relative concentration (standard deviation) | | | | | | Pairwise Wilcoxon rank-sum (*p*-value) | | | |
|---|---|---|---|---|---|---|---|---|---|---|---|---|---|---|---|---|---|
| N° of compound[a] | N° of compound on heatmap[b] | RT[c] | LRI[d] | MFG | MMC | BMC | BFG | MFG | MMC | BMC | BFG | Kruskal-Wallis *H* | *p*-value | MFG-MMC | MFG-BFG | MMC-BMC | BMC-BFG |
| 1 | 1 | 4.631 | 991 | 0.000 | **0.337** | 0.000 | 0.000 | 0.000 | 0.154 | 0.000 | 0.000 | 23.425 | **0.000** | **0.008** | NA | **0.008** | NA |
| 2 | 2 | 5.373 | 1037 | 0.000 | **0.594** | 0.000 | 0.000 | 0.000 | 0.251 | 0.000 | 0.000 | 23.425 | **0.000** | **0.008** | NA | **0.008** | NA |
| 3 | 3 | 5.546 | 1047 | 0.045 | **2.932** | 0.150 | 0.003 | 0.038 | 0.397 | 0.077 | 0.009 | 21.386 | **0.000** | **0.030** | 0.250 | **0.013** | **0.012** |
| 4 | 4 | 5.98 | 1072 | 0.051 | 0.015 | 0.011 | 0.036 | 0.075 | 0.016 | 0.017 | 0.066 | 2.704 | 0.440 | 1.000 | 1.000 | 1.000 | 1.000 |
| 5 | 5 | 6.053 | 1076 | 0.060 | 0.014 | 0.013 | 0.070 | 0.087 | 0.017 | 0.021 | 0.147 | 2.607 | 0.456 | 1.000 | 1.000 | 1.000 | 1.000 |
| 6 | 6 | 6.178 | 1084 | 0.100 | 0.037 | 0.483 | 0.110 | 0.158 | 0.049 | 0.449 | 0.202 | 5.000 | 0.172 | 1.000 | 1.000 | 0.438 | 0.546 |
| 7 | 7 | 6.276 | 1090 | 0.045 | 0.013 | 0.011 | 0.052 | 0.072 | 0.016 | 0.017 | 0.112 | 0.855 | 0.836 | 1.000 | 1.000 | 1.000 | 1.000 |
| 8 | 8 | 6.318 | 1093 | 0.035 | 0.015 | 0.011 | 0.043 | 0.059 | 0.016 | 0.017 | 0.094 | 0.522 | 0.914 | 1.000 | 1.000 | 1.000 | 1.000 |
| 9 | 9 | 6.443 | 1100 | 0.146 | 0.084 | 0.058 | 0.152 | 0.056 | 0.096 | 0.090 | 0.123 | 2.984 | 0.394 | 1.000 | 1.000 | 1.000 | 1.000 |
| 11 | 10 | 6.631 | 1111 | 0.130 | 0.053 | 0.044 | 0.112 | 0.168 | 0.061 | 0.069 | 0.250 | 1.375 | 0.711 | 1.000 | 1.000 | 1.000 | 1.000 |
| 12 | 11 | 6.93 | 1130 | 0.000 | **0.152** | 0.000 | 0.000 | 0.000 | 0.105 | 0.000 | 0.000 | 23.425 | **0.000** | **0.008** | NA | **0.008** | NA |
| 13 | 12 | 9.678 | 1290 | 0.000 | **0.301** | 0.000 | 0.000 | 0.000 | 0.160 | 0.000 | 0.000 | 23.425 | **0.000** | **0.008** | NA | **0.008** | NA |
| 14 | 13 | 13.102 | 1504 | 0.000 | **0.077** | 0.000 | 0.000 | 0.000 | 0.072 | 0.000 | 0.000 | 23.425 | **0.000** | **0.008** | NA | **0.008** | NA |
| 15 | 14 | 14.232 | 1581 | 0.000 | **0.092** | 0.000 | 0.000 | 0.000 | 0.039 | 0.000 | 0.000 | 23.425 | **0.000** | **0.008** | NA | **0.008** | NA |
| 16 | 15 | 14.331 | 1587 | 0.000 | **0.315** | 0.000 | 0.000 | 0.000 | 0.117 | 0.000 | 0.000 | 23.425 | **0.000** | **0.008** | NA | **0.008** | NA |
| 17 | 16 | 14.421 | 1595 | 0.000 | **0.107** | 0.000 | 0.000 | 0.000 | 0.055 | 0.000 | 0.000 | 23.425 | **0.000** | **0.008** | NA | **0.008** | NA |
| 18 | 17 | 15.878 | 1696 | 0.000 | 0.048 | 0.000 | 0.000 | 0.000 | 0.044 | 0.000 | 0.000 | 18.701 | **0.000** | **0.029** | NA | **0.029** | NA |
| 19 | 18 | 16.319 | 1729 | 0.000 | **0.078** | 0.000 | 0.000 | 0.000 | 0.060 | 0.000 | 0.000 | 23.425 | **0.000** | **0.008** | NA | **0.008** | NA |
| 20 | 19 | 17.468 | 1816 | 0.000 | **0.087** | 0.014 | 0.000 | 0.000 | 0.114 | 0.012 | 0.000 | 20.693 | **0.000** | **0.014** | NA | 0.124 | **0.028** |
| 21 | 20 | 17.728 | 1836 | 0.223 | **0.675** | 0.004 | 0.023 | 0.144 | 0.313 | 0.007 | 0.029 | 19.733 | **0.000** | 0.156 | **0.028** | **0.026** | 0.397 |
| 23 | 21 | 17.865 | 1847 | 0.000 | **0.068** | 0.000 | 0.000 | 0.000 | 0.048 | 0.000 | 0.000 | 23.425 | **0.000** | **0.008** | NA | **0.008** | NA |
| 24 | 22 | 18.041 | 1861 | 0.075 | **0.322** | 0.000 | 0.000 | 0.057 | 0.194 | 0.000 | 0.000 | 22.343 | **0.000** | 0.152 | **0.007** | **0.014** | NA |
| 25 | 23 | 18.269 | 1878 | 0.100 | **0.434** | 0.000 | 0.111 | 0.081 | 0.264 | 0.000 | 0.275 | 16.402 | **0.001** | 0.156 | 0.315 | **0.017** | 0.269 |

**Table S14** *(continuation)*

| | | | | | | | | | | | | | | | | | |
|---|---|---|---|---|---|---|---|---|---|---|---|---|---|---|---|---|---|
| 26 | 24 | 18.396 | 1889 | 0.000 | **0.232** | 0.000 | 0.000 | 0.000 | 0.172 | 0.000 | 0.000 | 23.425 | **0.000** | **0.008** | NA | **0.008** | NA |
| 27 | 25 | 18.542 | 1901 | 0.000 | **0.166** | 0.039 | 0.000 | 0.000 | 0.122 | 0.026 | 0.000 | 20.553 | **0.000** | **0.014** | NA | 0.130 | **0.028** |
| 28 | 26 | 18.658 | 1909 | 0.000 | **0.056** | 0.000 | 0.000 | 0.000 | 0.051 | 0.000 | 0.000 | 18.701 | **0.000** | **0.029** | NA | **0.029** | NA |
| 29 | 27 | 18.746 | 1917 | 0.000 | **0.101** | 0.000 | 0.000 | 0.000 | 0.092 | 0.000 | 0.000 | 23.425 | **0.000** | **0.008** | NA | **0.008** | NA |
| 30 | 28 | 18.987 | 1936 | 0.168 | 0.434 | 0.117 | 0.032 | 0.308 | 0.442 | 0.184 | 0.051 | 6.387 | 0.094 | 1.000 | 0.976 | 1.000 | 1.000 |
| 31 | 29 | 19.291 | 1961 | 0.939 | 1.083 | 1.246 | 0.997 | 0.551 | 0.777 | 0.514 | 0.448 | 2.030 | 0.566 | 1.000 | 1.000 | 1.000 | 1.000 |
| 32 | 30 | 19.677 | 1993 | 0.000 | **0.379** | 0.042 | 0.000 | 0.000 | 0.293 | 0.048 | 0.000 | 20.263 | **0.000** | **0.014** | NA | **0.041** | 0.092 |
| 33 | 31 | 19.756 | 2000 | 0.000 | **0.445** | 0.056 | 0.000 | 0.000 | 0.208 | 0.059 | 0.000 | 20.639 | **0.000** | **0.014** | NA | **0.025** | 0.092 |
| 34 | 32 | 19.988 | 2021 | 0.000 | 0.799 | 0.589 | 0.000 | 0.000 | 0.322 | 0.388 | 0.000 | 21.281 | **0.000** | **0.014** | NA | 1.000 | **0.007** |
| 35 | 33 | 20.522 | 2069 | 0.000 | 0.000 | **0.391** | 0.000 | 0.000 | 0.000 | 0.190 | 0.000 | 23.425 | **0.000** | NA | NA | **0.008** | **0.004** |
| 36 | 34 | 20.6 | 2078 | 0.000 | 0.731 | 2.049 | 0.000 | 0.000 | 0.509 | 0.128 | 0.000 | 23.249 | **0.000** | **0.014** | NA | **0.011** | **0.007** |
| 37 | 35 | 20.745 | 2088 | 0.000 | **2.154** | 1.075 | 0.000 | 0.000 | 0.479 | 0.277 | 0.000 | 22.998 | **0.000** | **0.014** | NA | **0.022** | **0.007** |
| 38 | 36 | 20.94 | 2107 | 0.000 | **2.162** | 1.355 | 0.000 | 0.000 | 0.201 | 0.172 | 0.000 | 23.249 | **0.000** | **0.014** | NA | **0.011** | **0.007** |
| 39 | 37 | 21.007 | 2113 | 0.000 | **0.497** | 0.000 | 0.000 | 0.000 | 0.205 | 0.000 | 0.000 | 23.425 | **0.000** | **0.008** | NA | **0.008** | NA |
| 40 | 38 | 21.342 | 2144 | 0.913 | 1.480 | 0.936 | 0.808 | 0.794 | 0.479 | 0.871 | 0.939 | 1.673 | 0.643 | 1.000 | 1.000 | 1.000 | 1.000 |
| 41 | 39 | 21.585 | 2167 | 0.434 | 1.297 | 0.510 | 0.436 | 0.429 | 0.568 | 0.532 | 0.579 | 6.915 | 0.075 | 0.182 | 1.000 | 0.390 | 1.000 |
| 42 | 40 | 21.723 | 2178 | 0.000 | 0.135 | **0.971** | 0.000 | 0.000 | 0.151 | 0.163 | 0.000 | 20.128 | **0.000** | 0.370 | NA | **0.024** | **0.007** |
| 44 | 41 | 22.133 | 2217 | 0.000 | **0.846** | 0.301 | 0.000 | 0.000 | 0.397 | 0.262 | 0.000 | 20.553 | **0.000** | **0.014** | NA | 0.130 | **0.028** |
| 45 | 42 | 22.289 | 2230 | 0.000 | 0.740 | 0.583 | 0.000 | 0.000 | 0.451 | 0.458 | 0.000 | 18.539 | **0.000** | **0.014** | NA | 1.000 | **0.028** |
| 46 | 43 | 22.605 | 2259 | 0.000 | 0.049 | 0.355 | 0.000 | 0.000 | 0.069 | 0.253 | 0.000 | 15.911 | **0.001** | 0.142 | NA | 0.219 | **0.028** |
| 47 | 44 | 22.677 | 2266 | 0.000 | 1.551 | 1.086 | 0.000 | 0.000 | 0.446 | 0.189 | 0.000 | 21.646 | **0.000** | **0.014** | NA | 0.660 | **0.007** |
| 48 | 45 | 22.807 | 2279 | 0.000 | 0.964 | **1.820** | 0.000 | 0.000 | 0.300 | 0.209 | 0.000 | 23.249 | **0.000** | **0.014** | NA | **0.011** | **0.007** |
| 49 | 46 | 22.984 | 2293 | 0.000 | **2.052** | 1.082 | 0.000 | 0.000 | 0.323 | 0.241 | 0.000 | 23.249 | **0.000** | **0.014** | NA | **0.011** | **0.007** |
| 50 | 47 | 23.104 | 2306 | 0.039 | **1.286** | 0.441 | 0.018 | 0.067 | 0.229 | 0.105 | 0.029 | 20.661 | **0.000** | **0.026** | 1.000 | **0.013** | **0.018** |
| 51 | 48 | 23.222 | 2316 | 0.000 | **0.638** | 0.267 | 0.000 | 0.000 | 0.205 | 0.143 | 0.000 | 20.818 | **0.000** | **0.014** | NA | 0.076 | **0.028** |
| 52 | 49 | 23.464 | 2339 | 0.000 | **0.512** | 0.000 | 0.000 | 0.000 | 0.169 | 0.000 | 0.000 | 23.425 | **0.000** | **0.008** | NA | **0.008** | NA |
| 53 | 50 | 23.928 | 2382 | 0.000 | **1.495** | 0.000 | 0.000 | 0.000 | 0.572 | 0.000 | 0.000 | 23.425 | **0.000** | **0.008** | NA | **0.008** | NA |
| 54 | 51 | 24.133 | 2401 | 0.000 | 0.377 | 0.256 | 0.000 | 0.000 | 0.277 | 0.149 | 0.000 | 18.356 | **0.000** | **0.048** | NA | 1.000 | **0.007** |

**Table S14** (*continuation*)

| | | | | | | | | | | | | | | | | | |
|---|---|---|---|---|---|---|---|---|---|---|---|---|---|---|---|---|---|
| 55 | 52 | 24.202 | 2408 | 0.000 | **1.034** | 0.000 | 0.000 | 0.000 | 0.461 | 0.000 | 0.000 | 23.425 | **0.000** | **0.008** | NA | **0.008** | NA |
| 56 | 53 | 24.769 | 2464 | 0.000 | **1.571** | 0.605 | 0.000 | 0.000 | 0.455 | 0.345 | 0.000 | 21.097 | **0.000** | **0.014** | NA | **0.043** | **0.028** |
| 57 | 54 | 24.849 | 2473 | 0.000 | 0.566 | 0.371 | 0.000 | 0.000 | 0.473 | 0.225 | 0.000 | 13.774 | **0.003** | 0.142 | NA | 1.000 | **0.028** |
| 58 | 55 | 25.032 | 2491 | 0.000 | **1.547** | 0.504 | 0.000 | 0.000 | 0.334 | 0.305 | 0.000 | 21.702 | **0.000** | **0.014** | NA | **0.011** | **0.028** |
| 59 | 56 | 25.114 | 2500 | 0.048 | **0.873** | 0.315 | 0.062 | 0.076 | 0.283 | 0.169 | 0.062 | 17.493 | **0.001** | **0.026** | 1.000 | **0.013** | 0.224 |
| 60 | 57 | 25.438 | 2534 | 0.000 | **1.309** | 0.011 | 0.000 | 0.000 | 0.225 | 0.017 | 0.000 | 20.278 | **0.000** | **0.014** | NA | **0.022** | 0.699 |
| 61 | 58 | 25.606 | 2551 | 0.000 | **0.490** | 0.000 | 0.000 | 0.000 | 0.397 | 0.000 | 0.000 | 23.425 | **0.000** | **0.008** | NA | **0.008** | NA |
| 62 | 59 | 25.901 | 2583 | 0.000 | **0.771** | 0.000 | 0.000 | 0.000 | 0.409 | 0.000 | 0.000 | 23.425 | **0.000** | **0.008** | NA | **0.008** | NA |
| 63 | 60 | 26.369 | 2631 | 0.000 | **0.418** | 0.034 | 0.000 | 0.000 | 0.119 | 0.068 | 0.000 | 20.278 | **0.000** | **0.014** | NA | **0.022** | 0.699 |
| 64 | 61 | 26.705 | 2666 | 0.000 | **1.123** | 0.211 | 0.000 | 0.000 | 0.605 | 0.178 | 0.000 | 15.911 | **0.001** | **0.048** | NA | 0.219 | 0.092 |
| 65 | 62 | 26.986 | 2695 | 0.283 | **1.247** | 0.641 | 0.232 | 0.165 | 0.330 | 0.188 | 0.171 | 18.879 | **0.000** | **0.013** | 1.000 | **0.013** | **0.028** |
| 66 | 63 | 27.272 | 2726 | 0.000 | **0.541** | 0.027 | 0.000 | 0.000 | 0.175 | 0.043 | 0.000 | 20.278 | **0.000** | **0.014** | NA | **0.022** | 0.699 |
| 67 | 64 | 27.492 | 2749 | 0.173 | 0.333 | 0.103 | 0.184 | 0.197 | 0.189 | 0.111 | 0.241 | 5.419 | 0.144 | 1.000 | 1.000 | 0.120 | 1.000 |
| 68 | 65 | 27.724 | 27745 | 0.000 | **0.359** | 0.000 | 0.000 | 0.000 | 0.275 | 0.000 | 0.000 | 18.701 | **0.000** | **0.029** | NA | **0.029** | NA |
| 69 | 66 | 27.879 | 2790 | 0.000 | **0.592** | 0.164 | 0.000 | 0.000 | 0.296 | 0.147 | 0.000 | 20.068 | **0.000** | **0.014** | NA | 0.325 | **0.028** |
| 70 | 67 | 27.954 | 2798 | 0.191 | 0.315 | 0.183 | 0.074 | 0.125 | 0.250 | 0.159 | 0.079 | 5.518 | 0.138 | 1.000 | 0.321 | 1.000 | 1.000 |
| 71 | 68 | 28.587 | 2864 | 0.335 | 0.590 | 0.416 | 0.308 | 0.386 | 0.254 | 0.369 | 0.265 | 2.984 | 0.394 | 1.000 | 1.000 | 1.000 | 1.000 |
| 72 | 69 | 28.72 | 2879 | 0.349 | 0.577 | 0.277 | 0.180 | 0.256 | 0.225 | 0.217 | 0.177 | 6.974 | 0.073 | 1.000 | 1.000 | 0.390 | 1.000 |
| 73 | 70 | 29.196 | 2928 | 0.000 | **0.429** | 0.180 | 0.000 | 0.000 | 0.224 | 0.190 | 0.000 | 18.914 | **0.000** | **0.014** | NA | 0.225 | 0.092 |
| 75 | 71 | 29.988 | 3011 | 0.000 | 0.177 | 0.222 | 0.000 | 0.000 | 0.118 | 0.164 | 0.000 | 18.338 | **0.000** | **0.014** | NA | 1.000 | **0.028** |
| 76 | 72 | 30.417 | 3056 | 1.336 | 1.326 | 1.161 | 1.091 | 0.296 | 0.212 | 0.331 | 0.526 | 1.060 | 0.787 | 1.000 | 1.000 | 1.000 | 1.000 |
| 77 | 73 | 30.728 | 3088 | 0.000 | 0.000 | 0.000 | **0.273** | 0.000 | 0.000 | 0.000 | 0.195 | 19.022 | **0.000** | NA | **0.020** | NA | **0.020** |
| 78 | 74 | 30.775 | 3094 | 0.000 | 0.837 | 0.891 | 0.000 | 0.000 | 0.258 | 0.201 | 0.000 | 21.045 | **0.000** | **0.014** | NA | 1.000 | **0.007** |
| 79 | 75 | 30.998 | 3118 | 0.000 | 0.446 | 0.517 | 0.000 | 0.000 | 0.198 | 0.148 | 0.000 | 20.995 | **0.000** | **0.014** | NA | 1.000 | **0.007** |

[a]Unique numeric label for each compound. [b]Label of a compound following the order shown in the respective heatmap. [c]Retention time (minutes). [d]Linear retention index. [e]Values in bold indicate a significant association between a given compound and the category highlighted in bold script, following the *indval* index with $p<0.001$. MFG: glands from *H. h. melicerta* females. MMC: claspers from *H. h. melicerta* males. BMC: claspers from *H. i. boulleti* males. BFG: glands from *H. i. boulleti* females.

Code of cell colours:

Unique to males
Unique to *H. hecale*
Unique to *H. hecale melicerta* males
Unique to *H. ismenius boulleti* males
Unique to *H. ismenius boulleti* females

**Table S15. Compositional similarities and differences between the chemical blends extracted from wings (*OW*) of males and females of *H. hecale melicerta* and *H. ismenius boulleti***

| | | | | Relative concentration (mean)[e] | | | | Relative concentration (standard deviation) | | | | | | Pairwise Wilcoxon rank-sum (*p*-value) | | | |
|---|---|---|---|---|---|---|---|---|---|---|---|---|---|---|---|---|---|
| **N° of compound [a]** | **N° of compound on heatmap [b]** | **RT[c]** | **LRI[d]** | **MFOW** | **MMOW** | **BMOW** | **BFOW** | **MFOW** | **MMOW** | **BMOW** | **BFOW** | **Kruskal-Wallis *H*** | ***p*-value** | **MFOW - MMOW** | **MFOW-BFOW** | **MMOW - BMOW** | **BMOW-BFOW** |
| 3 | 1 | 5.546 | 1047 | 0.029 | 0.057 | 0.000 | 0.000 | 0.057 | 0.054 | 0.000 | 0.000 | 6.764 | 0.080 | 1.000 | 1.000 | 0.251 | NA |
| 4 | 2 | 5.98 | 1072 | 0.114 | 0.081 | 0.037 | 0.040 | 0.179 | 0.132 | 0.003 | 0.005 | 0.400 | 0.940 | 1.000 | 1.000 | 1.000 | 1.000 |
| 5 | 3 | 6.053 | 1076 | 0.129 | 0.082 | 0.047 | 0.042 | 0.202 | 0.119 | 0.005 | 0.006 | 0.825 | 0.843 | 1.000 | 1.000 | 1.000 | 1.000 |
| 6 | 4 | 6.178 | 1084 | 0.201 | 0.129 | 0.073 | 0.078 | 0.263 | 0.166 | 0.008 | 0.001 | 0.746 | 0.862 | 1.000 | 1.000 | 1.000 | 1.000 |
| 7 | 5 | 6.276 | 1090 | 0.103 | 0.072 | 0.032 | 0.033 | 0.160 | 0.119 | 0.004 | 0.003 | 0.423 | 0.935 | 1.000 | 1.000 | 1.000 | 1.000 |
| 8 | 6 | 6.318 | 1093 | 0.085 | 0.061 | 0.027 | 0.036 | 0.134 | 0.101 | 0.003 | 0.012 | 1.070 | 0.784 | 1.000 | 1.000 | 1.000 | 1.000 |
| 9 | 7 | 6.443 | 1100 | 0.075 | 0.142 | 0.179 | 0.161 | 0.087 | 0.070 | 0.020 | 0.008 | 6.081 | 0.108 | 1.000 | 0.650 | 1.000 | 1.000 |
| 10 | 8 | 6.508 | 1104 | 0.013 | 0.043 | 0.044 | 0.028 | 0.025 | 0.050 | 0.030 | 0.024 | 3.269 | 0.352 | 1.000 | 1.000 | 1.000 | 1.000 |
| 11 | 9 | 6.631 | 1111 | 0.222 | 0.175 | 0.116 | 0.127 | 0.310 | 0.213 | 0.011 | 0.014 | 1.429 | 0.699 | 1.000 | 1.000 | 1.000 | 1.000 |
| 22 | 10 | 17.813 | 1843 | 0.000 | **0.505** | 0.007 | 0.000 | 0.000 | 0.163 | 0.005 | 0.000 | 14.976 | **0.002** | **0.045** | NA | **0.030** | 0.593 |
| 31 | 11 | 19.291 | 1961 | 0.031 | 0.120 | 0.166 | 0.094 | 0.063 | 0.142 | 0.143 | 0.066 | 3.441 | 0.328 | 1.000 | 1.000 | 1.000 | 1.000 |
| 33 | 12 | 19.756 | 2000 | 0.000 | **0.027** | 0.000 | 0.000 | 0.000 | 0.013 | 0.000 | 0.000 | 15.729 | **0.001** | **0.027** | NA | **0.027** | NA |
| 34 | 13 | 19.988 | 2021 | 0.014 | 0.053 | 0.007 | 0.012 | 0.013 | 0.035 | 0.005 | 0.021 | 8.320 | **0.040** | 0.349 | 1.000 | 0.073 | 1.000 |
| 36 | 14 | 20.6 | 2078 | 0.017 | 0.032 | 0.005 | 0.000 | 0.035 | 0.024 | 0.005 | 0.000 | 6.902 | 0.075 | 1.000 | 1.000 | 0.268 | 1.000 |
| 38 | 15 | 20.94 | 2107 | 0.000 | **1.295** | 0.000 | 0.000 | 0.000 | 0.267 | 0.000 | 0.000 | 15.729 | **0.001** | **0.027** | NA | **0.027** | NA |
| 40 | 16 | 21.342 | 2144 | 0.039 | 0.068 | 0.045 | 0.245 | 0.077 | 0.122 | 0.058 | 0.236 | 2.552 | 0.466 | 1.000 | 1.000 | 1.000 | 1.000 |

| | | | | | | | | | | | | | | | | | |
|---|---|---|---|---|---|---|---|---|---|---|---|---|---|---|---|---|---|
| 43 | 17 | 22.016 | 2205 | 0.000 | **0.192** | 0.000 | 0.000 | 0.000 | 0.113 | 0.000 | 0.000 | 12.673 | **0.005** | 0.068 | NA | 0.068 | NA |
| 45 | 18 | 22.289 | 2230 | 0.000 | **0.045** | 0.000 | 0.000 | 0.000 | 0.039 | 0.000 | 0.000 | 15.750 | **0.001** | **0.027** | NA | **0.027** | NA |
| 47 | 19 | 22.677 | 2266 | 0.000 | 0.112 | 0.000 | 0.000 | 0.000 | 0.104 | 0.000 | 0.000 | 12.673 | **0.005** | 0.068 | NA | 0.068 | NA |
| 48 | 20 | 22.807 | 2279 | 0.029 | 0.067 | 0.000 | 0.000 | 0.058 | 0.062 | 0.000 | 0.000 | 11.836 | **0.008** | 0.525 | 1.000 | **0.045** | NA |
| 50 | 21 | 23.104 | 2306 | 0.012 | **0.324** | 0.000 | 0.006 | 0.014 | 0.123 | 0.000 | 0.006 | 13.844 | **0.003** | 0.063 | 1.000 | 0.054 | 0.736 |
| 52 | 22 | 23.464 | 2339 | 0.000 | 0.040 | 0.000 | 0.000 | 0.000 | 0.038 | 0.000 | 0.000 | 12.673 | **0.005** | 0.068 | NA | 0.068 | NA |
| 53 | 23 | 23.928 | 2382 | 0.000 | **0.178** | 0.000 | 0.000 | 0.000 | 0.134 | 0.000 | 0.000 | 15.729 | **0.001** | **0.027** | NA | **0.027** | NA |
| 54 | 24 | 24.133 | 2401 | 0.000 | **0.216** | 0.000 | 0.000 | 0.000 | 0.177 | 0.000 | 0.000 | 15.729 | **0.001** | **0.027** | NA | **0.027** | NA |
| 59 | 25 | 25.114 | 2500 | 0.054 | 0.137 | 0.000 | 0.017 | 0.037 | 0.209 | 0.000 | 0.017 | 9.816 | **0.020** | 1.000 | 1.000 | 0.054 | 0.736 |
| 60 | 26 | 25.438 | 2534 | 0.068 | 0.341 | 0.000 | 0.000 | 0.127 | 0.323 | 0.000 | 0.000 | 12.576 | **0.006** | 0.360 | 1.000 | **0.045** | NA |
| 65 | 27 | 26.986 | 2695 | 0.184 | 0.245 | 0.046 | 0.257 | 0.116 | 0.256 | 0.037 | 0.098 | 7.302 | 0.063 | 1.000 | 1.000 | 0.255 | 0.343 |
| 72 | 28 | 28.72 | 2879 | 0.296 | 0.244 | 0.041 | 0.196 | 0.181 | 0.256 | 0.020 | 0.022 | 9.797 | **0.020** | 1.000 | 1.000 | **0.036** | 0.343 |
| 74 | 29 | 29.894 | 3001 | 0.143 | 0.207 | 0.127 | 0.007 | 0.184 | 0.144 | 0.020 | 0.006 | 6.828 | 0.078 | 1.000 | 1.000 | 1.000 | 0.343 |
| 76 | 30 | 30.417 | 3056 | 0.396 | 0.264 | 0.170 | 0.519 | 0.242 | 0.266 | 0.075 | 0.129 | 4.604 | 0.203 | 1.000 | 1.000 | 1.000 | 0.343 |
| 77 | 31 | 30.728 | 3088 | 0.063 | 0.000 | 0.000 | 0.008 | 0.060 | 0.000 | 0.000 | 0.013 | 9.910 | **0.019** | 0.081 | 0.995 | NA | 1.000 |
| 78 | 32 | 30.775 | 3094 | 0.000 | 0.125 | 0.048 | 0.000 | 0.000 | 0.188 | 0.021 | 0.000 | 13.138 | **0.004** | **0.045** | NA | 1.000 | 0.218 |
| 79 | 33 | 30.998 | 3118 | 0.044 | 0.000 | 0.000 | 0.000 | 0.055 | 0.000 | 0.000 | 0.000 | 11.784 | **0.008** | **0.049** | 0.356 | NA | NA |

[a]Unique numeric label for each compound. [b]Label of a compound following the order shown in the respective heatmap. [c]Retention time (minutes). [d]Linear retention index. [e]Values in bold indicate a significant association between a given compound and the category highlighted in bold script, following the *indval* index with $p<0.001$. MFOW: wings from *H. h. melicerta* females. MMOW: wings from *H. h. melicerta* males. BMOW: wings from *H. i. boulleti* males. BFOW: wings from *H. i. boulleti* females.

Code of cell colours:

- Unique to males
- Unique to females
- Unique to *H. hecale*
- Unique to *H. hecale melicerta* males
- Unique to *H. hecale melicerta* females

**Table S16. Compositional similarities and differences between the chemical blends extracted from the cuticle near the insertion of the wings (*B*) of males and females of *H. hecale melicerta* and *H ismenius boulleti***

| | | | | Relative concentration (mean)[e] | | | | Relative concentration (standard deviation) | | | | | | Pairwise Wilcoxon rank-sum (*p*-value) | | | |
|---|---|---|---|---|---|---|---|---|---|---|---|---|---|---|---|---|---|
| N° of compound[a] | N° of compound on heatmap[b] | RT[c] | LRI[d] | MFB | MMB | BMB | BFB | MFB | MMB | BMB | BFB | Kruskal-Wallis *H* | *p*-value | MFB-MMB | MFB-BFB | MMB-BMB | BMB-BFB |
| 3 | 1 | 5.546 | 1047 | 0.009 | 0.100 | 0.000 | 0.011 | 0.019 | 0.054 | 0.000 | 0.015 | 12.088 | **0.007** | 0.178 | 1.000 | 0.065 | 1.000 |
| 4 | 2 | 5.98 | 1072 | 0.022 | 0.030 | 0.016 | 0.032 | 0.035 | 0.005 | 0.022 | 0.020 | 2.415 | 0.491 | 1.000 | 1.000 | 1.000 | 1.000 |
| 5 | 3 | 6.053 | 1076 | 0.022 | 0.037 | 0.019 | 0.033 | 0.033 | 0.006 | 0.027 | 0.019 | 1.803 | 0.614 | 1.000 | 1.000 | 1.000 | 1.000 |
| 6 | 4 | 6.178 | 1084 | 0.054 | 0.060 | 0.025 | 0.083 | 0.120 | 0.009 | 0.035 | 0.063 | 4.560 | 0.207 | 0.956 | 1.000 | 1.000 | 0.636 |
| 7 | 5 | 6.276 | 1090 | 0.015 | 0.029 | 0.010 | 0.027 | 0.021 | 0.004 | 0.014 | 0.016 | 3.777 | 0.287 | 1.000 | 1.000 | 0.266 | 0.959 |
| 8 | 6 | 6.318 | 1093 | 0.013 | 0.025 | 0.005 | 0.024 | 0.018 | 0.005 | 0.011 | 0.014 | 5.754 | 0.124 | 1.000 | 1.000 | 0.178 | 0.448 |
| 9 | 7 | 6.443 | 1100 | 0.144 | 0.155 | 0.141 | 0.165 | 0.090 | 0.009 | 0.043 | 0.033 | 2.544 | 0.467 | 1.000 | 1.000 | 1.000 | 1.000 |
| 10 | 8 | 6.508 | 1104 | 0.010 | 0.081 | 0.009 | 0.042 | 0.022 | 0.006 | 0.019 | 0.050 | 8.618 | **0.035** | 0.090 | 1.000 | 0.090 | 1.000 |
| 11 | 9 | 6.631 | 1111 | 0.025 | 0.070 | 0.026 | 0.063 | 0.034 | 0.013 | 0.036 | 0.035 | 6.873 | 0.076 | 0.632 | 0.402 | 1.000 | 0.402 |
| 30 | 10 | 18.987 | 1936 | 0.188 | 0.191 | 0.037 | 0.053 | 0.166 | 0.076 | 0.042 | 0.068 | 6.808 | 0.078 | 1.000 | 1.000 | 0.117 | 1.000 |
| 31 | 11 | 19.291 | 1961 | 1.413 | 0.999 | 0.946 | 0.961 | 0.280 | 0.119 | 0.397 | 0.443 | 5.097 | 0.165 | 0.381 | 0.905 | 1.000 | 1.000 |
| 38 | 12 | 20.94 | 2107 | 0.024 | 0.206 | 0.018 | 0.005 | 0.040 | 0.186 | 0.028 | 0.011 | 10.050 | **0.018** | 0.272 | 1.000 | 0.108 | 1.000 |
| 40 | 13 | 21.342 | 2144 | 1.968 | 1.702 | 1.681 | 1.498 | 0.352 | 0.177 | 0.631 | 0.599 | 2.710 | 0.438 | 1.000 | 1.000 | 1.000 | 1.000 |
| 41 | 14 | 21.585 | 2167 | 0.636 | 0.496 | 0.859 | 0.728 | 0.661 | 0.168 | 0.496 | 0.417 | 2.364 | 0.500 | 1.000 | 1.000 | 1.000 | 1.000 |
| 49 | 15 | 22.984 | 2293 | 0.007 | 0.027 | 0.000 | 0.000 | 0.016 | 0.022 | 0.000 | 0.000 | 9.023 | **0.029** | 0.888 | 1.000 | 0.208 | NaN |
| 50 | 16 | 23.104 | 2306 | 0.027 | 0.091 | 0.000 | 0.020 | 0.037 | 0.067 | 0.000 | 0.028 | 8.195 | **0.042** | 1.000 | 1.000 | 0.065 | 1.000 |
| 59 | 17 | 25.114 | 2500 | 0.151 | 0.135 | 0.008 | 0.043 | 0.081 | 0.112 | 0.018 | 0.033 | 12.589 | **0.006** | 1.000 | 0.190 | 0.090 | 0.713 |
| 60 | 18 | 25.438 | 2534 | 0.128 | 0.232 | 0.000 | 0.000 | 0.125 | 0.132 | 0.000 | 0.000 | 12.254 | **0.007** | 1.000 | 0.360 | 0.054 | NaN |
| 63 | 19 | 26.369 | 2631 | 0.000 | 0.030 | 0.000 | 0.000 | 0.000 | 0.016 | 0.000 | 0.000 | 17.690 | **0.001** | **0.032** | NaN | **0.032** | NaN |
| 64 | 20 | 26.705 | 2666 | 0.000 | **0.030** | 0.000 | 0.000 | 0.000 | 0.016 | 0.000 | 0.000 | 17.690 | **0.001** | **0.032** | NaN | **0.032** | NaN |
| 65 | 21 | 26.986 | 2695 | 0.309 | 0.329 | 0.183 | 0.184 | 0.080 | 0.100 | 0.147 | 0.089 | 6.052 | 0.109 | 1.000 | 0.333 | 0.667 | 1.000 |
| 66 | 22 | 27.272 | 2726 | 0.015 | 0.056 | 0.014 | 0.000 | 0.021 | 0.031 | 0.031 | 0.000 | 9.584 | **0.022** | 0.632 | 1.000 | 0.330 | 1.000 |
| 67 | 23 | 27.492 | 2749 | 0.011 | 0.049 | 0.000 | 0.000 | 0.024 | 0.037 | 0.000 | 0.000 | 9.338 | **0.025** | 0.700 | 1.000 | 0.208 | NaN |
| 69 | 24 | 27.879 | 2790 | 0.024 | 0.027 | 0.011 | 0.005 | 0.044 | 0.021 | 0.024 | 0.011 | 3.153 | 0.369 | 1.000 | 1.000 | 1.000 | 1.000 |

**Table S16** (continuation)

| | | | | | | | | | | | | | | | | | |
|---|---|---|---|---|---|---|---|---|---|---|---|---|---|---|---|---|---|
| 70 | 25 | 27.954 | 2798 | 0.096 | 0.018 | 0.017 | 0.018 | 0.096 | 0.021 | 0.024 | 0.018 | 2.888 | 0.409 | 1.000 | 1.000 | 1.000 | 1.000 |
| 71 | 26 | 28.587 | 2864 | 0.045 | 0.043 | 0.000 | 0.012 | 0.086 | 0.013 | 0.000 | 0.017 | 8.624 | **0.035** | 1.000 | 1.000 | 0.062 | 1.000 |
| 72 | 27 | 28.72 | 2879 | 0.147 | 0.158 | 0.201 | 0.093 | 0.115 | 0.090 | 0.318 | 0.062 | 1.770 | 0.621 | 1.000 | 1.000 | 1.000 | 1.000 |
| 73 | 28 | 29.196 | 2928 | 0.058 | 0.057 | 0.042 | 0.011 | 0.062 | 0.071 | 0.072 | 0.025 | 2.611 | 0.456 | 1.000 | 0.948 | 1.000 | 1.000 |
| 74 | 29 | 29.894 | 3001 | 0.000 | 0.015 | 0.000 | 0.008 | 0.000 | 0.017 | 0.000 | 0.017 | 4.719 | 0.194 | 0.656 | 1.000 | 0.656 | 1.000 |
| 75 | 30 | 29.988 | 3011 | 0.006 | 0.010 | 0.007 | 0.007 | 0.013 | 0.019 | 0.015 | 0.015 | 0.202 | 0.977 | 1.000 | 1.000 | 1.000 | 1.000 |
| 76 | 31 | 30.417 | 3056 | 1.174 | 0.757 | 0.894 | 0.890 | 0.353 | 0.035 | 0.332 | 0.260 | 3.919 | 0.270 | 0.667 | 1.000 | 1.000 | 1.000 |
| 77 | 32 | 30.728 | 3088 | 0.000 | 0.000 | 0.000 | 0.175 | 0.000 | 0.000 | 0.000 | 0.249 | 13.208 | **0.004** | NaN | 0.076 | NaN | 0.076 |
| 78 | 33 | 30.775 | 3094 | 0.033 | 0.104 | 0.212 | 0.182 | 0.031 | 0.078 | 0.199 | 0.227 | 4.494 | 0.213 | 1.000 | 1.000 | 1.000 | 1.000 |
| 79 | 34 | 30.998 | 3118 | 0.026 | 0.000 | 0.103 | 0.024 | 0.041 | 0.000 | 0.116 | 0.034 | 6.449 | 0.092 | 1.000 | 1.000 | 0.265 | 0.636 |

[a]Unique numeric label for each compound. [b]Label of a compound following the order shown in the respective heatmap. [c]Retention time (minutes). [d]Linear retention index. [e]Values in bold indicate a significant association between a given compound and the category highlighted in bold script, following the *indval* index with $p<0.001$. MFB: cuticle from *H. h. melicerta* females. MMB: cuticle from *H. h. melicerta* males. BMB: cuticle from *H. i. boulleti* males. BFB: cuticle from *H. i. boulleti* females.

Code of cell colours:

Unique to *H. hecale*

Unique to *H. hecale melicerta* males

Unique to *H. ismenius boulleti* females

**Table S17. Tentative identification of compounds isolated from abdominal glands and wings of males and females of *H. hecale melicerta* and *H. ismenius boulleti***

| Label | | LRI[a] | Class | MW | Tentative identification[b] | Outstanding observation |
|---|---|---|---|---|---|---|
| X1 | 1 | 991 | Monoterpene | 136 | Myrcene[1,2] | |
| X2 | 2 | 1037 | Monoterpene | 136 | (*Z*)-β-ocimene[2] | |
| X3 | 3 | 1047 | Monoterpene | 136 | (*E*)-β-ocimene[1,2] | Dominant in *H. h. melicerta*'s claspers |
| X4 | 4 | 1072 | Hydrocarbon | | | |
| X5 | 5 | 1076 | Hydrocarbon | | | |
| X6 | 6 | 1084 | Phenol | 124 | O-methoxyphenol[2] | |
| X7 | 7 | 1090 | Hydrocarbon | | | |
| X8 | 8 | 1093 | Hydrocarbon | | | |
| X9 | 9 | 1100 | Hydrocarbon-alkane | 156 | Undecane | |
| X10 | 10 | 1104 | Unknown | | | |
| X11 | 11 | 1111 | Hydrocarbon | | | |
| X12 | 12 | 1130 | Monoterpene | 136 | Alloocimene[1,2] | |
| X13 | 13 | 1290 | Tetrahydropyran | 194 | Dihydroedulan[1] | |
| X14 | 14 | 1504 | Sesquiterpene | 204 | α-farnesene[1,2] | |
| X15 | 15 | 1581 | Unknown $M^+$224 | | | |
| X16 | 16 | 1587 | Unknown $M^+$224 | | | |
| X17 | 17 | 1595 | Unknown $M^+$222 | | | |
| X18 | 18 | 1696 | Unknown | | | |
| X19 | 19 | 1729 | Unknown | | | |
| X20 | 20 | 1816 | Unknown | | | |
| X21 | 21 | 1836 | Terpene | 278 | | |
| X22 | 22 | 1843 | Terpene | 268 | Hexahydrofarnesyl acetone (synonym: 6,10,14-trimethyl-2-pentadecanone) | Unique for *H. h. melicerta*'s male wings |
| X23 | 23 | 1847 | Unknown | | | |
| X24 | 24 | 1861 | Terpene | 278 | | Unique for *H. m. melicerta*'s glands and claspers |
| X25 | 25 | 1878 | Terpene | 278 | | |
| X26 | 26 | 1889 | Unknown | | | |
| X27 | 27 | 1901 | Hydrocarbon-alkane | 268 | Nonadecane[2] | |
| X28 | 28 | 1909 | Unknown | | | |
| X29 | 29 | 1917 | Unknown | | | |
| X30 | 30 | 1936 | Fatty acid | 254 | Hexadecenoic acid[2] | |
| X31 | 31 | 1961 | Fatty acid | 256 | Hexadecanoic acid[2] | |
| X32 | 32 | 1993 | Unknown $M^+$248 | | | |
| X33 | 33 | 2000 | Hydrocarbon-alkane | 282 | Eicosane[2] | |
| X34 | 34 | 2021 | Aldehyde | 250 | Octadecanal | |
| X35 | 35 | 2069 | Hydrocarbon $M^+$ 292 | 292 | | Unique in *H. i. boulleti*'s claspers |

| | | | | | | |
|---|---|---|---|---|---|---|
| X36 | 36 | 2078 | Hydrocarbon-alkene | 294 | Heneicosene[2] | Higher in *H. i. boulleti*'s than in *H. h. melicerta*'s claspers |
| X37 | 37 | 2088 | Unknown M | 252 | | |
| X38 | 38 | 2107 | Hydrocarbon-alkane | 296 | Heneicosane[2] | Dominant in *H. h. melicerta*'s male wings |
| X39 | 39 | 2113 | Unknown | | | |
| X40 | 40 | 2144 | Fatty acid | 280 + 282 | Octadecadienoic acid + Octadecenoic acid | |
| X41 | 41 | 2167 | Fatty acid | 284 | Octadecanoic acid | |
| X42 | 42 | 2178 | Hydrocarbon-alkene | 308 | Docosene | Higher in *H. i. boulleti*'s than in *H. h. melicerta*'s claspers |
| X43 | 43 | 2205 | Hydrocarbon-alkane | 310 | Docosane | |
| X44 | 44 | 2217 | Ester | 312 | Octadecyl acetate[1] | |
| X45 | 45 | 2230 | Aldehyde | 296 | Eicosanal | |
| X46 | 46 | 2259 | Unknown $M^+$ 294 | | | |
| X47 | 47 | 2266 | Unknown $M^+$ 278 | | | |
| X48 | 48 | 2279 | Hydrocarbon-alkene | 322 | Tricosene[2] | Higher in *H. i. boulleti*'s than in *H. h. melicerta*'s claspers |
| X49 | 49 | 2293 | Unknown $M^+$ 280 | | | |
| X50 | 50 | 2306 | Hydrocarbon-alkane | 324 | Tricosane[2] | |
| X51 | 51 | 2316 | Unknown | | | |
| X52 | 52 | 2339 | Hydrocarbon-alkane | 338 | 11-methyl-tricosane[2] | |
| X53 | 53 | 2382 | Ester | 338 | Eicosenyl acetate[1] | |
| X54 | 54 | 2401 | Unknown $M^+$322 | | | |
| | | | Hydrocarbon-alkane | 338 | Tetracosane | |
| X55 | 55 | 2408 | Ester | 340 | Eicosanyl acetate[1] | |
| | | | Hydrocarbon-alkane | 352 | 11-methyl-tetracosane | |
| X56 | 56 | 2464 | Unknown $M^+$324 | | | |
| X57 | 57 | 2473 | Hydrocarbon-alkene | 350 | Pentacosene | |
| X58 | 58 | 2491 | Unknown | | | |
| X59 | 59 | 2500 | Hydrocarbon-alkane | 352 | Pentacosane[2] | |
| X60 | 60 | 2534 | Hydrocarbon-alkane | 366 | 11-methyl-pentacosane[2] | |
| X61 | 61 | 2551 | Unknown $M^+$ 364 | | | |
| X62 | 62 | 2583 | Ester | 366 | Docosenyl acetate[1] | |
| X63 | 63 | 2631 | Hydrocarbon-alkane | 380 | Methyl-hexacosane | |
| X64 | 64 | 2666 | Unknown | | | |
| X65 | 65 | 2695 | Unknown $M^+$ | 380 | Heptacosane[2] | |

| | | | | | | |
|---|---|---|---|---|---|---|
| | | | 336[c] + alkane | | | |
| X66 | 66 | 2726 | Hydrocarbon-alkane | | Branched heptacosane | |
| X67 | 67 | 2749 | Hydrocarbon-alkane | | Branched heptacosane | |
| X68 | 68 | 2774 | Ester $M^+$334 | | | |
| X69 | 69 | 2790 | Unknown[c] + alkane | 394 | Octacosane | |
| X70 | 70 | 2798 | Triterpene | 410 | Squalene[2] | |
| X71 | 71 | 2864 | Steroid | | | |
| X72 | 72 | 2879 | Hydrocarbon-alkane | 408 | Nonacosane[2] | |
| X73 | 73 | 2928 | Hydrocarbon-alkane | 436 | 11,15-dimethyl-nonacosane[2] | |
| X74 | 74 | 3001 | Unknown $M^+$ 390 | | | |
| X75 | 75 | 3011 | Hydrocarbon-alkane | | | |
| X76 | 76 | 3056 | Steroid | 386 | Cholesterol[2] | |
| X77 | 77 | 3088 | Steroid | 398 | | Unique for female wings and *H. i. boulleti*'s glands and cuticle |
| X78 | 78 | 3094 | Hydrocarbon-alkane | 364 | 11,17-dimethyl hentriacontane | |
| X79 | 79 | 3118 | Tetrahydrofuran | 450 | 2-eicosanyl-5-heptyl-tetrahydrofuran[3] | Exclusive to *H. h. melicerta's* female wings |

[a] Linear retention indices calculated with a Rtx-5 MS fused silica capillary column

[b]Straight-chain hydrocarbon compared with commercial reference standards (Sigma-Aldrich Corporation). Other compound names are a tentative identification by comparison with published peak mass spectra from Wiley Registry of Mass Spectral data (7th edition), and retention indices from NIST Chemistry WebBook[4]. Some compounds have been previously found in *Heliconius hecale*[1,3] and *H. melpomene*[2].

[c] Only in claspers from both species

1. Estrada, C., Schulz, S., Yildizhan, S. & Gilbert, L. E. Sexual selection drives the evolution of antiaphrodisiac pheromones in butterflies. *Evolution (N. Y).* 65, (2011).

2. Schulz, S., Estrada, C., Yildizhan, S., Boppré, M. & Gilbert, L. An antiaphrodisiac in *Heliconius melpomene* butterflies. *J. Chem. Ecol.* 34, 82–93 (2008).

3. Schulz, S., Beccaloni, G., Nishida, R., Roisin, Y. & Mcneil, J. N. 2,5 dialkyltetrahydrofurans, common component of the cuticular lipids of Lepidoptera. *Zeitschrift fur Naturforsch.* 53c, 107–116 (1998).

4. NIST Chemistry WebBook, NIST Standard Reference Database Number 69. at <http://webbook.nist.gov>(2015)
